# SNPic: SNP Topic Modeling for Interpretable Clustering of Complex Phenotypes

**DOI:** 10.64898/2026.04.22.720106

**Authors:** Zhang Leyi, Christof Seiler, Doug Speed, Raphael Micheroli, Caroline Ospelt

**Affiliations:** Center of Experimental Rheumatology, Department of Rheumatology, University Hospital Zurich, University of Zurich, Zurich, Switzerland; Center for Quantitative Genetics and Genomics, Aarhus University, Aarhus, Denmark

**Author notes:** Corresponding author(s). E-mail(s); Contributing authors. These authors contributed equally to this work (Raphael Micheroli, Caroline Ospelt).

**Keywords:** Genome-Wide Association Studies, Probabilistic Topic Modeling, Interpretable Knowledge Graph, Systems Genetics, Precision Medicine

## Abstract

Genome-wide association studies (GWAS) have cataloged thousands of disease-associated variants, yet a central challenge remains: decoding the shared, pleiotropic architecture that links complex phenotypes. Existing approaches, including dimensionality reduction methods and regression genetic models, either lack interpretability or rely on external linkage disequilibrium (LD) reference panels, limiting their ability to recover coherent biological mechanisms.

Here we introduce the SNP topic model (SNPic), a generative probabilistic framework that reframes GWAS summary statistics as a structured corpus and models genetic architecture using principles from Natural Language Processing (NLP). By treating phenotypes as documents and genes or the whole corpus of traits as words, SNPic applies topic models, e.g. Latent Dirichlet Allocation (LDA), to infer latent “genetic topics”, representing interpretable, overlapping biological modules that jointly explain complex traits. This formulation enables simultaneous reconstruction of trait relationships and identification of their underlying molecular drivers.

SNPic integrates two complementary schemes: *Sumstat-as-word* for capturing global phenotypic structure and *Gene-as-word* for resolving mechanistic detail, within a unified modeling framework. To ensure robustness, we introduce a stability-optimized inference pipeline based on bootstrap resampling, allowing data-driven selection of topic number and filtering of stochastic signals. Across extensive simulations, SNPic consistently outperforms conventional dimensionality reduction methods in recovering latent structure under both linear and non-linear, highly overlapping genetic architectures.

Applied to integrated FinnGen and UK Biobank datasets, SNPic identifies reproducible genetic topics corresponding to distinct biological programs, including HLA-mediated immune processes and transporter-driven metabolic regulation, with strong tissue-specific support. The framework further generalizes across species, organizing complex traits in maize, *Arabidopsis thaliana*, and cattle into biologically coherent modules.

Together, these results establish SNPic as a scalable and interpretable framework that shifts GWAS analysis from association cataloging toward the construction of an interpretable knowledge graph representing the latent semantic architecture of the genome. By unifying statistical genetics with NLP, SNPic reframes GWAS analysis as a probabilistic language modeling task, enabling the systematic decoding of complex trait architectures and delivering a systemic graph of cross-phenotype relationships.

## 1 Introduction

Genome-wide association studies (GWAS) have identified thousands of disease-associated variants [1–3], yet analyzing traits individually limits our understanding of the shared genetic architecture underlying these phenotypes [4–7]. While integrative methods such as polygenic risk scores [8–10], genetic correlations [11, 12], cross-trait meta-analysis [13], and Mendelian Randomization [14, 15] have begun to address this gap, they primarily quantify macroscopic statistical overlaps or aggregate risk. Consequently, they often fall short of revealing the systemic biological mechanisms, such as specific causal pathways, latent molecular networks, or distinct pleiotropic etiologies, that underlie these observed correlations, highlighting the critical need for integrative systems genetics approaches [16, 17].

While widely used in genetics [18–22], traditional dimensionality reduction techniques face fundamental limitations in capturing highly complex trait architectures. Linear models like PCA oversimplify non-linear relationships [23, 24]. Consequently, large-scale linear applications, such as truncated SVD applied to the UK Biobank [25], often yield feature loadings with limited biological interpretability and restricted generative flexibility. On the other hand, non-linear manifold learning approaches like UMAP operate as a “black box” lacking transparent feature mapping [26]. By frequently distorting global geometry [27] and erasing local density [28], UMAP risks generating algorithmic artifacts rather than true biological structures. The real-world impact of these distortions is significant, for instance, some UMAP visualizations were recently criticized for artificially segregating continuous human genetic variation into misleading, discrete clusters [29]. Imposing rigid, discrete clustering boundaries on such flawed embeddings ultimately fails to capture the inherently overlapping and mixed etiologies of complex diseases [30, 31].

Beyond dimensionality reduction, statistical geneticists frequently utilize regression models (e.g., structural equation modeling [32]) to explore shared genetic architectures. While highly powerful for estimating genetic correlations, these approaches are fundamentally designed for continuous mathematical transformations and require external Linkage Disequilibrium (LD) reference panels. Consequently, their reliance on exact LD matching can restrict their scalability in large-scale cross-cohort integrations, where perfectly matched reference panels are often unavailable [33, 34]. Besides, recent studies have attempted to find systemic mechanisms from GWAS at the pathway level by linking variants to transcriptomic expressions based on pre-defined biological databases [35]. Because these database-driven pipelines rely on manually constructed biological “dictionaries”, they limit the discovery of novel, unannotated pleiotropic architectures that fall outside existing dictionaries. Furthermore, the statistical genetics and computational biology communities increasingly recognize the profound value of constructing comprehensive, interpretable knowledge graphs to systematically decode complex genetic architectures [36, 37].

However, the genetic architecture of complex traits closely mirrors the compositional structure of human language. In natural language, reading words in documents like “stock” infers a “finance” topic, while “Lakers” infers a “sport” topic. A single document containing both sets of words is naturally understood as a mixture of these underlying themes (e.g., the business of sports). Furthermore, higher word frequencies for certain groups of words increase the probability that a document will be categorized into the corresponding latent topic. By this exact same logic, specific sets of associated genes (analogous to words) regulate overlapping biological pathways (analogous to topics), which collectively drive complex phenotypes (analogous to documents). Taking gout as an example, observing genes such as “SLC2A9” strongly infers a “urate transport” topic [38], while genes such as “HLA-DQA1” infers an “autoimmune” topic [39, 40]. This word-topic representation effectively interprets both the distinct characteristics of gout and its genetic associations with autoimmune diseases. Such conceptual similarity reveals a powerful, yet largely unexplored analytical pattern. By conceptualizing the genome as a biological corpus rather than a matrix of effect sizes, we can directly leverage Natural Language Processing (NLP) [41, 42]. NLP is a branch of Artificial Intelligence that enables computers to understand, interpret, and generate human language [43]. NLP models, such as probabilistic topic models, are intrinsically designed to decode latent, mixed-membership structures from massive datasets without necessitating external reference panels, offering a more natural mathematical fit for the highly combinatorial nature of genetic mechanisms [31, 44].

A widely adopted topic model is Latent Dirichlet Allocation (LDA) [45, 46], a Bayesian generative model that mathematically formalizes this exact biological-linguistic correspondence. While biomedical statisticians have adapted LDA to model complex compositions in phylogenetic trees, gene expression, and microbiome profiles [47–50], recent efforts have expanded into individual-level clinical models, including treeLFA [51], age-dependent topic modeling [52], and MixEHR-SAGE [53]. These methods approach the mining of Electronic Health Records by effectively treating individual patients as documents, successfully uncovering complex clinical multimorbidities and patient-level phenotypes.

Here, we introduce the SNP Topic Model (SNPic), the first framework that directly applies NLP topic modeling to GWAS summary statistics. The core innovation of SNPic lies in a novel dual-scheme architecture that reframes genetic data through two complementary “document-word” analogies. First, to capture broad phenotypic relationships, the *Sumstat-as-word* scheme treats a focal phenotype as a *document* and the corpus of all other analyzed phenotypes as its *words*. Second, to decode detailed biological mechanisms, the *Geneas-word* scheme maps phenotypes (*documents*) directly to specific genes (*words*). Consequently, the inferred latent *topics* operate at two distinct resolutions: resolving macro-level disease clusters and identifying the specific groups of genes driving the phenotypes with latent mechanisms. Supported by a rigorous statistical pipeline to ensure reproducibility, SNPic transforms GWAS interpretation from merely cataloging isolated associations to decoding the genetic architecture of complex phenotypes as a composition of interpretable biological modules.

We demonstrate SNPic’s distinct advantages through comprehensive simulations and the analysis of diverse real-world datasets, accurately reconstructing complex, biologically meaningful pleiotropic architectures across human diseases, Maize, *Arabidopsis thaliana*, and Cattle. By translating isolated genetic signals into a cohesive genetic language, SNPic enables the systematic construction of an interpretable knowledge graph of trait relationships, providing intuitive, systems-level biological interpretations that were previously neglected by conventional methodologies.

## 2 Results

### 2.1 Framework Overview

SNPic is a probabilistic topic modeling approach to deconstruct the latent genetic architecture of complex traits. The workflow processes GWAS summary statistics through a pipeline with noise filtering, dual-scheme representation, and stability-optimized inference (Figure 1).

**Fig. 1:**
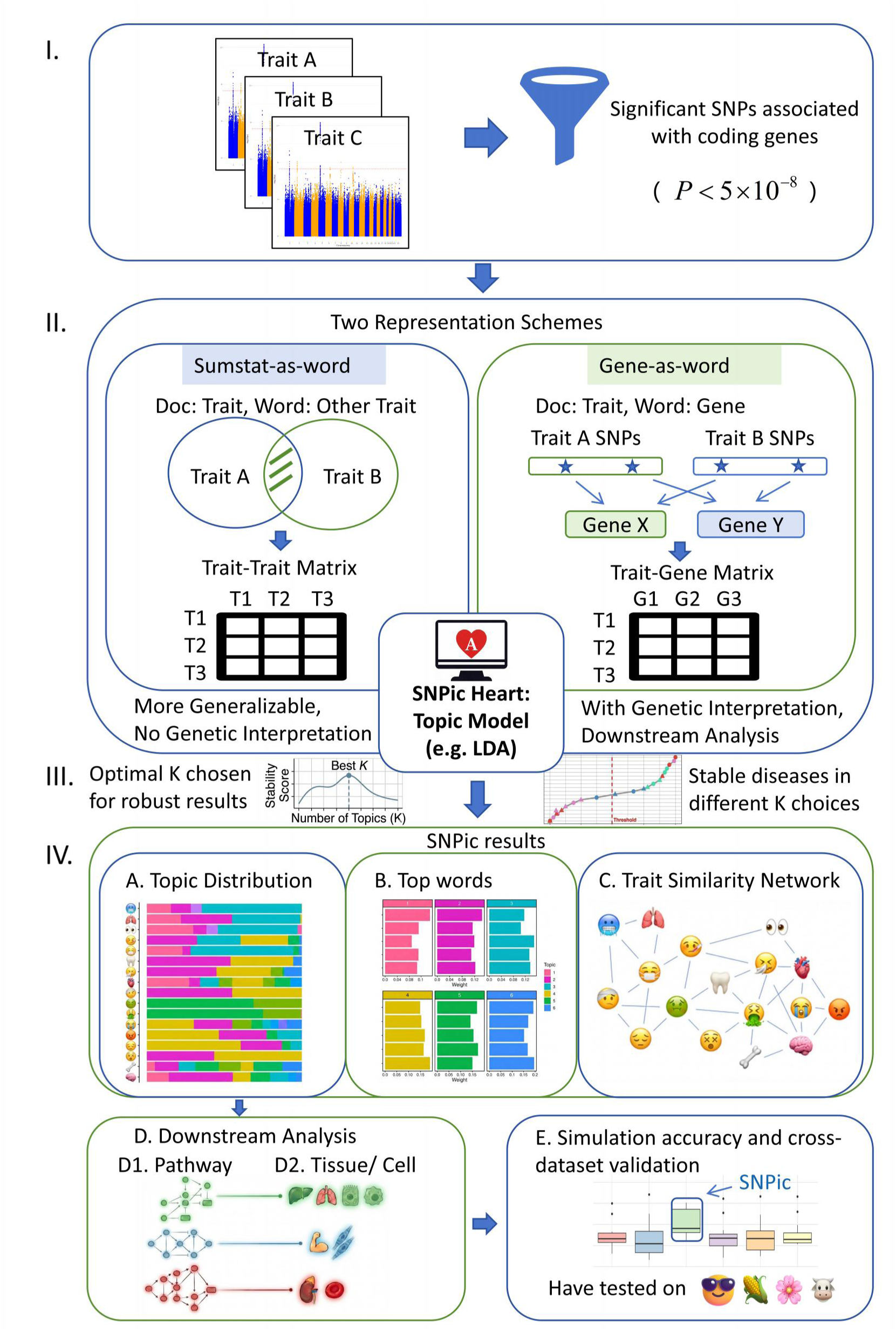
Overview of the SNPic Framework Workflow. The pipeline consists of four main stages: **(I) Data Input:** GWAS summary statistics are filtered for significant coding SNPs. **(II) Representation & Modeling:** Data is transformed into *Sumstat-as-word* and *Gene-as-word* matrices, which are then processed via Probabilistic Topic Modeling (LDA or Gaussian Mixed Membership Model). **(III) Stability Analysis:** A bootstrap procedure determines the optimal topic number (*K*) and filters out unstable traits. **(IV) Output & Validation:** SNPic generates disease-topic distributions, driver genes, and trait similarity networks to power downstream biological analyses (e.g., pathway and tissue enrichments) across diverse species. *Note: Visual elements are illustrative examples to demonstrate the workflow and do not represent actual study results*.

To ensure robust signal detection, we first standardized input data by filtering for significant SNPs using a strict and standard genome-wide significance threshold of *p <* 5 × 10^−8^ [54] (Figure 1 I). To avoid overrepresentation of large LD blocks, significant SNPs were clumped (e.g., *r*^2^ *<* 0.1 within 250kb) prior to intersection analysis. These filtered signals were then transformed into mathematical “documents” using two complementary representation schemes:

#### Summary-Statistics-as-word (Sumstat-as-word)

This scheme treats each trait as a document and the entire corpus of other traits as words, constructing a *Trait-Trait Matrix* (Figure 1, II left). Phenotypic co-occurrence is quantified as the number of shared significant SNPs intersection after LD clumping. This approach captures macroscopic phenotypic relationships and is particularly generalizable for species lacking detailed genomic annotations, as it relies solely on statistical overlap of SNPs.

#### Gene-as-word

This scheme maps significant SNPs to target genes (e.g., using the nearest-gene strategy directly from FinnGen summary statistics information) to construct a *Trait-Gene Matrix* (Figure 1, II right).

Here, traits act as documents and genes act as words. This formulation translates statistical associations into biological units, enabling downstream functional interpretations such as pathway enrichment and tissuespecific expression analysis.

An unbiased stability analysis is embedded in SNPic to replace arbitrary hyper-parameter selection. As illustrated in Figure 1 III, the model employs a bootstrap-based procedure to plot the Stability Score against the Number of Topics (*K*). This allows for the data-driven determination of the optimal *K* and the filtering of stochastic noise. Only associations with a Stability (Confidence) Score *>* 0.25 or top 50% stable traits are retained, ensuring that the resulting Trait Similarity Networks and Topic Distributions reflect persistent biological signals.

Upon establishing a stable model, SNPic outputs a suite of analytical results to decipher the genetic architecture (Figure 1, IV). The framework systematically generates the overarching topic distribution across all analyzed traits (Figure 1 IV-A) and identifies the top defining words (e.g., driver genes) that characterize each latent topic (Figure 1 IV-B). Furthermore, these probabilistic assignments are utilized to construct a Trait Similarity Network (Figure 1 IV-C), visually mapping the complex pleiotropic connections between distinct phenotypes. To transform these statistical topics into interpretable biological insights, SNPic integrates well-established downstream analyses. By focusing on the top 50 words per topic, SNPic performs targeted pathway enrichment analysis (Figure 1 IV-D1) and tissue-specific expression analysis (Figure 1 IV-D2) to elucidate the mechanisms of each theme. To assess the performance of SNPic, we validated its capabilities using controlled simulation data alongside public GWAS cohorts from plants (e.g., maize and *Arabidopsis thaliana*), animals (cattle), and large-scale human biobanks (Figure 1 IV-E).

### 2.2 Simulation Performance

To validate the mathematical accuracy of SNPic against ground truth, we conducted comprehensive simulations across three scenarios with progressively increasing structural complexity: (1) *Case 1: Linear with Four Groups* (Figure 2 (A-B)), (2) *Case 2: Linear with Six Groups* (Figure 2 (C-D)), and (3) *Case 3: Nonlinear with Periodic Assignment* (Figure 2 (E-F)). Each simulation compared SNPic against two established dimensionality reduction methods, PCA and UMAP, under both Normal and Gaussian data generation schemes. The clustering accuracy was quantified as the proportion of correctly reconstructed group memberships relative to the ground-truth simulated structure. Simulation rounds 1-10 progressively increased group overlap from 0% to 90% in 10% increments, with rounds 11 and 12 representing extreme overlap conditions of 95% and 99%, respectively. For concise visualization and evaluation, the graphical comparisons presented in Figure 2 and the numerical values discussed below represent the mean clustering accuracy averaged across all 12 of these simulation rounds for each scenario. Detailed performance metrics itemized for every individual round and overlap condition are available in Supplementary Table.

**Fig. 2:**
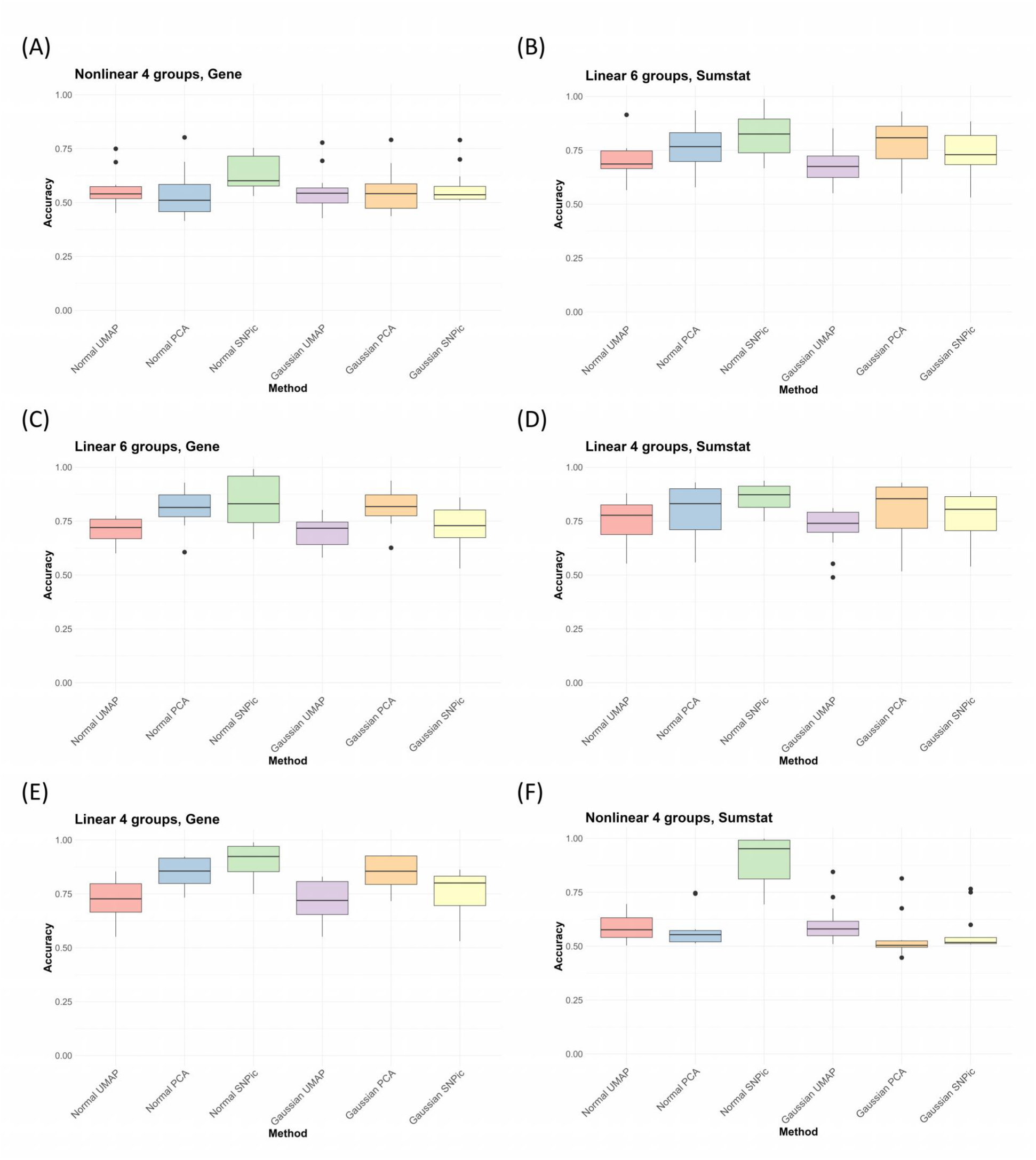
Performance comparison across all simulation scenarios. (**A-B**) **linear: 4 groups** demonstrates SNPic’s robustness in recovering basic genetic architecture under (A) Gene-as-word and (B) Sumstat-as-word scheme. (**C-D**) **linear: 6 groups** shows SNPic’s superior scaling properties with increased complexity under (C) Gene-as-word and (D) Sumstat-as-word scheme. (**E-F**) **non-linear periodic assignment** reveals SNPic’s substantial advantage in capturing complex non-linear relationships under (E) Gene-as-word and (F) Sumstat-as-word scheme.

Across all averaged scenarios, SNPic consistently demonstrated robust performance, achieving superior or comparable reconstruction accuracy relative to PCA while significantly outperforming UMAP in both linear and non-linear situations. The number of inferred topics (*N_topics_*) dynamically adapted between 4–10 across simulations, closely reflecting the true number of simulated groups and maintaining interpretability. In linear architectures (Figure 2 A-D), SNPic showed stable accuracy across increasing overlap conditions.

Under the *Gene-as-word* scheme, Normal SNPic achieved mean accuracies of 0.900 (4-group) and 0.841 (6- group), substantially outperforming both PCA and UMAP. The performance advantage was equally evident in Sumstat-as-word schemes, where Normal SNPic achieved the highest mean accuracy (0.860 for 4-group, 0.819 for 6-group) among all methods.

In the most challenging non-linear periodic assignment (Figure 2 E-F), SNPic demonstrated substantial advantages, particularly under the Sumstat-as-word scheme where Normal SNPic achieved a mean accuracy of 0.895, substantially surpassing PCA (0.577) and UMAP (0.591). This superior performance highlights SNPic’s capability as a generative probabilistic model to capture shared yet non-linear patterns of genetic contributions that are typically lost in algebraic embedding approaches.

### 2.3 Generalization across species

We then applied the Sumstat-as-word scheme to plant and animal data to evaluate SNPic’s generalizability across species and experimental conditions. Specifically, we wanted to test the model’s ability to mitigate batch effects in crops, recover functional modules in model organisms, and organize physiological traits such as livestock. Given the limited availability of comprehensive SNP-to-gene mapping resources in non-human species, we employed only the Sumstat-as-word approach for non-human validations.

#### 2.3.1 Maize (Zea mays)

We applied the *Sumstat-as-word* scheme to maize transcriptomics data collected from distinct experimental batches [55]. To validate the model’s biological relevance, we focused on key agronomic phenotypes including plant height, ear height, and kernel traits (Figure 3A). Utilizing the stability-driven selection strategy, we identified an optimal number of topics of *K* = 15 for the maize dataset (Figure 3B).

**Fig. 3:**
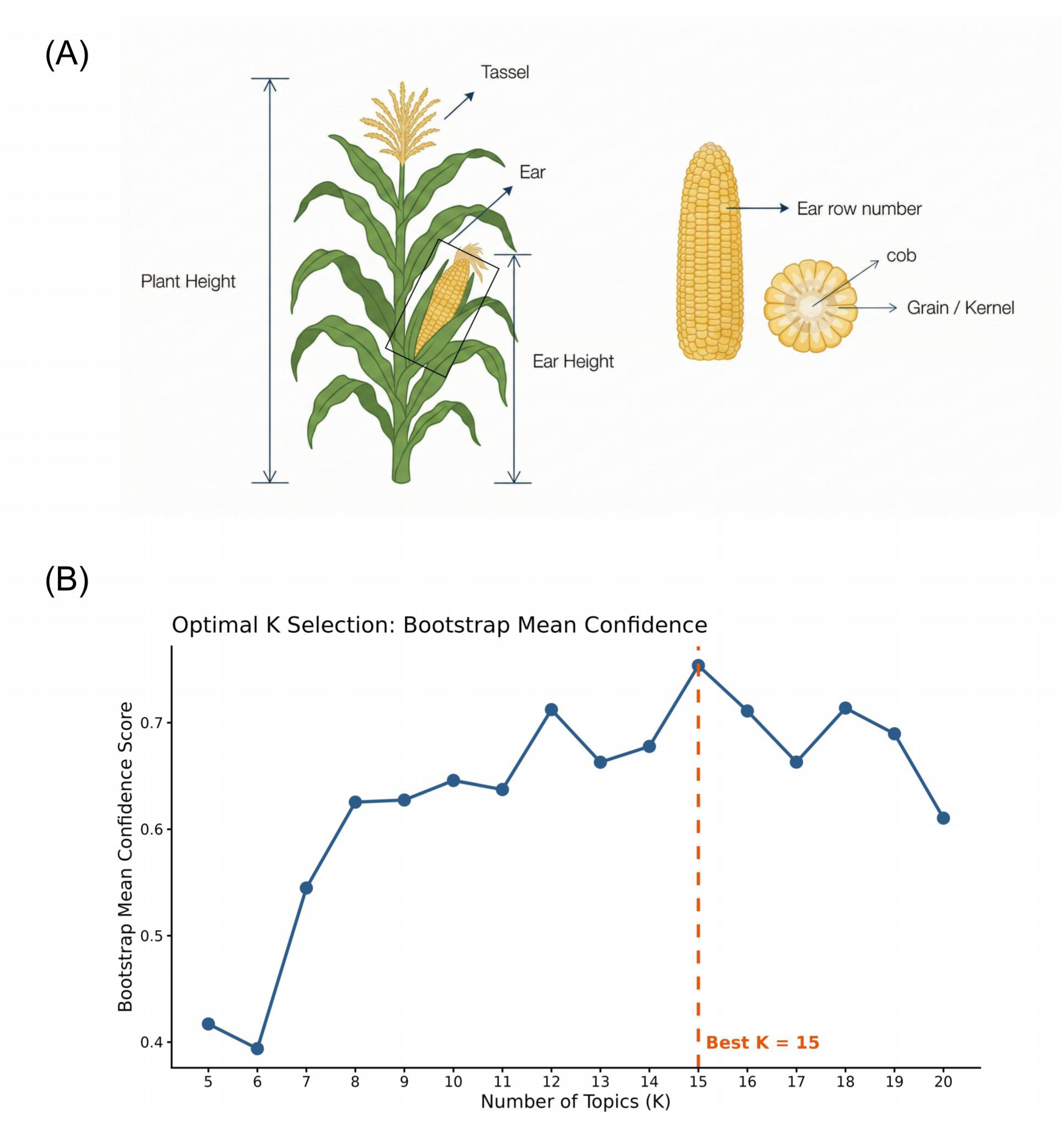
Maize Traits Illustration and Model Selection. (A) Maize Traits Illustration. The figure displays key phenotypic traits analyzed in maize (including Plant height, Ear height, and kernel traits). **(B) Optimal K Selection.** Bootstrap Mean Confidence score plot indicating the optimal number of topics (*K* = 15) driven by stability.

The detailed topic probability distributions and top-associated words revealed that SNPic effectively identifies consistent trait relationships across datasets (Figure 4A, B). For example, across different experimental batches, Ear Weight (represented interchangeably as EW and Ear weight) exhibits strong associations with Topics 6, 13, and 14, while Cob Weight (represented as CW and Cob weight) is primarily linked to Topics 6, 8, and 14. Their convergence on shared genetic components, specifically Topics 6 and 14, across disparate batches indicates that while batch effects inevitably introduce technical variation into the data, they do not overshadow the underlying biological architecture, and these inherent inter-group relationships between these biologically coupled traits are robustly preserved. Building upon this, the trait similarity network further demonstrates SNPic’s capability for co-analysis across multiple datasets (Figure 4C). Despite differences in genetic backgrounds and measurement protocols, the network demonstrates that the clustering is not completely dictated by batch effects. Instead, it successfully preserves broader biological associations. For instance, nodes representing “ear height” and “plant height” from different batches (indicated by distinct node shapes) not only formed tightly connected, trait-specific sub-networks (indicated by uniform node colors), but also maintained a clear inter-group topology linking these distinct traits together. This confirms SNPic’s capacity to extract robust, interconnected biological signals and retain true phenotypic relationships, even in the presence of technical batch variation.

**Fig. 4:**
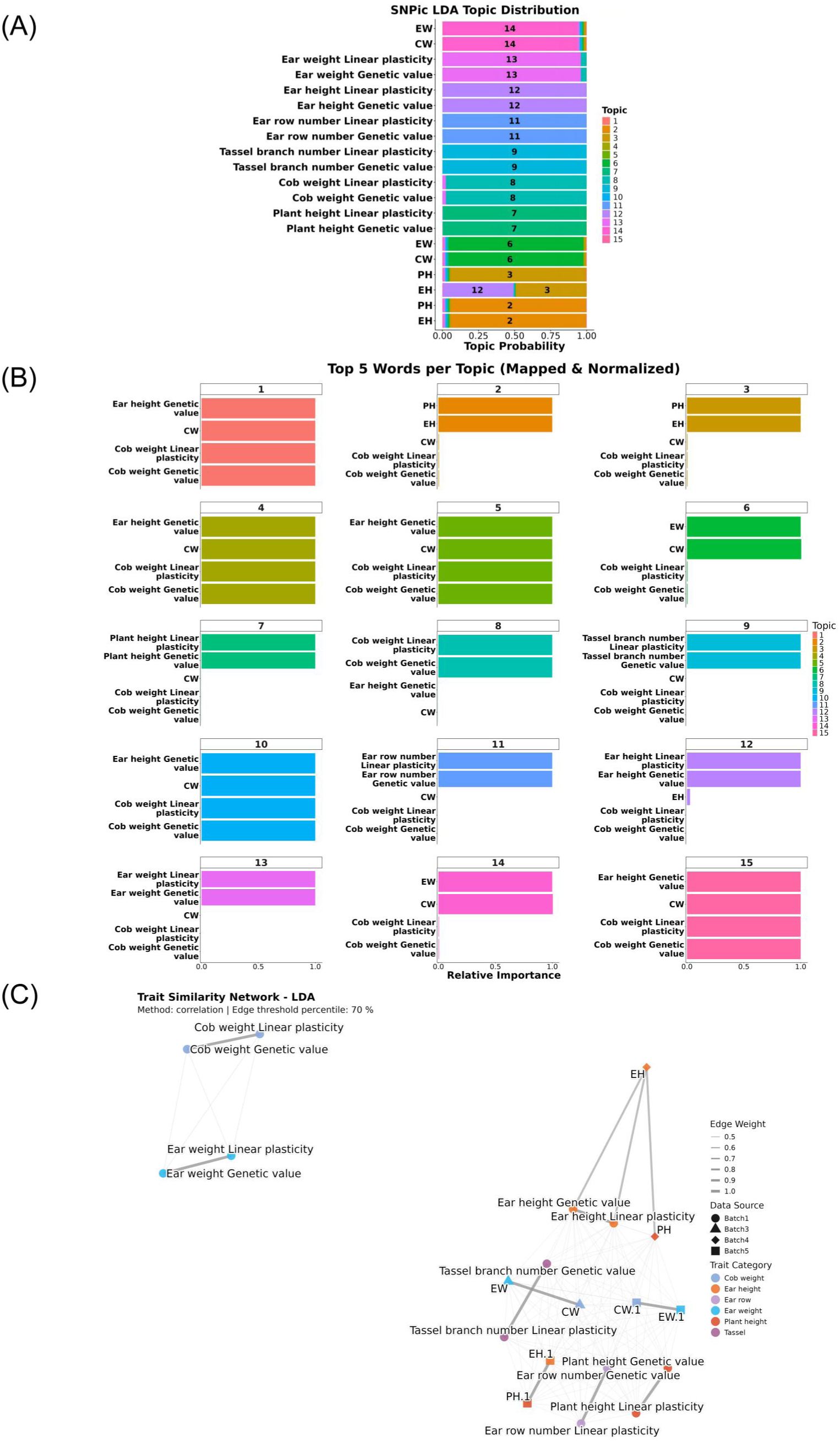
Topic distribution, feature importance, and network integration for maize agronomic traits. (A) Topic Distribution. Sorted barplot displaying the probability distributions of the 15 topics across different traits and batches. **(B) Top Words per Topic.** The top words driving the classification for each identified topic. **(C) Trait Similarity Network.** With a 70% edge threshold, the network integrates data across distinct batches to highlight inter-group relationships, mapping QTL clusters by agronomic trait (node color) and experimental batch (node shape). **Abbreviations:** CW: cob weight; EH: ear height; ER: ear row; EW: ear weight; PH: plant height.

#### 2.3.2 Arabidopsis thaliana

To assess biological precision, we applied the *Sumstat-as-word* approach to a classic GWAS cohort of *Arabidopsis thaliana* [56]. This dataset serves as a ground-truth benchmark as phenotypes have defined functional annotations. The optimal number of topics was identified as *K* = 5 based on the bootstrap mean confidence score (Supplementary Figure 1A). The resulting topic model demonstrated a robust capacity to cluster phenotypes according to their pre-defined functional categories, as visualized in the trait similarity network (Figure 5). The details of each trait are in Supplementary Table 1 and originally in the supplementary table of the paper [56].

**Fig. 5:**
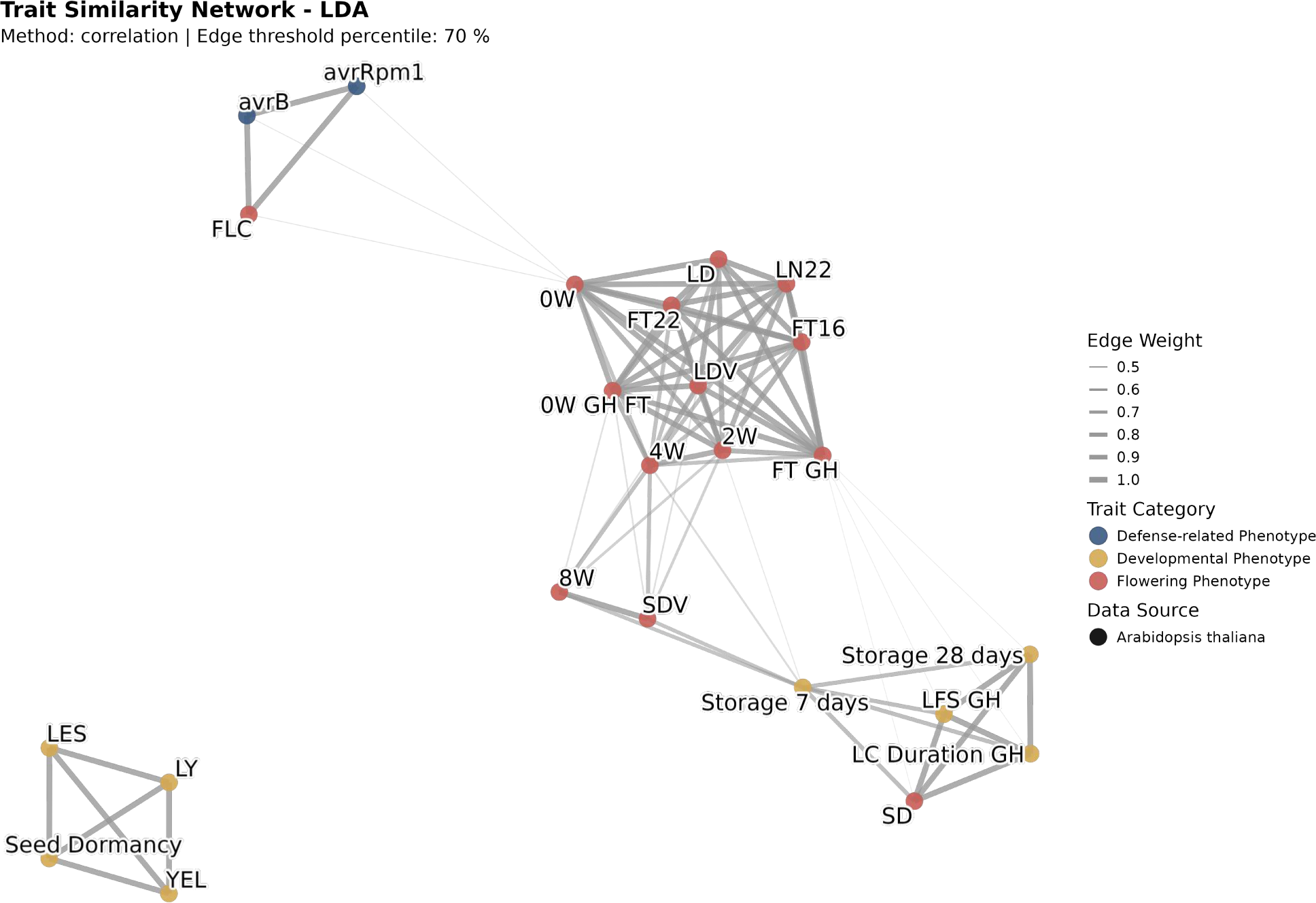
SNPic recovers biological structure in *Arabidopsis thaliana* GWAS data. The Trait Similarity Network (Edge threshold = 70%) clusters *Arabidopsis* phenotypes based on shared genetic architecture. Node colors represent broad functional categories as defined in the original study.

A particularly insightful finding was the recovery of a “function-interaction” link: the topic distribution (Supplementary Figure 1B) clearly reveals that FLC (Flowering Locus C), a central regulator of flowering time, is strongly associated with the same latent genetic topic (Topic 2) as the defense response phenotypes AvrRpm1 and avrB. While ostensibly distinct, this connection is biologically supported: polymorphisms in the *FRI* (Frigida) gene regulate *FLC* expression, and the same genomic region produces a multi-genic association with the *AvrRpm1* defense response [56–58]. This reveals SNPic’s ability to highlight pleiotropic connections that cross traditional functional boundaries.

#### 2.3.3 Cattle

We extended the analysis to livestock using the *Sumstat-as-word* framework on the cattle farmGTEx atlas [59]. The dataset comprised 43 economically important traits (details in Supplementary Table 2). The model identified an optimal configuration of *K* = 5 topics (Supplementary Figure 2A). The SNPic similarity network (Figure 6) organized these traits into biologically coherent groups matching manual annotations: Body Conformation, Health/Lifespan, Milk Production, and Reproduction. For instance, reproductive traits like “age at first calving” and “days to first breeding” formed a tight cluster distinct from, yet connected to, body conformation traits. The distinct topic assignments and driving top words for this structure are detailed in Supplementary Figure 2B and C.

**Fig. 6:**
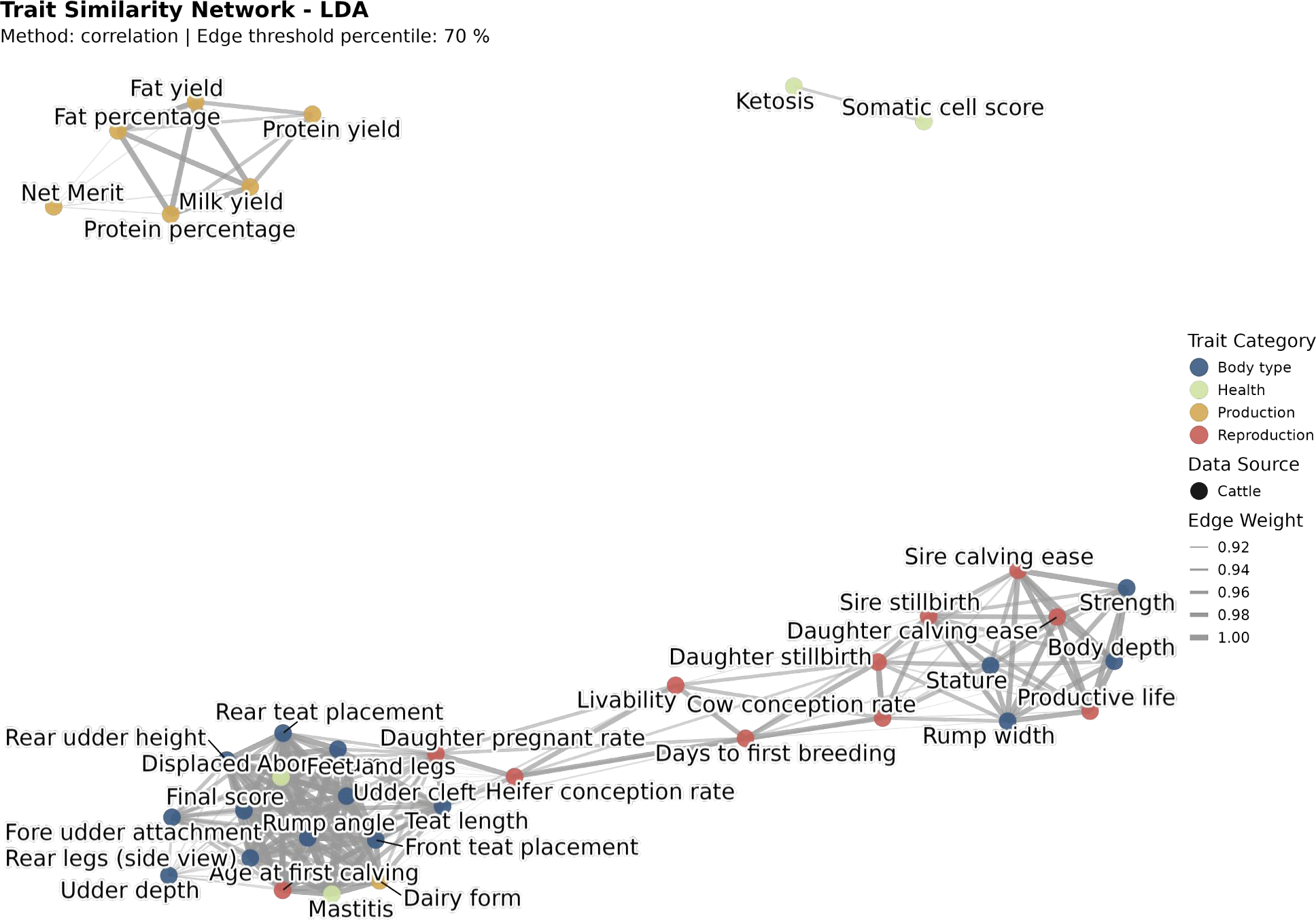
SNPic deciphers the genetic architecture of cattle complex traits. The Trait Similarity Network (Edge threshold = 70%) reveals the genetic relatedness among cattle production traits. Node colors represent annotated functional categories from the origin paper.

SNPic provided a robust and accessible framework for initial exploration of complex trait architecture in non-model organisms. Researchers can readily apply this method to available genomic datasets, with the option to incorporate *Gene-as-word* analyses when additional functional annotations become available.

The plant and animal genomics applications validated and highlighted SNPic’s utility as a generalizable tool for integrative genomics in diverse biological kingdoms from multiple experimental designs.

### 2.4 Human Cross-Biobank Integration

To construct a map of human disease architecture, we applied the SNPic framework to a combined dataset of FinnGen and UK Biobank (UKBB) summary statistics. We employed an unbiased, stability-driven pipeline to filter noise, validated the integration of cross-biobank data, and dissected the biological meaning of the inferred latent topics. The details of selected diseases are provided at Supplementary Table 3.

#### 2.4.1 Stability-Driven Model Selection

We first addressed the challenge of determining the optimal granularity of the genetic architecture. By evaluating the intra-*K* bootstrap volatility, we calculated the mean confidence score across a range of topic numbers. To avoid over-fragmentation and favor model parsimony, we implemented a strict tolerance-based selection strategy: we identified the global maximum stability score and established an acceptable tolerance threshold of 0.02 below this peak. The optimal model was defined as the smallest *K* whose stability score fell within this high-confidence range, leading to the definitive selection of *K* = 15 (Figure 7A). This strategy optimally balances model complexity with structural robustness for this corpus of complex traits.

**Fig. 7:**
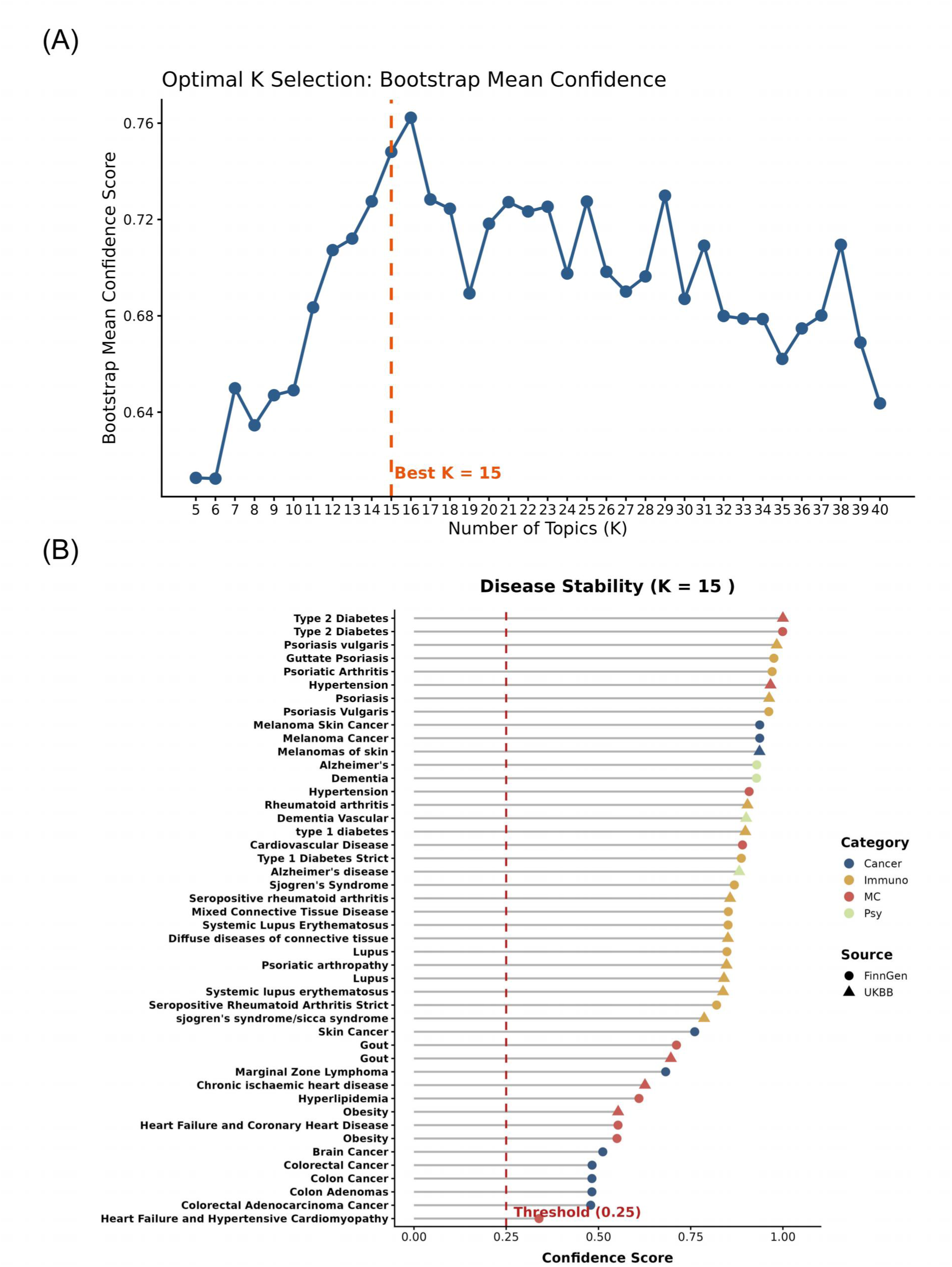
Stability optimization of the human complex traits. (**A**) Model stability curve across different numbers of topics (*K*). An optimal robust structure at *K* = 15 was identified using a parsimony strategy, selecting the smallest *K* within a 0.02 tolerance range of the global maximum confidence score. (**B**) Disease stability ranking (**Cancer**; **Immuno**: Immunology; **MC**: Metabolic/Cardio; **Psy**: Psychiatric/Neuro). A dynamic threshold (red dashed line at 0.25) filters out unstable phenotypes, ensuring high-confidence downstream analysis.

To further purify the results, we evaluated the stability of individual diseases within this optimal model. As shown in the ranking plot (Figure 7B), phenotypes exhibit varying degrees of coherence. We applied a dynamic thresholding strategy, setting the stability threshold to retain the top 50% of the diseases or achieve a Confidence Score *>* 0.25. This filtering step ensured that subsequent analyses were driven by persistent biological signals rather than stochastic noise. All evaluated traits in this human dataset achieved a Confidence Score well above this threshold, meaning no highly polygenic or environmentally sensitive traits were excluded, confirming that the inferred genetic structure is statistically robust and highly reproducible across disparate data sources.

#### 2.4.2 Interpretable Genetic Architecture

Focusing on the high-confidence disease set (*K* = 15), we deconstructed the latent genetic architecture using the learned topic distributions and corresponding gene vocabularies.

The *Topic Distribution* matrix (Figure 8A) visually captured the “mixed membership” etiology of complex traits. Systemic autoimmune diseases exhibited broad, overlapping profiles (e.g., lupus spanned Topics 1, 4, 5, 7, 8), reflecting their systemic multi-pathway nature. In contrast, traits like gout (dominated by Topic 2) or Alzheimer’s disease (dominated by Topic 6) displayed highly concentrated, specific topic assignments.

**Fig. 8:**
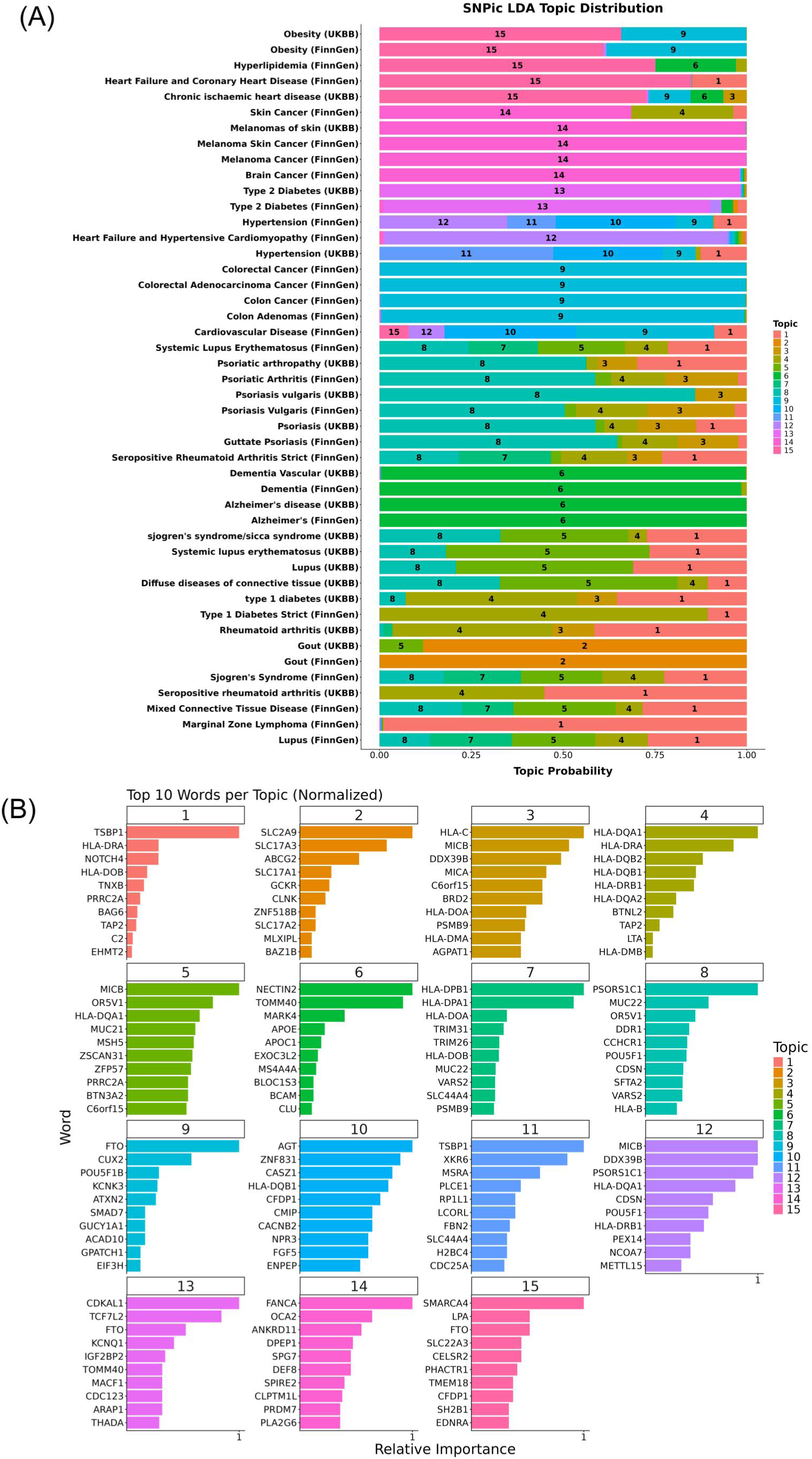
SNPic topic distribution and top genes. (**A**) Disease-topic distribution profile for stable diseases, illustrating the genetic topic architecture inferred by LDA (*K* = 15). (**B**) Top 10 genes with the highest normalized loading weights for each topic, defining the precise biological identity of the latent modules.

This architecture was biologically grounded by the *top genes* defining each topic (Figure 8B), acting as a transparent dictionary for the latent space.

For instance:

- **Topics 1, 3, 4, and 7** collectively structured the highly interconnected genetic basis of a broad spectrum of immune and autoimmune disorders. Topic 1 was primarily defined by *TSBP1*, *HLA-DRA*, *NOTCH4*, and *HLA-DQB1*, and it was dominant in marginal zone lymphoma (MZL) while contributing to systemic conditions like Sjogren’s syndrome and mixed connective tissue disease. Complementing this, Topic 3 captured MHC Class I genetics (*HLA-C*, *MICB*, *DDX39B*, *MICA*), whereas Topics 4 and 7 were enriched for MHC Class II alleles (*HLA-DQA1*, *HLA-DRA*, *HLA-DQB1*, *HLA-DPB1*). Together, this cluster formed the core architecture of systemic multi-pathway diseases, strongly driving the shared genetic risk for systemic lupus erythematosus, lupus, rheumatoid arthritis, and type 1 diabetes.
- **Topic 2** was exclusively defined by key urate transporters *SLC2A9*, *SLC17A3*, and *ABCG2*, aligning with its dominant probability loading in gout.
- **Topic 6** was defined by *NECTIN2*, *TOMM40*, *MARK4*, and *APOE*. While this topic dominated Alzheimer’s disease and dementia, it also showed significant contributions to hyperlipidemia, underscoring a critical genetic intersection where dysregulated lipid metabolism pathways potentially prime the landscape for neurodegeneration.

Comprehensive gene lists and analysis for all 15 topics are available in Supplementary Note.

#### 2.4.3 Similarity and Downstream Biological Analyses

##### Disease Similarity Network

Based on the inferred topic distributions, we generated the SNPic Similarity Network (Figure 9A). The global topology clearly deciphered major pathophysiological domains: cancer, autoimmune, metabolic, and psychiatric, while highlighting the pleiotropic connections among them.

**Fig. 9:**
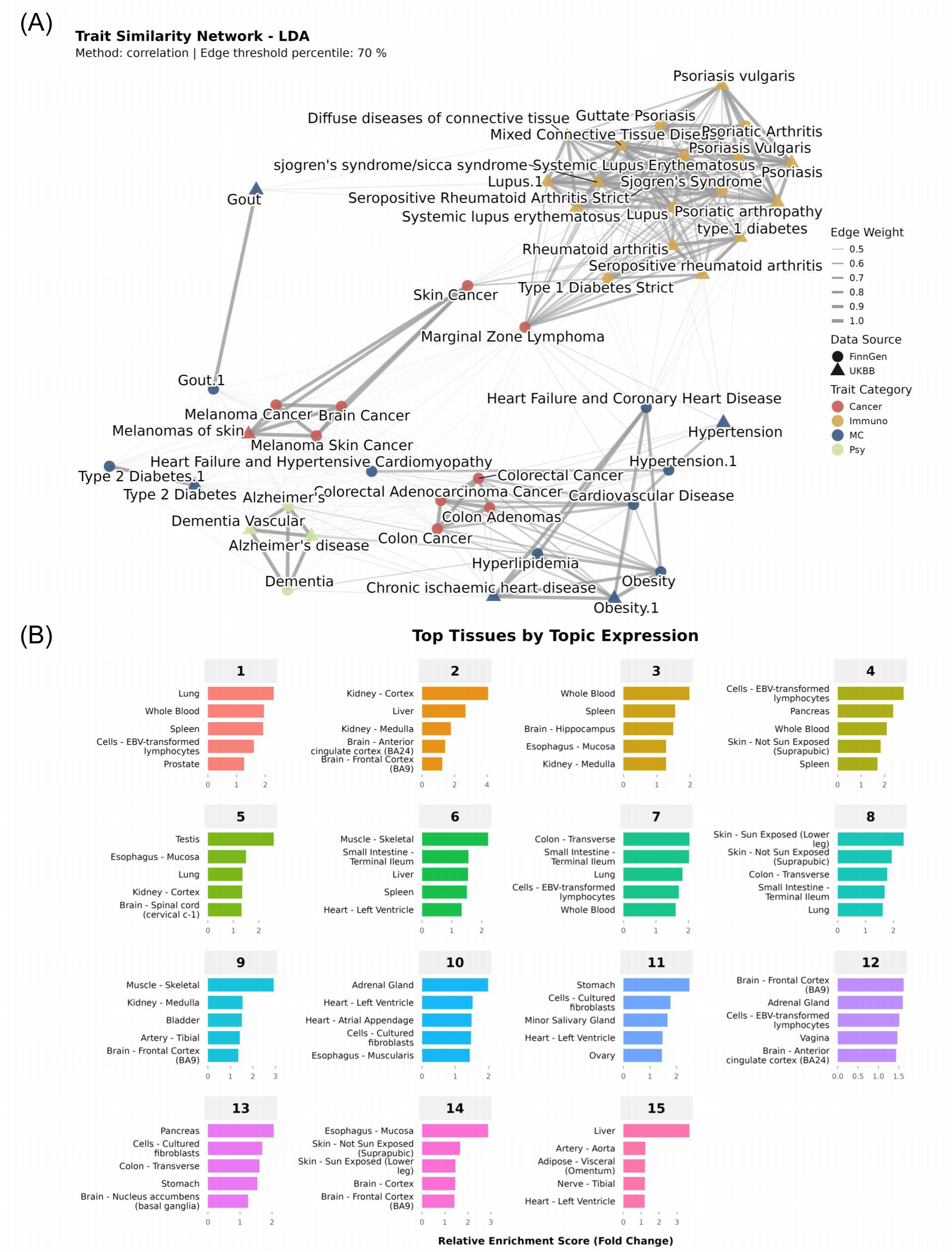
Network analysis and downstream biological validations. (A) Disease Similarity Network: The network visualizes the genetic relatedness across stable phenotypes based on their topic probability profiles. Nodes are colored by broad disease category (**Cancer**, **Immuno**: Immunology, **MC**: Metabolic/Cardio, **Psy**: Psychiatric/Neuro) and shaped by data source (FinnGen vs. UKBB). **(B) Tissue-Specific Expression:** A grid of horizontal bar charts displaying the relative enrichment score of the top genes for each of the 15 topics across their top five corresponding GTEx tissues.

The inferred network reproduced established clinical relationships across disease categories. The autoimmune cluster formed a dense module comprising rheumatoid arthritis, psoriasis, lupus, and type 1 diabetes, consistent with their shared HLA-driven genetic architecture. Within cancers, colon and colorectal cancers clustered together, whereas melanoma and brain cancers formed another cluster. The network also identified a connection between neurodegenerative traits (e.g., Alzheimer’s disease) and cardiovascular conditions (e.g., heart failure and hyperlipidemia), consistent with previously reported shared components involving lipid metabolism, inflammatory pathways, and vascular biology. In contrast, gout was allocated into an isolated module. Crucially, SNPic also indicated pleiotropic links crossing traditional clinical boundaries. For example, the model showed a genetic association connecting some cardiovascular diseases with the autoimmune disease cluster. This computationally derived genetic link is corroborated by epidemiological studies, showing a significantly increased incidence of cardiovascular diseases among aging patients with rheumatoid arthritis [60].

##### Functional Enrichment Analysis

To systematically validate the biological relevance of the inferred topics, we performed enrichment analyses using KEGG pathways and GO Biological Processes (Supplementary Table 4). The results indicated that SNPic successfully unmixed raw GWAS data into highly coherent functional modules:

Topic 2 was highly specific to “urate metabolic process” and “urate transport”, validating its exclusive association with gout.

Topics 1, 4, and 7 shared deep enrichments for “MHC protein complex assembly” and “antigen processing and presentation”, but differentiated into specific disease risks (e.g., Topic 7 was linked to “autoimmune thyroid disease” and Topic 4 to “asthma”). Furthermore, Topic 3 comprised specific cytotoxic and transplant-related immune activities, enriched for “cell killing” and “type I diabetes mellitus”.

Topic 6 showed distinct enrichment for “amyloid-beta clearance”, directly explaining its dominance in Alzheimer’s disease, while also linking to “cholesterol metabolism”.

Detailed enrichment profiles for all topics are provided in the Supplementary Material.

##### Tissue-Specific Expression

Finally, we examined the tissue-specific expression patterns of the top genes defining each topic to understand their anatomical context (Figure 9B). After normalizing the GTEx data by tissue to account for baseline global transcriptomic inflation, the analysis revealed striking tissue specificity that aligned these topics with their biological sites of action:

- Topics 1, 3, 4, and 7 showed robust expression in “whole blood”, “spleen”, “cells - EBV-transformed lymphocytes”, and “lung”, consistent with their roles in systemic autoimmunity, respiratory immune responses, and circulating immune cell functions.
- Topic 2 prominently featured the “kidney - cortex” and “liver”, matching the physiological sites for urate processing and excretion.

The complete tissue mappings of all 15 topics are provided in the Supplementary Material.

#### 2.4.4 Case Study: Reconstructing the Neuro-Cardiovascular Axis through Topic 6

We further explored the structural bridge observed within the *Disease Similarity Network* (Figure 9A and Figure 10A), which directly connects neurodegenerative diseases (e.g., Alzheimer’s disease, dementia) to specific cardio-metabolic conditions (e.g., hyperlipidemia, chronic ischaemic heart disease).

**Fig. 10:**
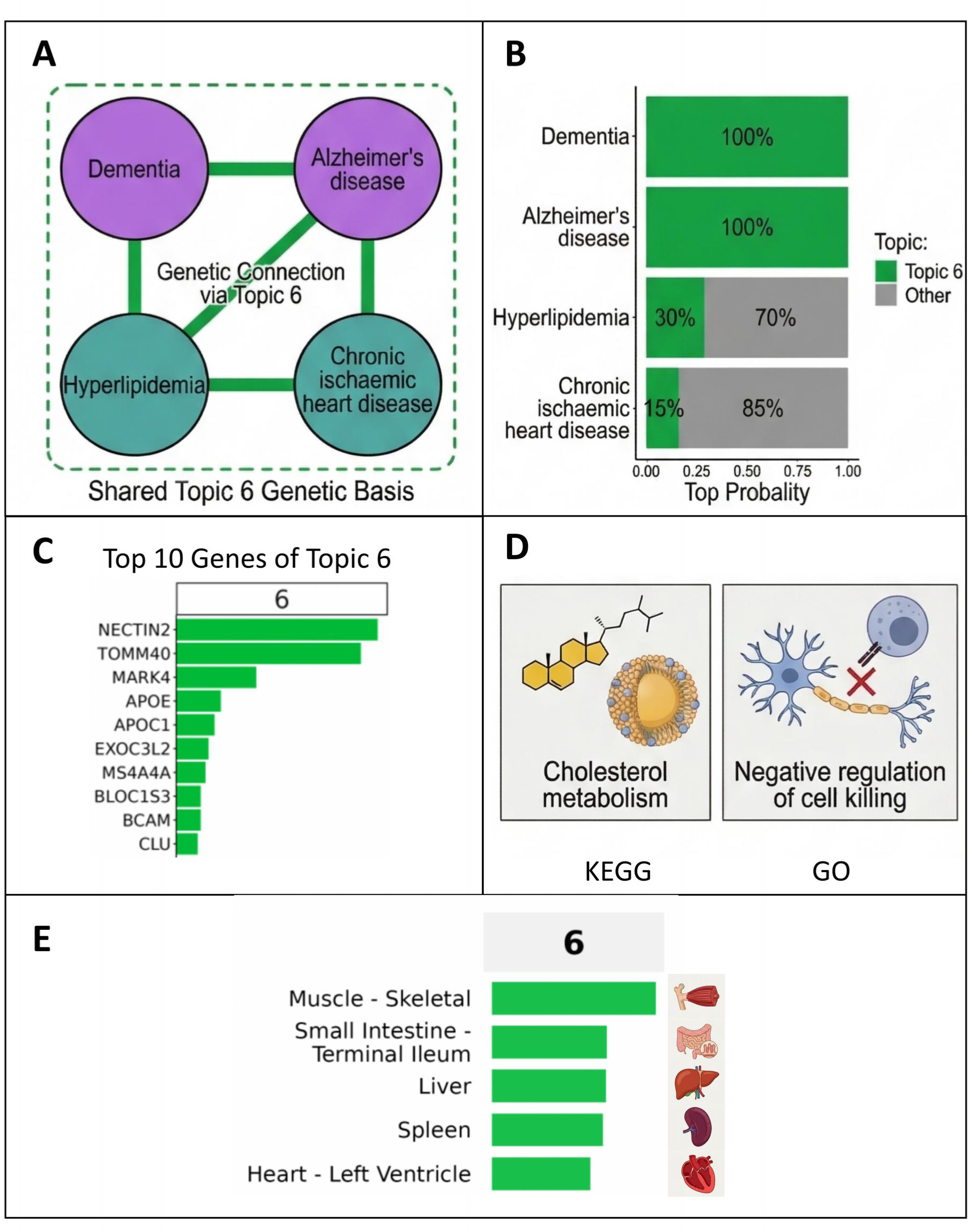
Reconstructing the neuro-cardiovascular axis via Topic 6. **(A)**: Simplified disease network demonstrating the genetic connection between neurodegenerative diseases (Dementia, Alzheimer’s disease) and cardio-metabolic conditions (Hyperlipidemia, Chronic ischaemic heart disease) via the shared genetic basis of Topic 6. **(B)**: LDA topic distribution of Topic 6 across the four diseases, showing Topic 6 accounts for 100% of the genetic architecture of Dementia and Alzheimer’s disease, while contributing some to Hyperlipidemia and Chronic ischaemic heart disease as well. **(C)**: Top 10 genes ranked by relative importance in Topic 6, including well-known disease risk genes, e.g., *NECTIN2*, *TOMM40*, and *MARK4*. **(D)**: Functional enrichment analysis of Topic 6, showing the core shared pathways: cholesterol metabolism (KEGG) and negative regulation of cell killing (GO). **(E)**: Tissue specificity score of the Topic 6 genetic module, with the highest enrichment in the Liver, followed by Spleen and the Small Intestine terminal ileum.

Examining the *Topic Distribution* (Figure 8A and Figure 10B) showed that the mathematical basis of this connection was based on Topic 6. While this topic almost entirely led the genetic architecture of Alzheimer’s and dementia, it reappeared as a substantial component in the genetic profiles of hyperlipidemia and chronic ischaemic heart disease.

To decode the biology of this “Neuro-Cardiovascular” axis, we analyzed the *Top Genes*, *Functional Enrichment*, and *Tissue-Specific Expression* profiles of topic 6. The top genes in topic 6 were *NECTIN2*, *TOMM40*, *MARK4*, *APOE*, *APOC1*, and *CLU* (Figure 8B Figure 10C). This topic integrated lipid biology pathway (KEGG: “Cholesterol metabolism”) with immune and vascular stress responses (GO: “negative regulation of cell killing”) (Figure 10D and Supplementary Table 4). However, tissue expression analysis showed that this genetic module localized primarily to metabolic and immune organs (e.g., small intestine, spleen, liver) rather than the central nervous system (Figure 9B and Figure 10E). This suggested a hypothesis that one component of this shared genetic risk might originate from peripheral lipid dysregulation.

To independently validate the computationally derived module, we projected the top 50 genes of Topic 6 into the STRING database [61]. The resulting protein-protein interaction (PPI) network formed a highly interconnected functional hub (Figure 11, Panel A), centered around core hub genes including apolipoproteins (*APOE, APOC1*), complement factors (*CLU*), and other top-ranked driver genes of Topic 6 such as *TOMM40* and *NECTIN2*, consistent with our gene importance ranking analysis. Complementing the PPI network, PubMed literature co-occurrence analysis further supported the functional and clinical relevance of this module. The gene set showed significant enrichment in published studies focused on Alzheimer’s disease, lipid metabolism, and neuro-cardiovascular comorbidity (Figure 11, Panel B) [62–66], with robust statistical significance supported by the low false discovery rate. The systemic genetic overlap captured by Topic 6 provides a molecular basis for emerging clinical models, which identify shared pathobiological mechanisms, such as lipid dysregulation, vascular inflammation, and ischemic damage, linking Alzheimer’s disease with ischemic stroke and cardiovascular events [62]. This shared lipid-centric etiology is independently corroborated by recent pharmacological evidence demonstrating that statins, first-line therapeutics for hyperlipidemia and cardiovascular disease, exhibit significant protective effects against the onset of Alzheimer’s disease [67]. Ultimately, this demonstrated SNPic’s potential to not only visually cluster diseases but to reveal the specific pleiotropic modules, offering direct translational insights for precision medicine and drug repurposing.

**Fig. 11:**
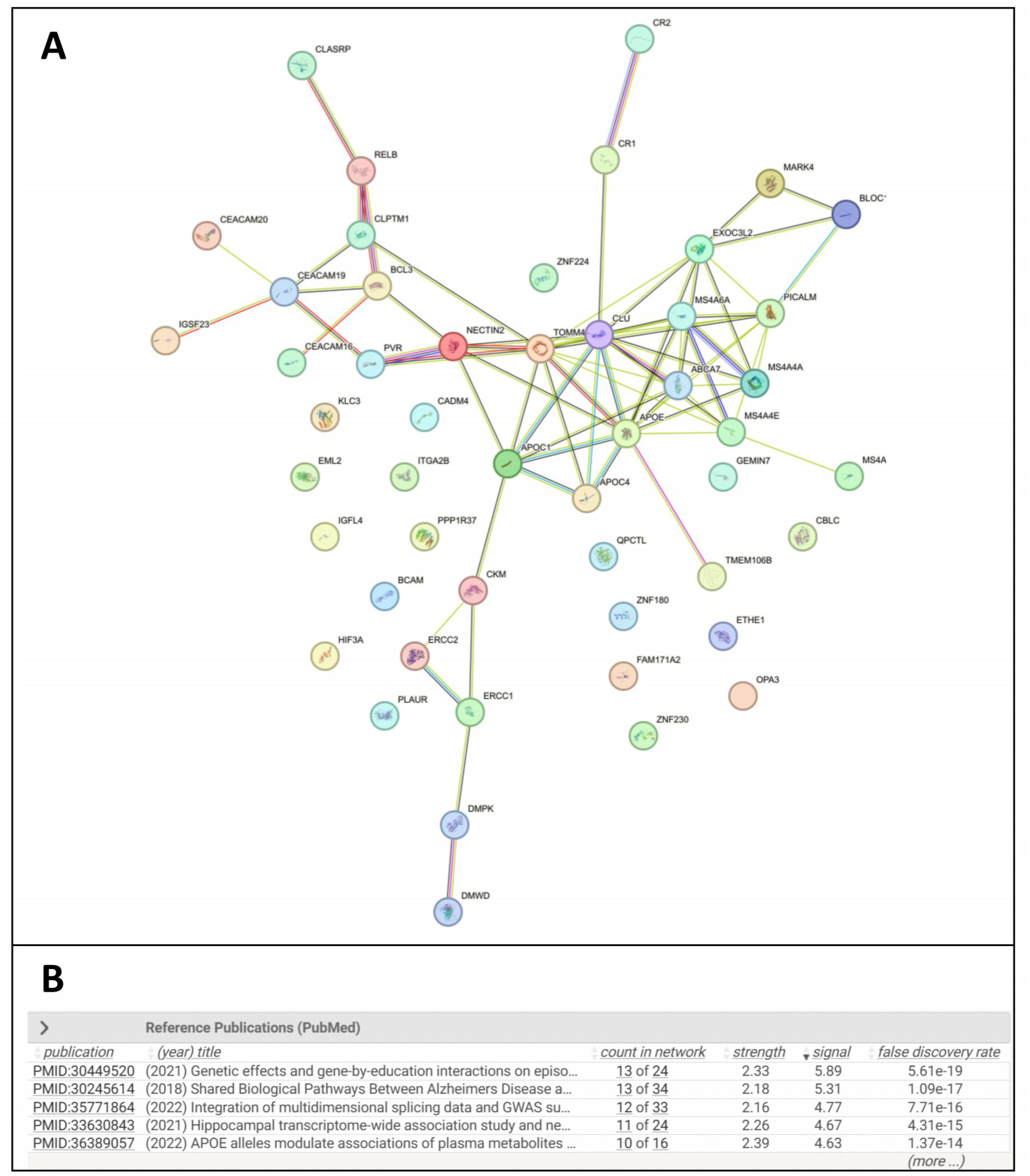
Independent validation of the Topic 6 genetic module via STRING database analysis. Panel. **A**: Protein-protein interaction (PPI) network constructed using the top 50 genes of Topic 6. Nodes represent individual genes, with node size corresponding to connectivity degree, and edges representing experimentally validated or database-curated functional protein interactions. The network forms a highly interconnected functional hub, with core hub genes including *APOE*, *APOC1*, *CLU*, *TOMM40*, and *NECTIN2*, the topranked driver genes of Topic 6 identified by our SNPic framework. **Panel B**: PubMed literature co-occurrence analysis of the gene set from the PPI network, ranked by false discovery rate (FDR). The table shows significant co-occurrence of the Topic 6 gene module in published studies related to Alzheimer’s disease, lipid metabolism, and neuro-cardiovascular comorbidity, providing independent literature-based validation of the module’s functional and clinical relevance.

## 3 Discussion

GWAS have cataloged thousands of trait-associated significant variants, yet a central challenge remains: moving beyond isolated signals to understand how discrete genetic factors group into coherent programs that shape complex phenotypes. Here we introduce SNPic, a generative framework that addresses this challenge through topic modeling, reframing genetic problems as NLP tasks. By analogizing phenotypes to documents and genetic features, e.g. genes, to words, SNPic infers latent probabilistic topics of genetics, which are interpretable modules that collectively explain the observed genetic architecture across traits.

Our simulations suggest that this generative framework offers a more direct and interpretable route to recovering latent structure in overlapping genetic architectures compared to descriptive correlation-based methods. While PCA and UMAP capture broad patterns of similarity, their performance is limited under the non-linear dependencies and high-dimensional overlap characteristic of polygenic architectures. In contrast, SNPic’s probabilistic model recovered the underlying latent structure in both overlapping and non-linear scenarios (Figure 2). This property aligns with the biological reality of pleiotropy, where a single variant often participates in multiple biological processes.

SNPic’s inference pipeline uses a data-driven, bootstrap-based procedure for hyper-parameter selection to identify a stable genetic architecture and determine the optimal number of topics from integrated FinnGen and UKBB data. By applying a dynamic thresholding strategy, that retains the top 50% of phenotypes or ensures a Stability Score *>* 0.25, our pipeline is designed to filter out noisy signals. Disease stability exhibited category-specific trends: immunological and cardiovascular disorders generally demonstrated higher stability, whereas highly polygenic or environmentally sensitive traits ranked lower. While conditions with diffuse genetic architectures (e.g., cancers) generally ranked lower in stability compared to the major-effect loci (e.g., the HLA region) driving many autoimmune diseases, all evaluated human traits maintained sufficient structural coherence to pass this dynamic threshold.

Critically, within this high-confidence set, the convergence of identical phenotypes from distinct biobanks (FinnGen and UKBB) into the same semantic clusters provides evidence that SNPic captures biological mechanisms rather than technical confounders from LD panels. Since LD panel structures and genotyping platforms differ between these cohorts, the observed consistency in topic distribution implies that the model is driven by shared functional architecture rather than localized correlation blocks. This demonstrates SNPic’s utility as a powerful tool for cross-cohort meta-analysis. While established frameworks like GenomicSEM provide powerful tools for modeling continuous genetic covariance, they are highly sensitive to mismatches in LD reference panels. In contrast, SNPic operates under a fundamentally different mathematical background: it acts as an LD-free, discrete generative model. This structural independence from external LD panels enables robust cross-biobank integration. For example, when integrating data from the UK Biobank and FinnGen, which are two populations with different demographic histories and unique LD architectures, SNPic uncovers shared biological mechanisms without the confounding bias that LD-dependent models would introduce.

Building on this robust cross-cohort integration, a foundational element enabling SNPic’s clear interpretability is its *Gene-as-word* mapping strategy. We directly adopted the nearest gene mapping provided by FinnGen, strictly utilizing coding-region genes to meet the requirements of pathway and tissue-specific analyses, to establish a clean baseline for the snp-to-gene strategy of SNPic. It is worth noting that the stringency of this baseline—requiring variants to reach genome-wide significance (*P <* 5 × 10^−8^) and map strictly within coding regions—inherently acts as a rigorous initial quality control. For instance, while we initially selected the UK Biobank cohorts for heart failure and non-Hodgkin’s lymphoma to serve as direct cross-biobank counterparts to FinnGen, they were naturally excluded prior to topic inference because they lacked any SNPs meeting these dual criteria. This strict prerequisite ensures that our downstream semantic structures are built exclusively upon highly confident, functional biological signals. To evaluate the impact of different SNP-to-gene mapping strategies, e.g. variant-to-function (V2F) mapping, we performed a sensitivity analysis by integrating a multi-tissue eQTL strategy using the complete GTEx v8 database. Both approaches yielded highly comparable results, effectively corroborating key findings, such as the shared genetic architecture between Alzheimer’s disease and hyperlipidemia (Supplementary Figures 17 - 21). However, compared to our baseline of nearest-gene strategy, the V2F integration introduced significant analytical noise. Aggregating pan-tissue V2F evidence unavoidably introduced an influx of false-positive regulatory links driven by off-target tissue noise, which artificially fractured the latent semantic structure, forced the optimal topic number (K) to jump to 25, and notably decreased overall computational efficiency. Currently, the definitive optimal SNP-to-gene mapping strategy remains an open and unresolved question in the field [68, 69]. Therefore, we provide the classical physical-distance-based nearest-gene strategy as an easily accessible, stable, and reliable baseline that effectively avoids these systematic multi-tissue biases. To maximize analytical flexibility, SNPic features a unified, modular interface allowing users to effortlessly supply custom SNP-to-gene mapping files. As future research continues to refine SNP-to-gene methodologies, new mapping schemes can be integrated into this interface. Consequently, future studies can leverage this open architecture to deeply investigate and determine the optimal mapping strategy specifically tailored for the SNPic framework.

The true value of this carefully mapped topic model lies in the interpretability of its components. SNPic’s genetic topics proved to be biologically coherent modules, each defined by a concise set of driver genes (Figure 8B) and linked to specific physiological systems. For example, autoimmune diseases decomposed into subtopics (e.g., Topics 4 and 7) implicating distinct HLA Class II alleles such as *HLA-DQA1* and antigen presentation pathways, recapitulating known immunological divisions [70–73]. Metabolic architectures were overwhelmingly defined by highly specific components such as Topic 2, driven by key urate transporters like *SLC2A9*, *SLC17A3*, and *ABCG2*, tightly linking gout with specific metabolite profiles. As a robust positive control for trans-systemic pleiotropy, SNPic successfully captured the link and latent mechanisms between MZL and the systemic autoimmune cluster [74–76].

The embedded tissue-specific expression analysis directly grounded these abstract genetic topics into human anatomy (Figure 9B). It is well-documented that tissues with pervasive baseline transcription, such as the testis, which broadly expresses the vast majority of the protein-coding genome, can disproportionately dominate standard downstream enrichments. To ensure our mappings accurately reflect true functional anatomy rather than non-specific background noise, we explicitly normalized the GTEx data by tissue. This rigorous step successfully mitigated global transcriptomic inflation, resulting in a highly specific anatomical mapping. Immune topics (e.g., Topics 4 and 7) localized robustly to hematopoietic and lymphoid tissues like the spleen and whole blood, while metabolic topics aligned with their precise physiological sites, such as Topic 2 (urate metabolism) in the kidney cortex and liver, and Topic 13 (type 2 diabetes) in the pancreas. Furthermore, this mapping clarifies cross-organ etiology, as it correctly assigns oncological components (e.g., Topic 14 for melanoma) to the skin, while revealing that the shared genetic component dominating Alzheimer’s disease (Topic 6) is primarily driven by peripheral metabolic tissues like the liver and small intestine rather than central nervous system tissues.

The generalizability of SNPic is strongly supported by its application across diverse species. It is important to note that, compared to the extensively curated and well-powered human GWAS datasets, the plant and animal data used in this study are less robust in terms of sample size, statistical power, and annotation completeness. To account for these inherent limitations and adopt a more exploratory, discovery-oriented approach, we adjusted our visualization strategy for these non-human datasets: we bypassed the stability filtering step. However, to ensure that the inferred pleiotropic connections remain highly reliable despite this methodological relaxation, we maintained a stringent correlation threshold of 70% for the network construction. In maize, it successfully clustered agronomic traits by biology rather than experimental batch. In *Arabidopsis thaliana*, SNPic recovered a known pleiotropic connection, grouping the flowering-time regulator FLC with the defense response phenotype AvrRpm1, a link supported by prior genetic studies [56–58]. In cattle, it autonomously organized complex production traits into a semantically sensible network matching expert annotation (Figure 6). This cross-species success suggests that the genome’s inherent modularity and reuse of functional units is a fundamental principle that SNPic is uniquely poised to reveal.

While powerful, the current framework has certain limitations. Its reliance on summary statistics, while highly scalable, precludes the modeling of haplotype structure or individual-level epistasis. Additionally, because topics are inferred from statistical associations, the model cannot definitively resolve causal relationships between distinct phenotypes (i.e., distinguishing where one disease directly causes another). However, drawing upon our core NLP analogy, just as a document is fundamentally constructed by its constituent words rather than the reverse, a complex trait is directionally driven by its underlying germline genes rather than the trait driving the genes. Consequently, while SNPic cannot ascertain trait-to-trait causality like Mendelian Randomization, the latent topics we identify are populated by genes that inherently act as directional biological drivers of the associated phenotypes. While our stability-optimized pipeline provides a robust, data-driven strategy to determine the optimal *K* and filter noisy phenotypes, we acknowledge that this selection relies on heuristic hyper-parameters (e.g., the 0.02 tolerance and the 0.25 dynamic threshold). While these boundaries inevitably retain a degree of arbitrariness, all evaluated human traits in our primary analysis consistently exceeded the 0.25 stability threshold. This confirms that our heuristic parameters did not artificially truncate or bias the final cross-cohort findings. To maximize transparency and support broader exploratory research, SNPic provides users the option to bypass this stability filter. For a comprehensive comparison against our main text’s optimal configuration, we provide alternative human data results including the Sumstat-as-word model, Gaussian LDA, unfiltered trait analysis, and FinnGen-only analysis in the Supplementary Materials (Supplementary Figures 4 - 16).

Unlike conventional statistical frameworks that rely on pairwise correlations or linear projections, topic models are able to recover latent mixed-membership structures from high-dimensional sparse data. This property makes them uniquely suited to model genetic pleiotropy, where variants simultaneously contribute to multiple biological processes in a context-dependent manner. Building on this capability, a primary innovation of SNPic lies in delivering a comprehensive, interpretable knowledge graph of the genetic architecture. This systems-level graph captures the complexity of biological functions and contextualizes genetic variants across multiple phenotypes, offering a macro-level perspective of biobank-scale genomics and micro-level interpretation of biological mechanism.

More broadly, SNPic represents an early step toward a conceptual unification of statistical genetics and Natural Language Processing. By treating GWAS summary statistics as a structured corpus rather than a collection of independent effect sizes, we demonstrate that the genome can be modeled using the same probabilistic principles that govern language. In this view, genetic variants function not merely as signals, but as elements of a combinatorial vocabulary whose meaning emerges only through context-dependent co-occurrence patterns. This reframing shifts GWAS analysis from correlation estimation toward semantic reconstruction, opening a new analytical paradigm in which biological systems are decoded as latent, interpretable compositions. Looking forward, SNPic provides a flexible foundation for numerous extensions. Different omic layers (e.g., transcriptomics, epigenomics, proteomics) can be naturally incorporated as parallel or hierarchical vocabularies, enabling a unified multi-modal topic model that captures cross-layer biological semantics. It could be adapted for longitudinal data to model the dynamic evolution of genetic contributions over time or disease stages. Most translationally, the disease-specific topic profiles (Figure 8) could serve as quantitative “genetic fingerprints” to stratify patient populations, predict comorbidity risk, or identify repurposable drug targets based on shared topic involvement rather than single shared genes.

In summary, we present SNPic, a probabilistic framework that redefines the analytical landscape of complex trait genetics. By conceptualizing genetic associations as a highly structured probabilistic language, SNPic deconstructs fragmented, biobank-scale GWAS catalogs into an interpretable lexicon of “genetic topics”. These inferred topics serve as fundamental, reusable biological modules that successfully dismantle rigid clinical boundaries, exposing the true pleiotropic spectrum of human pathology. Validated through rigorous mathematical simulations, stability-optimized inference on massive human cohorts, and generalization across diverse plant and animal species, our findings demonstrate that a generative, mixed-membership approach is essential for capturing the interconnected reality of the genome. Ultimately, SNPic shifts the field from cataloging isolated variants toward reconstructing an interpretable knowledge graph of the genome’s latent semantic architecture. By providing a highly scalable, privacy-preserving, and biologically transparent analytical lens, SNPic establishes a powerful new cornerstone for integrative genomics, paving the way for next-generation patient stratification and precision medicine. This work also builds a conceptual bridge between two traditionally separate fields: statistical genetics and NLP, suggesting that advances in one domain may directly transfer methodological innovations to the other.

## 4 Methods

### 4.1 The SNPic Framework

SNPic is a computational framework that adapts topic modeling to analyze GWAS summary statistics, transforming them into interpretable representations that reveal latent genetic structures. The core analogy treats complex traits as documents composed of genetic “words”.

#### 4.1.1 Data Representation Schemes

Formally, let a corpus of *p* diseases or phenotypes be represented as a set of documents *D* = {*d*_1_*, d*_2_*, …, d_p_*}. The words, denoted by *V*, consists of *N* distinct genetic features (“words”). SNPic implements two complementary schemes for defining this corpus:

1. **Summary Statistics-as-Word (Sumstat-as-word):** The word is the corpus of all genotyped or imputed SNPs, such that *V* = *S* = {*s*_1_*, s*_2_*, …, s_m_*}. A word *s_j_* is considered present in document *d_i_* if the SNP achieves genome-wide significance in the GWAS for disease *d_i_*.
2. **Gene-as-Word:** The word is defined as a set of genes, *V* = *G* = {*g*_1_*, g*_2_*, …, g_n_*}. Significant SNPs are mapped to their putative target genes via *SNP* → *G* mapping, with the current study employing the nearest-gene strategy, which can be user-adjustable. Some analysis proves the rationale in nearest-gene strategy [69]. This scheme enables direct biological interpretation through pathway enrichment and tissue expression analysis.

#### 4.1.2 Document-Term Matrix Construction

The input to the topic model is a *document-term* matrix 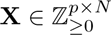.

For the *Gene-as-Word* scheme:

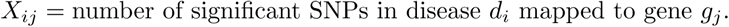

For the *Sumstat-as-word* scheme, *X_ij_* is the number of significant SNPs appeared in both diseases *d_i_* and *d_j_*.

#### 4.1.3 Model Decomposition into Latent Genetic Topics

SNPic employs a generative topic modeling approach where every disease is modeled as a mixture of multiple topics, and each topic is characterized by a distribution over genetic features. Given *N* diseases and *M* genetic features, SNPic infers *K* latent genetic topics through two probability distributions:

- **Disease-topic distribution Θ** ∈ R*^N^*^×*K*^: Each row ***θ****_i_* represents disease *d_i_* as a probability distribution over *K* topics.
- **Topic-word distribution Φ** ∈ R*^K^*^×*M*^ : Each row ***ϕ****_k_* represents topic *t_k_* as a probability distribution over *M* words.

The inferred latent topics correspond to coherent biological modules, enabling representation of any disease as an interpretable combination of fundamental genetic themes.

### 4.2 Topic Modeling with Latent Dirichlet Allocation

Under the discrete topic modeling formulation, each disease is treated as a document composed of multiple genetic tokens derived from GWAS summary statistics.

For disease *d_i_*, we construct a multiset of genetic features by expanding the document-term matrix *X*: each genetic feature *v_j_* (gene or SNP) is repeated *X_ij_* times, yielding a tokenized representation suitable for LDA.

For each disease *d_i_*, the topic proportions are drawn as:

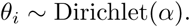

The generative process is defined as follows:

1. For each topic *k* ∈ {1*, …, K*}, draw a topic-specific distribution over genetic features:

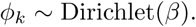
2. For each disease *d_i_*:
  (a) Draw topic proportions ***θ****_i_*.
  (b) For each genetic token *n* = 1*, …, N_i_*:
    (i) Draw a topic assignment *z_in_* ∼ Multinomial(***θ****_i_*).
    (ii) Draw a genetic feature *v_in_* ∼ Multinomial(*ϕ_zin_*).

### 4.3 Gaussian Mixed-Membership Model for Continuous Representations

While LDA assumes discrete token counts, GWAS-derived representations can also be naturally modeled as continuous-valued association strengths. To accommodate this setting, we additionally implement a Gaussian mixed-membership model as a continuous analogue of topic modeling.

In this formulation, each disease *d_i_* is represented by a continuous vector *X_i_*∈ R*^M^*, and is modeled as a convex combination of latent topics:

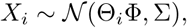

where Θ*_i_*denotes the disease-specific topic membership vector, Φ represents topic-specific genetic profiles, and Σ captures residual covariance not explained by topic structure.

This model relaxes the discrete multinomial assumption of LDA and enables direct modeling of standardized or continuous GWAS-derived features. The Gaussian and LDA formulations are analyzed in parallel to assess the robustness of inferred genetic architectures.

### 4.4 Stability-Optimized Topic Modeling

#### 4.4.1 Bootstrap Stability Analysis

Standard LDA inference is inherently stochastic, resulting in slight variations in topic distributions across runs due to the probabilistic nature of Gibbs sampling. To ensure robust topic inference and quantify the reliability of genetic relationships, we implemented a bootstrap-based stability analysis.

For each candidate topic number *K*, we generated 50 bootstrap replicates by resampling the genetic features within each trait. Specifically, for each trait document, a new feature count vector was drawn from a multinomial distribution parameterized by its original word frequencies, thereby maintaining the total significant variant count per trait while introducing stochastic perturbations. For each replicate, we performed LDA inference using a collapsed Gibbs sampler (burn-in=500 and iterations=1000) to estimate the trait-topic distribution matrix **Θ**. We then calculated the pairwise trait-trait similarity matrix using the Pearson correlation coefficient of their topic proportion vectors, scaling the coefficients to a [0, 1] range. These replicate-specific similarity matrices were used to quantify the estimation uncertainty across stochastic perturbations, forming the basis for our subsequent model selection and noise filtering procedures.

#### 4.4.2 Optimal Topic Number Selection

Selecting the optimal number of topics (*K*) is critical for balancing model resolution with structural robustness. Crucially, our evaluation framework assesses stability strictly through intra-*K* comparisons—measuring the variance of trait-trait similarities across bootstrap replicates within the same *K*—rather than attempting to align or compare topics across different *K* values. This design deliberately avoids the artifactual “bell-shaped” bias commonly observed in inter-*K* alignment metrics (where intermediate *K* values are systematically favored due to combinatorial matching constraints), thereby ensuring a fair and unbiased assessment across the entire tested spectrum.

For each candidate *K*, we quantified the stability of the learned structure using a “Confidence Score”. First, we defined the *Volatility* of a specific trait *t_i_* as the average standard deviation of its similarity scores with all other traits across the intra-*K* bootstrap iterations:

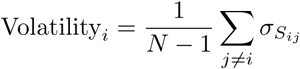

where *σ_Sij_* is the standard deviation of the similarity between trait *i* and *j* across the bootstraps for that specific *K*, and *N* is the total number of traits.

We then normalized this metric to derive a Confidence Score for each trait, where higher values indicate greater stability:

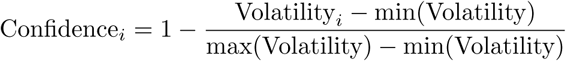

To determine the optimal *K*, we implemented a parsimony-driven selection strategy grounded in Occam’s Razor, prioritizing model simplicity while preserving near-maximal structural stability. We first calculated the mean Confidence Score across all traits for each candidate *K* to identify the global maximum stability (*C*_max_). To avoid the over-fragmentation of coherent biological pathways into artifactual narrow topics, we established a strict performance tolerance range defined as *C*_max_ − 0.02. The optimal *K* was then selected as the minimum *K* value whose mean Confidence Score fell within this pre-specified range.

While the one standard error (1SE) rule is a well-established heuristic for parsimonious model selection in regularized regression [77], we opted for this fixed 0.02 tolerance strategy for both computational and theoretical reasons. Rigorously implementing the 1SE rule requires nested cross-validation, which would impose a prohibitive computational burden on top of our existing full LDA Gibbs sampling and bootstrap replicates. Furthermore, as noted in prior methodological work [77], the 1SE rule itself remains an arbitrary threshold that can inadvertently increase parameter estimation bias. Given that optimal *K* selection in latent Dirichlet allocation broadly remains an open statistical challenge, we adopted a pragmatic approach. By calibrating our tolerance to 0.02 on a normalized [0, 1] scale, we tolerate at most a 2% relative reduction in stability performance. This margin is biologically negligible, effectively bypassing the computational bottleneck of the 1SE rule while successfully identifying the simplest model that retains highly replicable genetic associations.

#### 4.4.3 Stable Trait Filtering

Low-heritability traits or those with ambiguous genetic architectures often fail to cluster consistently. To mitigate noise, we applied a dynamic thresholding strategy to filter out unstable traits. For the selected optimal *K*, we established a dynamic stability threshold defined as the minimum between 0.25 and the median Confidence Score across all phenotypes.

Traits were retained for downstream network construction and biological interpretation only if their Confidence Score met or exceeded this dynamic threshold. This mathematical formulation guarantees that at least 50% of the analyzed phenotypes are retained for robust network construction, while capping the threshold’s strictness at 0.25. This quality control step ensures that the reported “genetic topics” and trait clusters reflect persistent biological signals rather than stochastic artifacts of the modeling process. However, to support broader exploratory analyses—particularly in datasets with lower statistical power or sparse annotations (e.g., non-model organisms)—the SNPic framework also provides a flexible option to bypass this filtering step, preserving the complete phenotypic landscape for downstream network construction.

### 4.5 Simulation Study

To rigorously evaluate the performance of SNPic in recovering latent genetic structures, we conducted a series of controlled simulations under varying levels of genetic overlap and structural complexity. We utilized LDAK (Version 6.1) to generate synthetic genotype data for 100,000 individuals across 500,000 SNPs [78]. Phenotypes were simulated with a high heritability (*h*^2^ = 0.8) and a prevalence of 0.1 to ensure robust genetic signals.

#### 4.5.1 Simulation Framework

The simulation pipeline followed a systematic six-step procedure to mirror real-world GWAS analysis:

1. **Causal Variant Definition:** We defined subsets of causal SNPs with precisely controlled pairwise overlaps to simulate shared genetic architectures.
2. **Phenotype Generation:** Phenotypes were generated based on the additive effects of the selected causal variants.
3. **GWAS Analysis:** Standard GWAS were performed on the synthetic cohorts to obtain summary statistics.
4. **Signal Extraction:** Significant associations were filtered using a threshold of *p <* 10^−4^ to retain informative genetic signals.
5. **Gene Mapping:** Significant SNPs were mapped to gene features to construct a *Gene-as-word* representation.
6. **Matrix Construction:** Finally, document-term matrices were constructed for input into the topic modeling framework.

#### 4.5.2 Simulation Schemes

We designed three distinct scenarios to test the model’s robustness against different architectural topologies:

- **Case 1 (Linear: Four Groups):** A baseline scenario where SNPs were distributed across four groups with linearly increasing pairwise overlap, ranging from 0% (distinct etiologies) to 99% (near-complete pleiotropy).
- **Case 2 (Linear: Six Groups):** An expanded linear model with six overlapping groups to assess

scalability and performance in higher-dimensional latent spaces.

- **Case 3 (Non-linear: Periodic Assignment):** A complex scenario where SNP group assignments followed a non-linear periodic function defined by val = sin(*G/*50), where *G* represents the gene index. This case was designed to evaluate the model’s ability to disentangle non-linear genetic relationships.

#### 4.5.3 Model Evaluation and Benchmarking

We benchmarked the performance of SNPic against two widely used dimensionality reduction techniques: PCA and UMAP. For SNPic, we evaluated both the standard discrete LDA model (Normal SNPic) and a continuous Gaussian Mixed-Membership model (Gaussian SNPic). The optimal number of topics (*K*) was determined by sweeping through a range of *K* ∈ [4, 10] and selecting the model that maximized reconstruction accuracy.

##### Evaluation Metric

Clustering performance was quantified by comparing the inferred disease-disease similarity matrices against the ground-truth structure. The ground truth similarity was derived from the known overlap of causal SNPs. Accuracy was defined as:

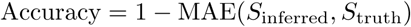

where MAE is the Mean Absolute Error between the inferred and true similarity matrices. This metric provides a rigorous measure of how well the inferred latent structure preserves the true underlying genetic relationships.

### 4.6 Data Resources and Preprocessing

To rigorously evaluate the robustness and broad applicability of the SNPic framework, we analyzed a highly diverse collection of real-world genomic datasets. By applying our methodology across massive human clinical cohorts, agricultural crops, and livestock populations, we aimed to demonstrate the model’s generalizability across different biological scales and experimental designs.

#### 4.6.1 Human GWAS Datasets

We integrated summary statistics from two large-scale biobanks:

1. FinnGen Consortium (Release 12): Comprising *>*500,000 samples [79]. We selected phenotypes spanning rheumatic, psychiatric, cardiovascular, and neoplastic categories.
2. UKBB: Summary statistics from the Pan-UKBB project [80, 81] were used for cross-dataset validation, specifically matching rheumatic phenotypes from FinnGen.

All data were lifted over to the GRCh37 assembly to accommodate the UKBB data version. To ensure robust signal detection across these real-world datasets, we filtered for significant SNPs using a strict genome-wide significance threshold of *p <* 5 × 10^−8^. As a pragmatic exception to retain informative genetic signals for under-powered traits, marginal zone lymphoma was filtered at a suggestive significance threshold of *p <* 5 × 10^−6^, since no SNPs for this trait passed the standard strict threshold but the suggestive threshold still captures biologically relevant variations [82]. Following this, significant SNPs were clumped (*r*^2^ *<* 0.1 within 250kb) to account for linkage disequilibrium. For cross-dataset validation, analysis was restricted to the intersection of SNPs matched in both cohorts.

#### 4.6.2 Non-coding Gene Filtering

To establish a clean and biologically interpretable “golden baseline” for the *Gene-as-word* scheme, we deliberately restricted our words strictly to protein-coding genes. It is worth noting that while the core SNPic framework does not strictly mandate the exclusion of non-coding genes, this filter was applied primarily because the downstream pathway and tissue enrichment modules are currently not equipped to process non-coding data. This conservative strategy isolates the intrinsic decomposition capability of the generative model while minimizing potential false-positive noise often introduced by complex variant-to-function (V2F) mapping algorithms in non-coding regions [69].

To systematically implement this filter, we constructed a comprehensive dictionary of valid protein-coding genes using the authoritative HUGO Gene Nomenclature Committee (HGNC) database [83]. We retrieved the complete HGNC dataset and filtered specifically for entities classified under the locus group of “protein-coding gene”. To maximize matching accuracy and account for historical nomenclature updates across different GWAS cohorts, we compiled an exhaustive vocabulary encompassing official gene symbols, alongside their documented alias and previous symbols.

For the primary SNP-to-gene projection, we employed a fundamental nearest-gene strategy. The resulting mapped genes were then rigorously cross-referenced against our HGNC-derived dictionary. Any mapped genetic features falling into non-coding regions, pseudogenes, or uncharacterized RNA genes were explicitly pruned from the dataset. This rigorous curation ensured that the final document-term matrix inputted into the LDA model consisted exclusively of high-confidence, biologically translatable protein-coding elements. Furthermore, to provide a comparative perspective and evaluate the impact of functional mapping, we conducted a supplementary analysis substituting the nearest-gene approach with a comprehensive V2F strategy. For this, we utilized multi-tissue expression quantitative trait loci (eQTLs) derived from the GTEx v8 database [84] to map variants to their functional target genes. To ensure analytical consistency, these eQTL-derived target genes were subjected to the identical HGNC protein-coding filtration process before topic modeling.

#### 4.6.3 Cross-Species Datasets

To demonstrate the broader utility of SNPic, we incorporated multi-omics summary statistics from diverse agro and model species:

- **Plant Genomics:** We obtained the eQTL summary statistics for Maize (*Zea mays*) directly from Supplementary Table S6 of Lei et al. [55]. For *Arabidopsis thaliana*, the classic GWAS summary statistics originally published by Atwell et al. [56] were retrieved from the easyGWAS public database [85].
- **Animal Genomics:** Multi-tissue TWAS data for cattle, representing 43 economically important traits,

were sourced directly from Supplementary Table 11, and categorized from Supplementary Table 10 of the farmGTEx project published by Liu et al. [59].

Unlike the human GWAS data, which required stringent post-hoc LD clumping and genome-wide significance thresholding, we did not apply additional quality control filters to the cross-species datasets. Because these summary statistics were sourced directly from the curated supplementary materials of peer-reviewed studies, we relied on the rigorous, species-specific QC pipelines already implemented by the original authors. Furthermore, given the comparatively limited sample sizes, lower statistical power, and sparser genetic architectures typical of non-human studies, imposing aggressive human-centric filtering parameters would risk discarding valuable, biologically relevant signals. Therefore, all provided significant variants from these curated sources were directly utilized for the *Sumstat-as-word* modeling. Comprehensive metadata detailing the specific traits, sample sizes, and curated mappings for these non-human datasets are consolidated in the Supplementary Tables of this paper (e.g., Supplementary Table 1 and Supplementary Table 2).

### 4.7 Downstream Analysis for Topic Interpretation

#### 4.7.1 Pathway Enrichment Analysis

For each topic, we extracted top 50 genes ranked by topic-specific weights and performed enrichment analysis using:

1. **KEGG Pathways**: Using enrichKEGG from clusterProfiler [86, 87]
2. **Gene Ontology Biological Process**: Using enrichGO with org.Hs.eg.db [88, 89] Significance threshold was *p <* 0.05 with FDR correction [90].

#### 4.7.2 Tissue-Specific Expression Analysis

We developed a pipeline to analyze tissue-specific expression patterns of topic-associated genes using GTEx data (V8). The analysis involved:

1. **Gene Expression Retrieval**: For each topic’s top genes, we queried the GTEx API to obtain median gene expression across 54 tissues.
2. **Topic-Tissue Association**: For each topic *k*, we calculated mean expression across its constituent genes for each tissue *t*:

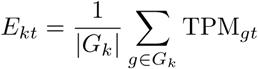

where *G_k_*is the set of top genes for topic *k*, and TPM*_gt_* is the expression of gene *g* in tissue *t*.

1. **Visualization**: To illustrate the anatomical localization of the genetic topics, we generated a grid of horizontal bar charts. For each topic, the top five tissues with the highest mean expression (TPM) are displayed, utilizing consistent color mapping to distinguish individual topics.

#### 4.7.3 Similarity Network Construction

Disease similarity networks were constructed based on the robust similarity matrix, *S*_robust_. An edge weight threshold of 0.70 was applied to retain only strong genetic correlations. In the resulting network visualizations, nodes were colored by disease category and shaped to distinguish the data source (FinnGen vs. UKBB).

#### 4.7.4 Topic Distribution Visualization

For the optimal model (selected *K*), we visualized the disease-topic probability distributions using sorted stacked bar plots. A consistent color mapping was maintained across all figures to ensure comparability between topic distributions, top gene profiles, and tissue enrichments.

### 4.8 Implementation and Availability

SNPic is implemented in R and available at https://github.com/KirakiraZLY/SNPic. The framework supports parallel computation for bootstrap analysis and includes functions for all visualization types described above. Additionally, Images A-D in Part IV of the graphical abstract (Figure 1), the maize illustration (Figure 3A), as well as the case study illustration of Alzheimer’s and Hyperlipidemia (Figure 10), were generated using Google’s Gemini 3 Flash Image (Nano Banana 2) model.

## Supporting information

This file contains the supplementary text, supplementary figures, and supplementary tables supporting the main manuscript.

## 5. Acknowledgements

We gratefully acknowledge Professor David Blei, a principal developer of Latent Dirichlet Allocation. His insightful seminar held at ETH Zurich in February 2025 provided the initial inspiration for this work and the foundational conceptual framework connecting Natural Language Processing to statistical genetics that underpins SNPic.

We gratefully acknowledge the participants and investigators of the FinnGen study and the UK Biobank study, whose dedication to advancing human health through large-scale biobank research provided the essential data for this study.

We also gratefully acknowledge Mengyu Lei and Qi Zhang, who generously shared and curated the plant and animal genetic datasets, which enabled the cross-species validation and comparative analyses presented in this study. Their contributions greatly enriched the scope and robustness of our model.

## Declarations

### Funding

This work was supported by an Investigator Initiated Study grant from Janssen-Cilag AG (pharmaceutical company of Johnson & Johnson).

### Competing interests

The authors declare that they have no competing interests.

### Ethics approval and consent to participate

Not applicable. This study exclusively utilized publicly available, de-identified GWAS summary statistics (e.g., FinnGen, UK Biobank) and public plant and animal datasets. No new human or other species subjects were recruited or experimented upon by the authors. Ethical approvals and informed consents for the original data collections can be found in their respective primary publications.

### Consent for publication

Not applicable.

### Data availability

The data that support the findings of this study are publicly available. Human GWAS summary statistics can be downloaded from the FinnGen project R12 website (https://www.finngen.fi/) and the Pan-UKB initiative (https://pan.ukbb.broadinstitute.org/). For the cross-species analyses, the original maize eQTL data were sourced from Supplementary Table S6 of Lei et al. (https://doi.org/10.1093/g3journal/jkaf140). The *Arabidopsis thaliana* GWAS data from Atwell et al. (https://doi.org/10.1038/nature08800) are hosted and publicly accessible via the easyGWAS database (https://easygwas.biochem.mpg.de/gwas/myhistory/public/3/). The cattle TWAS data were retrieved from Supplementary Table 11 of Liu et al. (https://doi.org/10.1038/s41588-022-01153-5). All specifically curated traits used in the cross-species analyses are comprehensively documented in the Supplementary Tables of this manuscript. Furthermore, pre-processed datasets, including the SNP-to-gene mapping files and demo matrices used in this study, are provided in the data folder of the project’s GitHub repository.

### Materials availability

Not applicable.

### Code availability

The SNPic framework is open-source. All original R scripts, shell scripts for data preprocessing, and detailed instructions required to reproduce the analyses presented in this study are freely available on GitHub at https://github.com/KirakiraZLY/SNPic.

### Author contribution

Z.L. conceptualized the project and designed the original methodology. C.S. and D.S. provided critical input on the statistical modeling and contributed to the design of the stability evaluation metrics. Z.L. developed the SNPic software, performed the formal analyses, and curated the datasets. Z.L., R.M., and C.O. contributed to the biological interpretation of the topics. Z.L. wrote the original draft. R.M. and C.O. acquired the funding, supervised the project, and critically reviewed the manuscript. All authors read and approved the final manuscript.

### AI Usage Declaration

During the preparation of this work, the authors used Large Language Models (LLMs) to improve the readability and language fluency of the manuscript. After using this tool, the authors reviewed and edited the content as needed and take full responsibility for the content of the publication. Additionally, Images A-D in Part IV of the graphical abstract (Figure 1), the maize illustration (Figure 3A), as well as the case study illustration of Alzheimer’s disease and Hyperlipidemia (Figure 10), were generated using Google’s Gemini 3.1 Flash Image model (formerly evaluated under the codename Nano Banana 2).

## Notes

https://github.com/KirakiraZLY/SNPic

### Competing Interest Statement

The authors have declared no competing interest.

