## Supplementary material for "SNPic: SNP Topic Modeling for Interpretable Clustering of Complex Phenotypes": This file contains the supplementary text, supplementary figures, and supplementary tables supporting the main manuscript.

#### 1 Supplementary Note: Case Study: Interpreting Gout through SNPic

To illustrate the interpretative power of SNPic, we trace the analysis of Gout across its multi-modal outputs. This case study demonstrates how the framework translates raw genetic data into clear biological insights through a structured, four-step sequence.

As shown in the *Topic Distribution* (Figure 8A), Gout is almost exclusively driven by a single genetic module: Topic 2. By examining the *Top Genes* for this topic (Figure 8B), we immediately identify its molecular drivers as well-known urate transporters, specifically *SLC2A9*, *SLC17A3*, and *ABCG2*.

The *Functional Enrichment* analysis (Supplementary Table 4) further supports this, showing that Topic 2 is statistically dominated by the "urate metabolic process" (GO), as well as "ECM-receptor interaction" and "Insulin resistance" (KEGG).

Because Gout possesses such a unique and specific genetic etiology, it appears as an isolated node in the *Disease Similarity Network* (Figure 9A), successfully separating itself from complex, multi-pathway clusters like autoimmune diseases.

Finally, the *Tissue-Specific Expression* analysis (Figure 9B) grounds this genetic signature in human anatomy, mapping Topic 2 directly to the "Kidney - Cortex". This seamless sequence, moving from an abstract probability distribution to specific transporter genes, through an isolated network position, and finally to the human kidney, demonstrates how SNPic provides a transparent and biologically intuitive roadmap for complex disease research.

#### 2 Supplementary Note: Case Study: The Autoimmune Axis in Marginal Zone Lymphoma

Another demonstration of SNPic's ability to transcend traditional clinical classifications is the positioning of Marginal Zone Lymphoma (MZL). Clinically categorized as a hematological malignancy, MZL defies expectations in the Disease Similarity Network (Main Text Figure 9A) by embedding itself deeply within the systemic autoimmune cluster, forming tight connections with Seropositive Rheumatoid Arthritis, Sjogren's Syndrome, and Systemic Lupus Erythematosus. Crucially, SNPic autonomously reconstructed this structural bridge between immunology and oncology entirely devoid of any prior clinical knowledge, utilizing solely raw GWAS summary statistics.

Deconstructing its genetic architecture via the Topic Distribution reveals that MZL shares an extensive genetic overlap with these autoimmune conditions, prominently driven by **Topic 1** (alongside components of Topics 4, 5, 7, and 8). Topic 1 is characterized by critical immune-regulatory and MHC Class II genes (e.g., *TSBP1*, *HLA-DRA*, *NOTCH4*, *HLA-DOB*). Furthermore, the tissue-specific expression analysis maps Topic 1 directly to "Whole Blood", "Spleen", and, most notably, "Cells - EBV-transformed lymphocytes".

By autonomously grouping MZL with autoimmune conditions rather than solid tumors (e.g., Melanoma, Brain Cancer), SNPic successfully captures the shared biological mechanism between systemic autoimmunity and B-cell tumors. This demonstrates SNPic's capacity to independently recover complex, cross-disciplinary mechanistic links directly from the underlying statistical architecture.

#### 3 Supplementary Note: Sumstat-as-word results of Human data

To complement the *Gene-as-word* biological interpretation presented in the main text, we additionally applied the *Sumstat-as-word* representation scheme to the integrated FinnGen and UK Biobank (UKBB) dataset. In this specific formulation, the document-term matrix is constructed using phenotypic traits as both the "documents" and the "words". The frequency of a "word" is quantified by the number of shared significant SNPs (after LD clumping) between a given pair of traits. This annotation-free approach is designed to capture macroscopic pleiotropic relationships and disease-disease co-occurrences directly from raw GWAS summary statistics, without relying on any external SNP-to-gene mapping algorithms.

**Stability-Driven Model Selection**

Following the same rigorous stability pipeline used in the main analysis, we evaluated the bootstrap stability across a range of topic numbers ($K$). Using our parsimony-driven tolerance strategy, we identified $K = 13$ as the optimal model complexity for the *Sumstat-as-word* representation (Supplementary Figure 11A). We then applied our dynamic thresholding strategy to evaluate individual disease stability within this $K = 13$ model (Supplementary Figure 11B). Consistent with the *Gene-as-word* results, identical conditions from independent biobanks (e.g., Psoriasis, Type 2 Diabetes, Alzheimer's disease) consistently ranked at the top, confirming that the foundational genetic overlap between these traits is highly robust regardless of the specific mathematical representation scheme employed.

**Topic Architecture and Trait-level Interpretation**

The detailed structural decomposition of this scheme is visualized in Supplementary Figure 12. Panel A displays the mixed-membership topic distribution for each phenotype. Crucially, in the *Sumstat-as-word* framework, the "Top Words" defining each topic (Panel B) are actual phenotypic traits rather than individual genes. This provides a direct, clinical-level interpretation of the latent modules. For instance, autoimmune-related topics are explicitly defined by the co-occurrence of traits like Rheumatoid Arthritis and Type 1 Diabetes, whereas metabolic topics are heavily weighted by Type 2 Diabetes and Obesity. This confirms that SNPic can reliably extract systemic clinical relationships and comorbidity structures purely from raw genetic overlap.

**Disease Similarity Network**

Finally, Supplementary Figure 13 illustrates the robust disease similarity network derived from the *Sumstat-as-word* topic proportions. Strikingly, the global topology mirrors the architectural findings from the *Gene-as-word* network. The network successfully reconstructs the major pathophysiological boundaries, forming tight, biologically coherent clusters for autoimmune disorders, cardio-metabolic syndromes, and neoplastic diseases. The consistency of these macroscopic structures across two fundamentally different word schemes strongly underscores the capability of the SNPic framework to extract genuine, foundational pleiotropic signals from large-scale GWAS cohorts.

#### Supplementary Note: Extended Topic Analysis for Human Cross-Biobank Integration

In addition to the primary topics discussed in the main text (Topics 1, 2, 3, 4, 6, and 7), the SNPic framework successfully delineated several other highly coherent functional modules governing the genetic architecture of complex traits.

##### Extended Interpretable Genetic Architecture (Top Genes)

- **Topic 8 (Skin/Psoriasis):** Characterized by *PSORS1C1*, *MUC22*, and *CCHCR1*, this module specifically drives the architecture of Psoriasis and Psoriatic Arthritis, effectively isolating the dermatological genetic component from broader systemic immunity.
- **Topics 9 and 15 (Obesity & Gastrointestinal):** Topic 15 (driven by *SMARCA4*, *LPA*, and the obesity-associated *FTO*) and Topic 9 (driven by *FTO* and *CUX2*) form major structural blocks that co-dominate Obesity, Hyperlipidemia, and Colorectal cancers, illustrating the deep pleiotropy between metabolic syndrome and systemic malignancies.
- **Topics 10, 13, and 14 (Systemic & Malignant):** Topic 10 is driven by *AGT* (Angiotensinogen) and *ZNF831*, linking directly to Hypertension. Topic 13 is characterized by *CDKAL1* and *TCF7L2*, classical

drivers of Type 2 Diabetes. Topic 14 is defined by *FANCA* and *OCA2* (melanin synthesis), strongly aligning with its prominent loading in Melanoma and Skin Cancer.

- **Topics 11 and 12 (Cardiovascular & Specialized Immunity):** Topic 11 is led by *TSBP1*, *XKR6*, and *MSRA*, contributing primarily to Hypertension alongside Topic 10. Topic 12 is defined by *MICB*, *DDX39B*, and *PSORS1C1*, showing a distinct association profile that contributes notably to Cardiovascular Disease and Heart Failure while maintaining strong immune-related gene signatures.

#### Extended Functional Enrichment Analysis

*Metabolic and Endocrine Topics:* Topic 15 demonstrates strong enrichment for "lipid localization", "cholesterol homeostasis", and "Bile secretion", reflecting core lipid metabolism. Topic 13 highlights endocrine functions, significantly enriched for "hormone secretion", "Insulin secretion", and "Maturity onset diabetes of the young", closely aligning with its clinical associations with Type 2 Diabetes.

*Specialized Immune Topics:* Topic 8 isolates specific cytotoxic and transplant-related immune activities, enriched for "Allograft rejection", and specifically captures "MHC class I" pathways. Topic 12 is directly enriched for the "Rheumatoid arthritis" KEGG pathway, while Topic 5 captures "nucleosome assembly" and "Systemic lupus erythematosus".

*Cardiovascular, Structural, and Malignant Topics:* Topic 10 provides a precise mechanistic basis for hypertension, heavily enriched for "regulation of blood pressure", "Renin secretion", and "Adrenergic signaling in cardiomyocytes". Topic 9 identifies structural growth components, enriched for "regulation of chondrocyte differentiation" and "regulation of cartilage development". Finally, Topic 14, driving melanoma and skin cancer risk, is significantly enriched for the "melanin biosynthetic process" and "Cornified envelope formation", accurately capturing the dermatological pathology.

#### Extended Tissue-Specific Expression

- **Specialized Immune Topics (Topics 5 and 12):** The model captures the multi-organ pathology of specific autoimmune diseases exceptionally well. Topic 5 (driving Systemic Lupus Erythematosus) prominently features the "Kidney - Cortex" and "Esophagus - Mucosa", perfectly reflecting complications like lupus nephritis and mucosal involvement. Meanwhile, Topic 12 (strongly associated with Rheumatoid Arthritis) highlights expected immune drivers in "Cells - EBV-transformed lymphocytes", but also reveals profound enrichment in the "Brain - Frontal Cortex (BA9)" and "Adrenal Gland", suggesting a strong genetic interplay between the immune system and neuro-endocrine stress axes.
- **Gastrointestinal and Broad Metabolic Topics (Topics 13 and 15):** Topic 13 (driving Type 2 Diabetes) highlights the "Pancreas", aligning with insulin secretion pathways. Topic 15, which drives Heart Failure and Hyperlipidemia, displays strong expression in systemic metabolic and vascular tissues such as the "Liver", "Artery - Aorta", and "Adipose - Visceral (Omentum)", alongside direct cardiac involvement ("Heart - Left Ventricle"). This indicates that the genetic risk for these cardiovascular traits is driven by a combination of upstream metabolic dysfunction and localized vascular/cardiac biology.
- **Skin and Dermatological Topics (Topics 8 and 14):** Highly localized to "Skin - Sun Exposed (Lower leg)" and "Skin - Not Sun Exposed (Suprapubic)". Topic 8 (driving Psoriasis) captures both systemic immune drivers and target-tissue barrier genes in the skin and gastrointestinal tract ("Colon - Transverse", "Small Intestine"). Topic 14 (Melanoma and Skin Cancer) similarly maps strongly to esophageal and skin tissues, accurately reflecting its underlying epithelial and melanocytic risk factors.
- **Cardiovascular and Systemic Topics (Topics 10 and 11):** These topics provide a highly coherent anatomical basis for blood pressure regulation and hypertensive traits. Topic 10 exhibits striking specificity for the "Adrenal Gland", "Heart - Left Ventricle", and "Heart - Atrial Appendage", reflecting adrenergic signaling and fluid homeostasis. Complementing this, Topic 11 captures broader systemic and structural

remodeling, highlighting “Heart - Left Ventricle” alongside “Cells - Cultured fibroblasts” and mucosal tissues (“Stomach”), pointing toward fibrotic and vascular resistance mechanisms.

- **Structural and Growth Topics (Topic 9):** Displays strong enrichment in “Muscle - Skeletal”, “Kidney - Medulla”, and “Artery - Tibial”, aligning with its functional enrichment in chondrocyte differentiation, structural development, and tissue morphology.

### 4 Figures

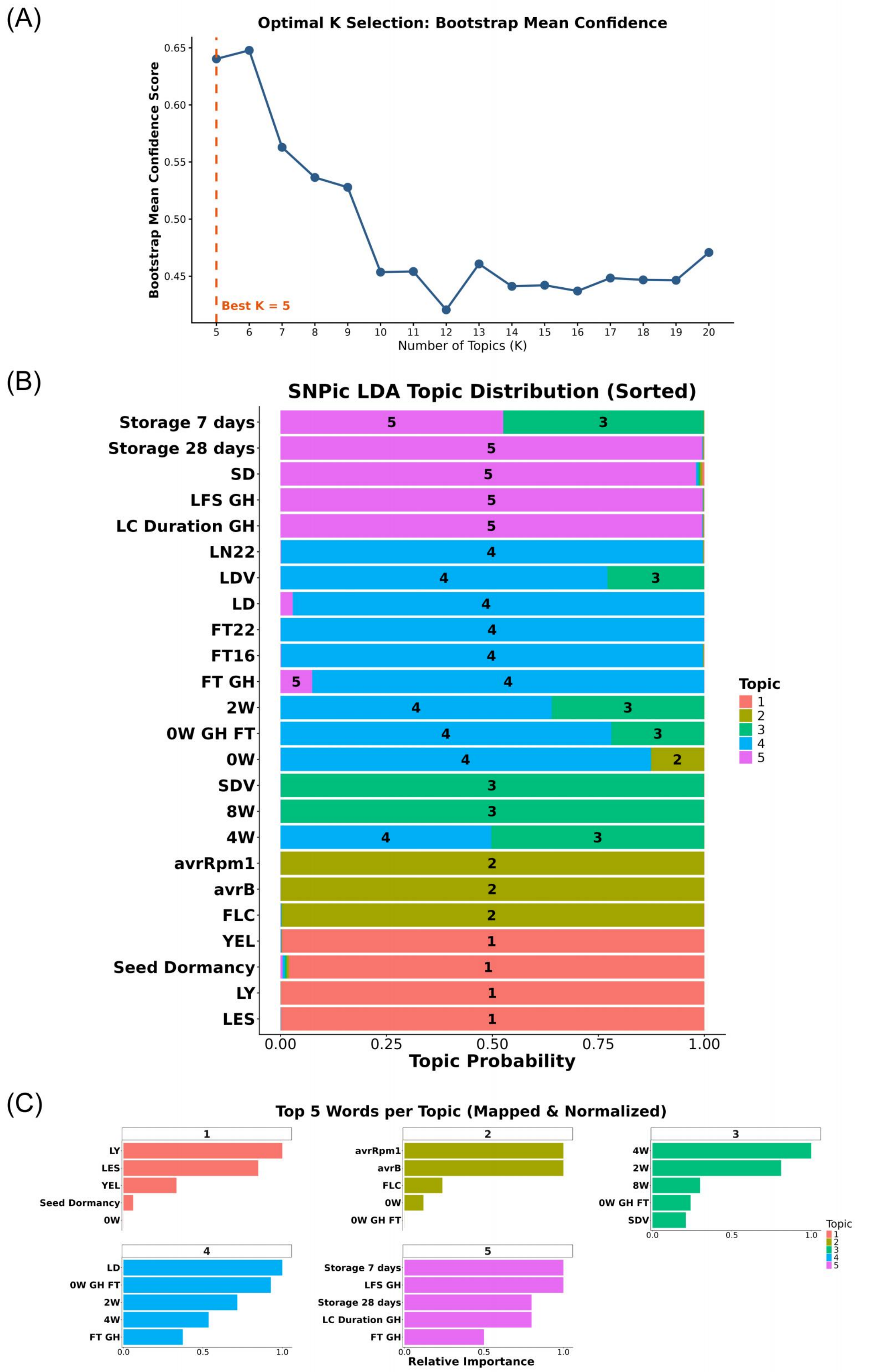


**Fig. 1**: **Topic modeling statistics and feature importance for *Arabidopsis thaliana* GWAS data. (A) Optimal K Selection.** Bootstrap mean confidence evaluation yielding an optimal number of $K = 5$ topics. **(B) Topic Distribution.** Sorted barplot displaying the probability distributions of the 5 latent topics across the analyzed *Arabidopsis* phenotypes. **(C) Top Words per Topic.** The top 5 mapped words (relative importance) driving the classification for each identified topic, highlighting the underlying genetic features defining each functional module.

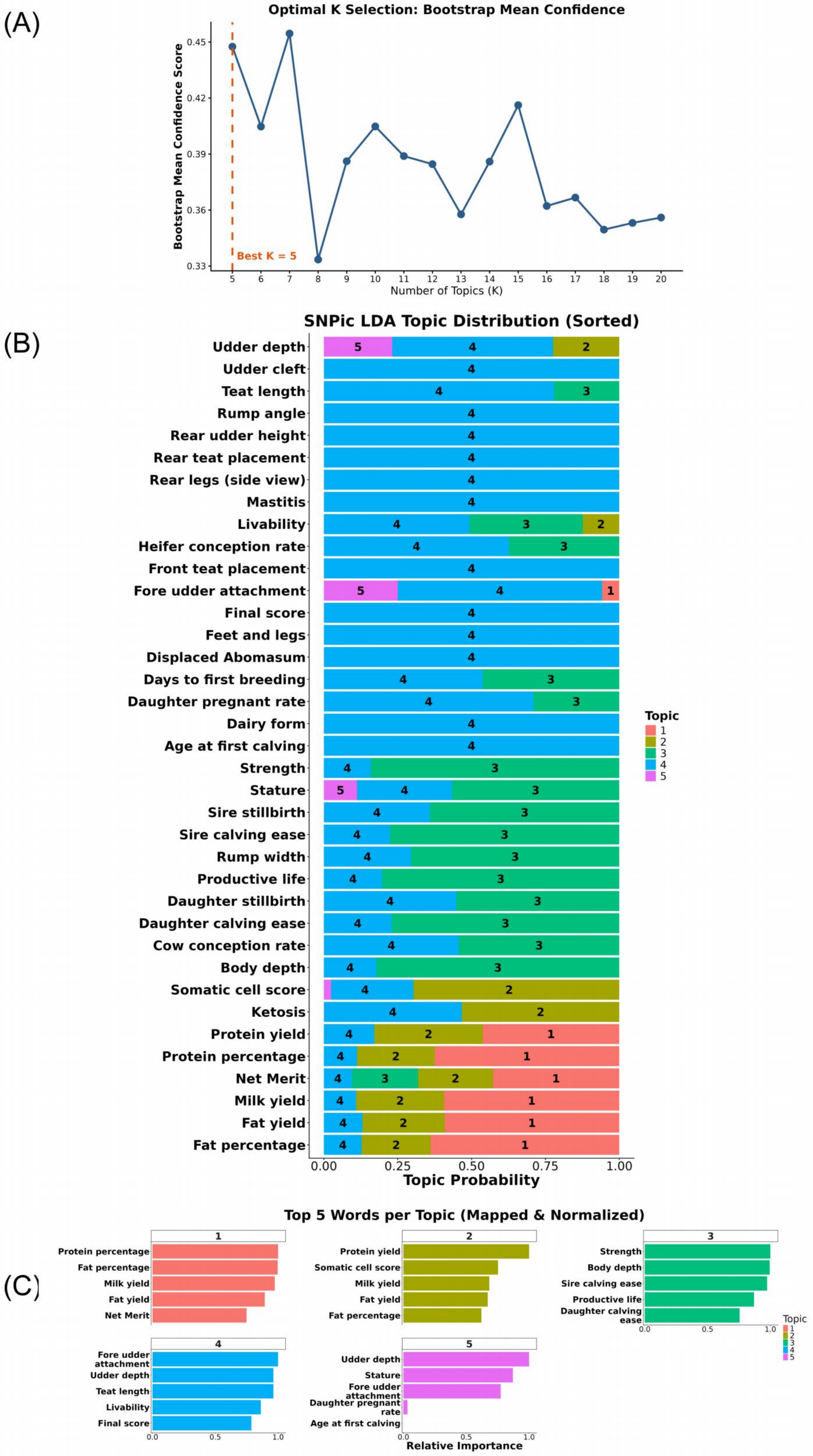


**Fig. 2**: **Topic modeling statistics and feature importance for cattle complex traits. (A) Optimal K Selection.** Bootstrap mean confidence scores indicating an optimal topic number of $K = 5$. **(B) Topic Distribution.** Sorted barplot showing the probability distributions of the 5 topics across the economically important cattle traits. **(C) Top Words per Topic.** The top 5 mapped words (relative importance) that define each of the 5 topics, providing biological and genetic context to the clustered production, health, and reproduction traits.

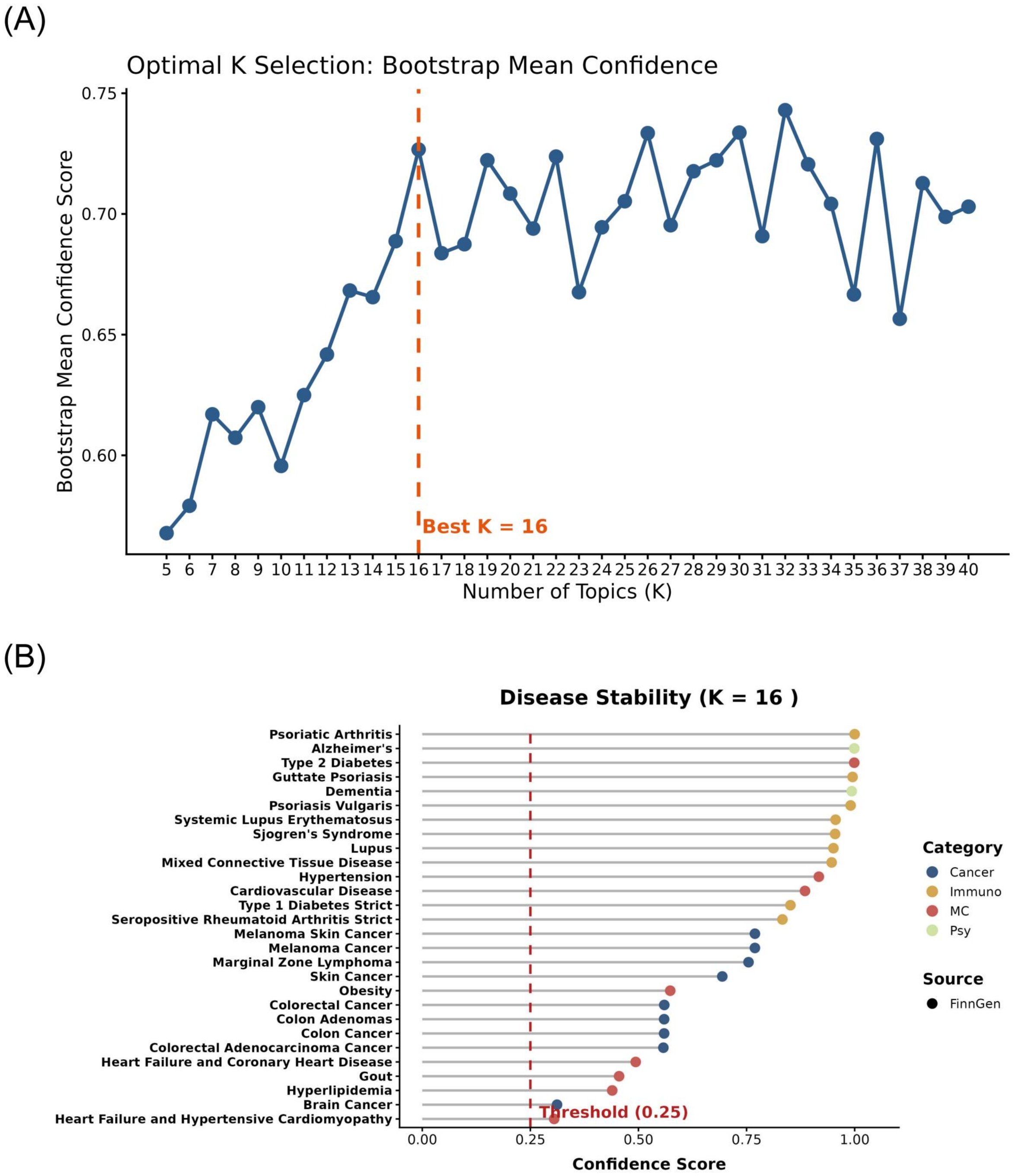


**Fig. 3**: **Stability optimization for the FinnGen-only cohort (*Gene-as-word* LDA).** (**A**) Model stability curve identifying $K = 16$ as the optimal structure. (**B**) Disease stability ranking within the FinnGen dataset.

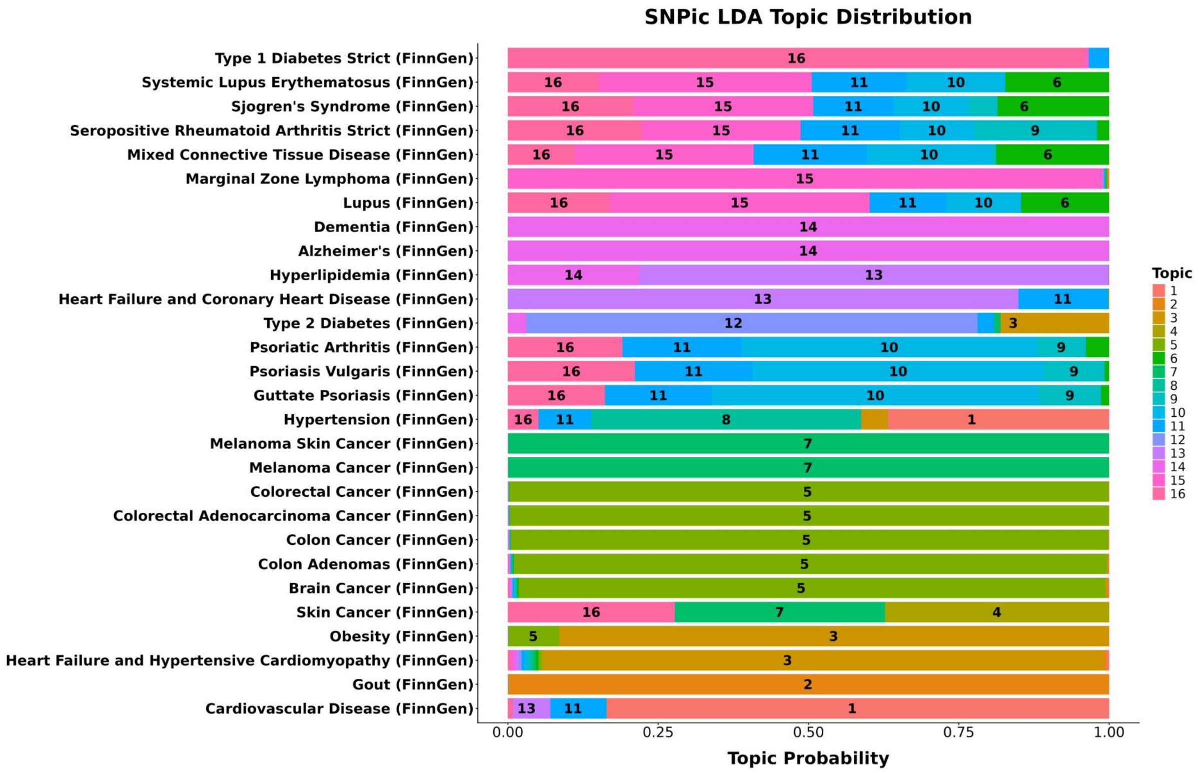


**Fig. 4**: **Topic distribution for the FinnGen-only cohort (*Gene-as-word* LDA).** Topic probability distributions across FinnGen complex traits.

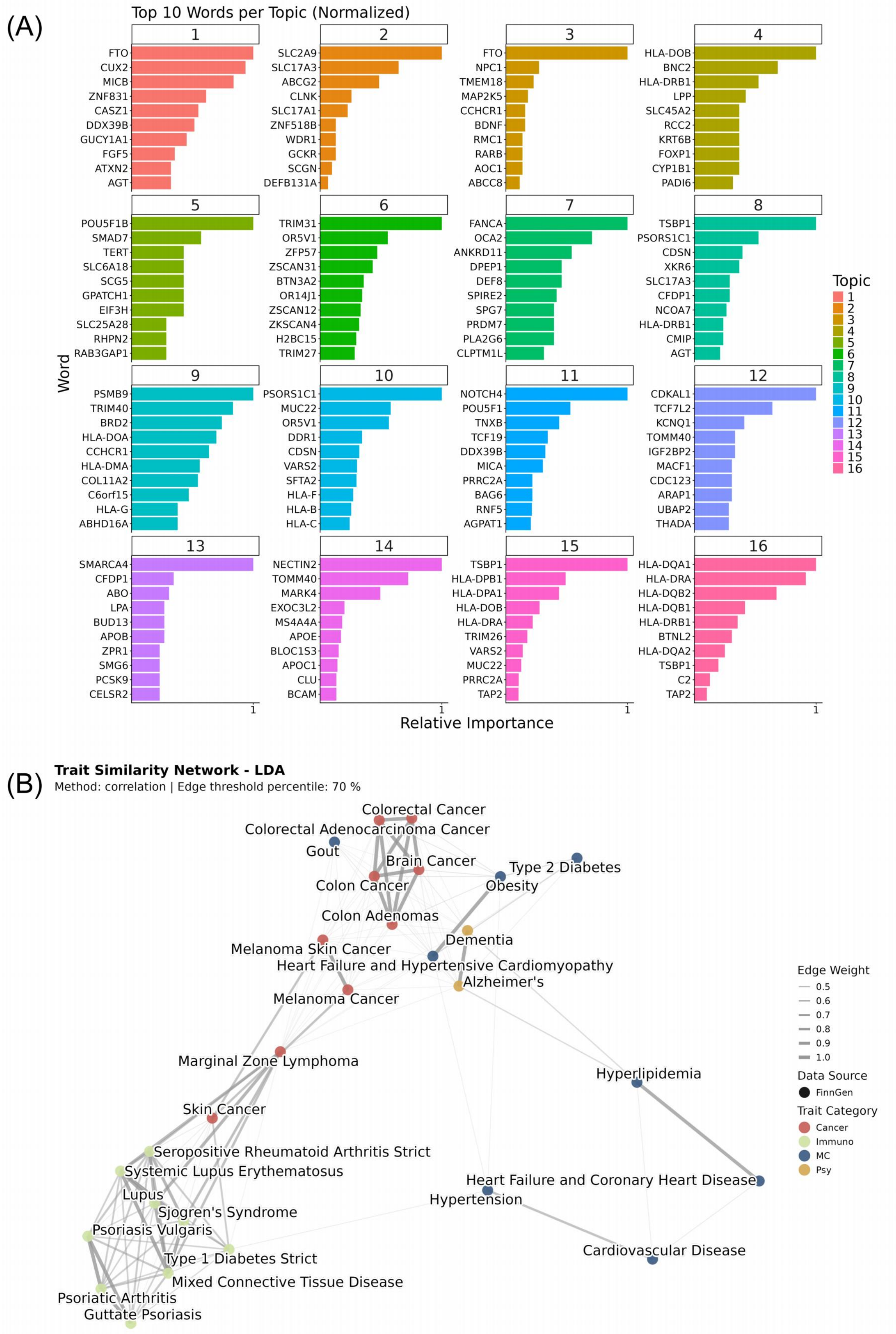


**Fig. 5**: **Genetic architectures for the FinnGen-only cohort (*Gene-as-word* LDA).** (**A**) Top driving genes per topic. (**B**) Disease similarity network for FinnGen traits.

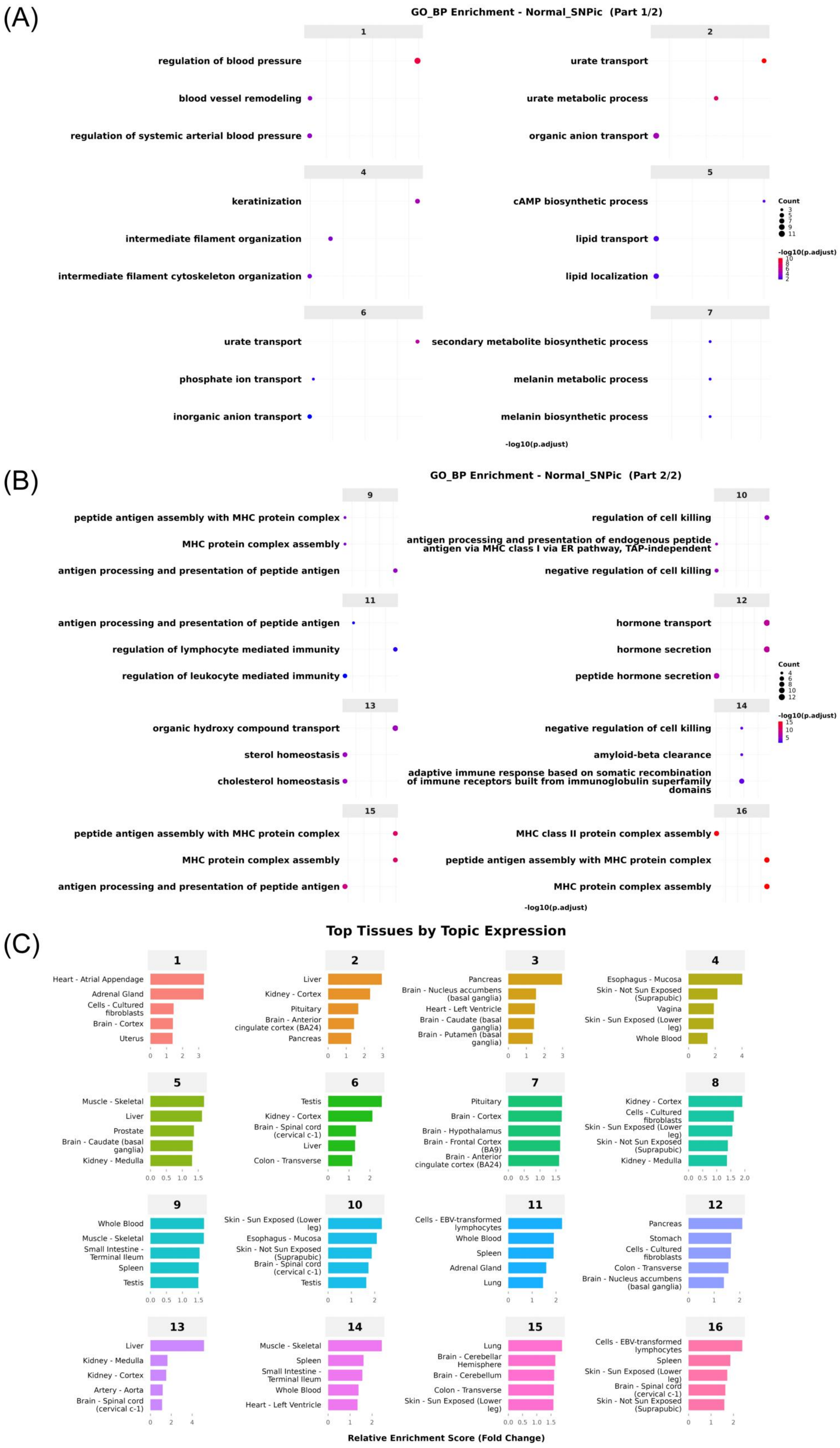


**Fig. 6**: **Biological annotations for the FinnGen-only cohort (*Gene-as-word* LDA).** (**A**) GO pathway enrichments for topics 1-8. (**B**) GO pathway enrichments for topics 9-16. (**C**) Tissue-specific expression profiles.

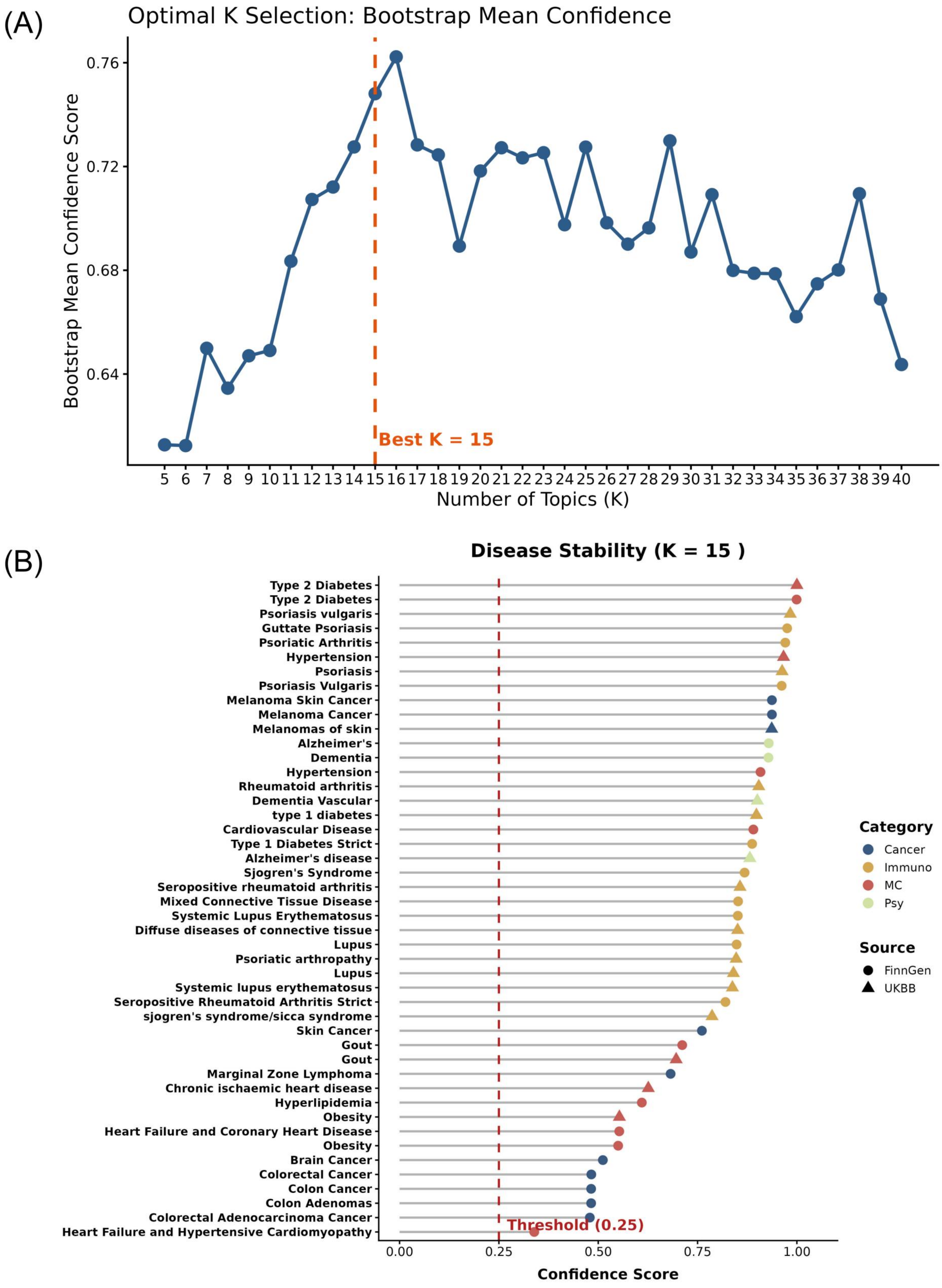


**Fig. 7**: **Stability optimization under the Gaussian Topic Model (*Gene-as-word*).** (**A**) Model stability curve indicating $K = 15$ as the optimal structure. (**B**) Disease stability ranking.

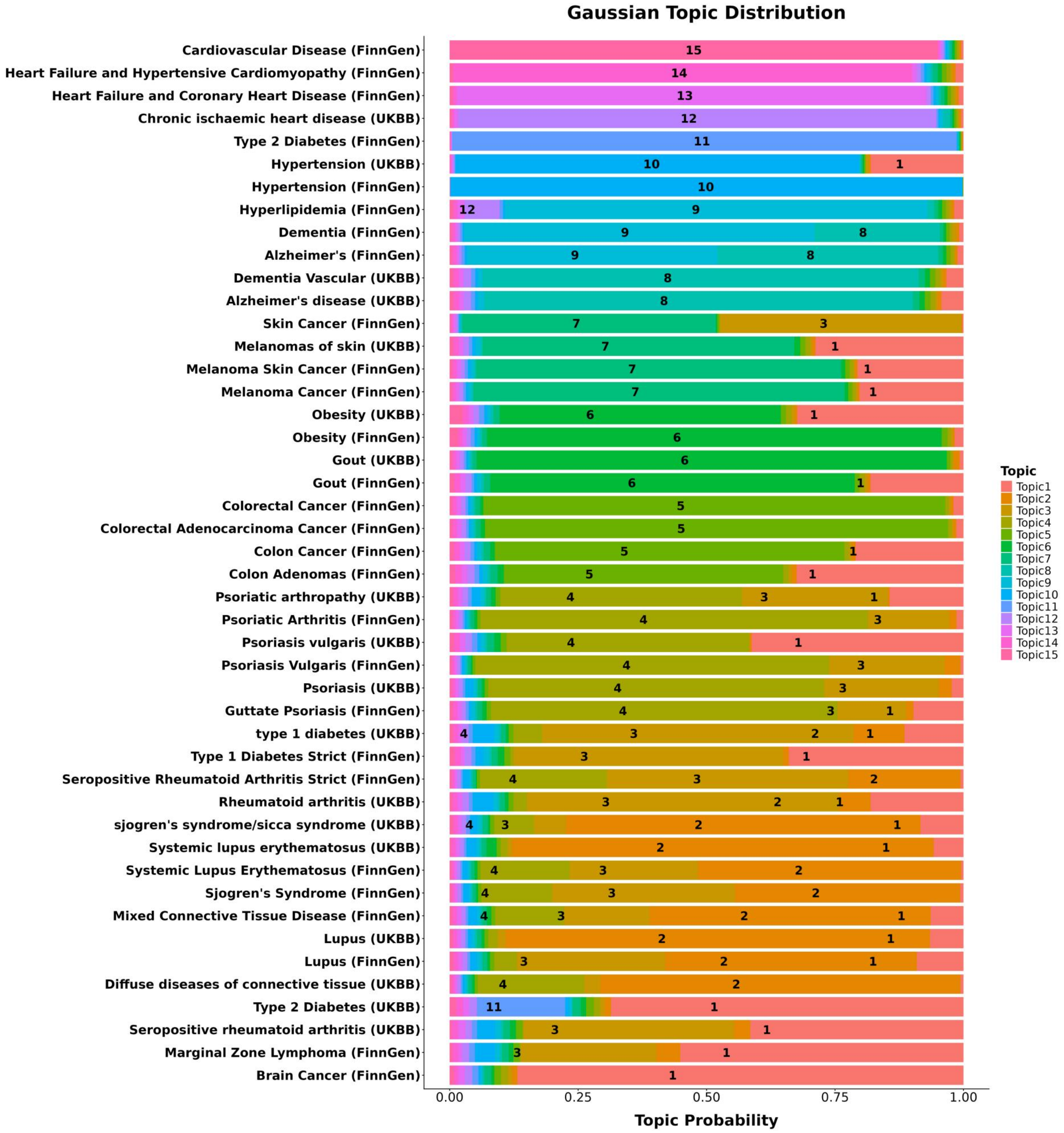


**Fig. 8**: **Topic distribution under the Gaussian Topic Model (*Gene-as-word*).** Topic probability distributions across complex traits.

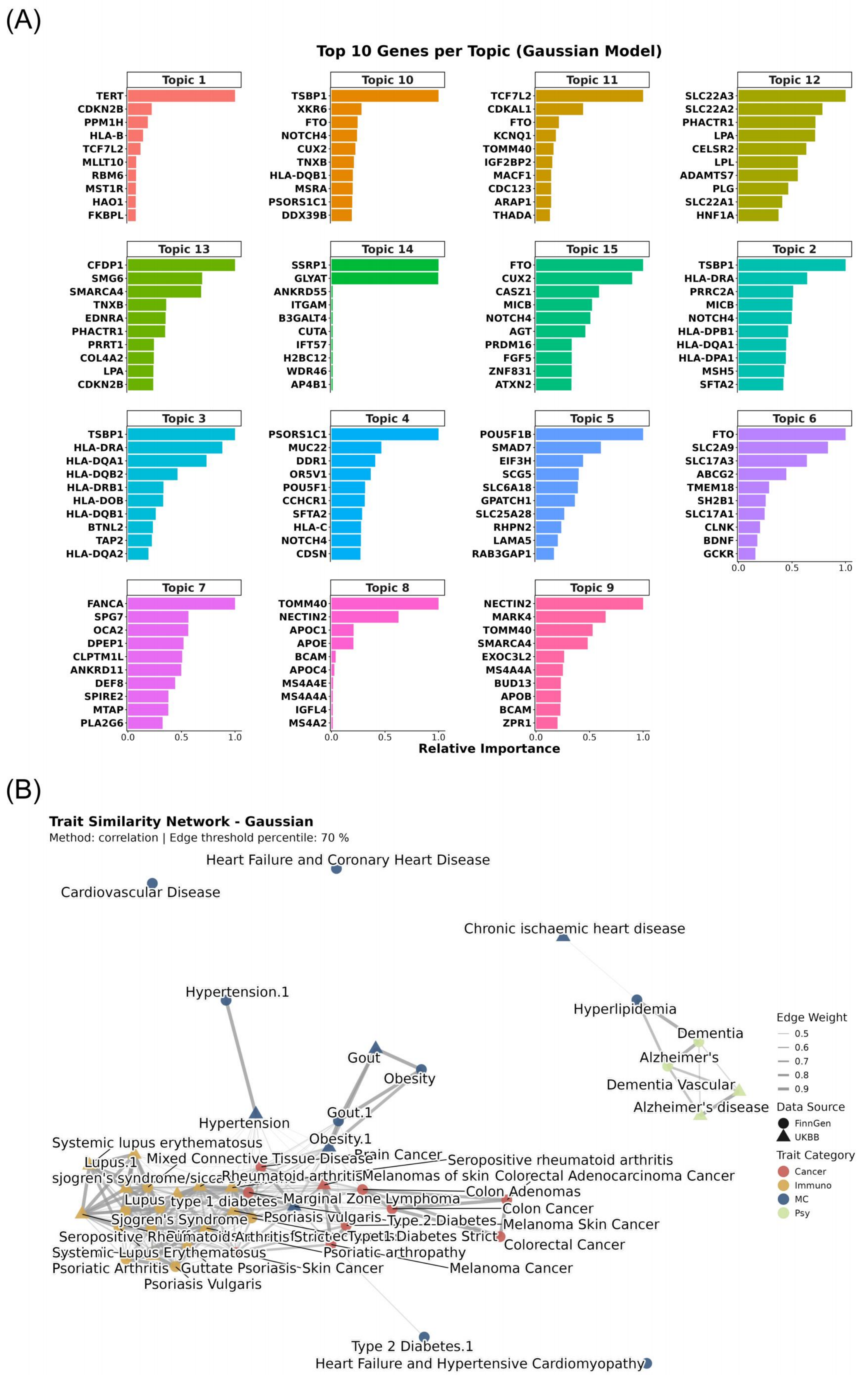


**Fig. 9**: **Genetic architectures under the Gaussian Topic Model (*Gene-as-word*).** (**A**) Top driving genes per topic. (**B**) Disease similarity network.

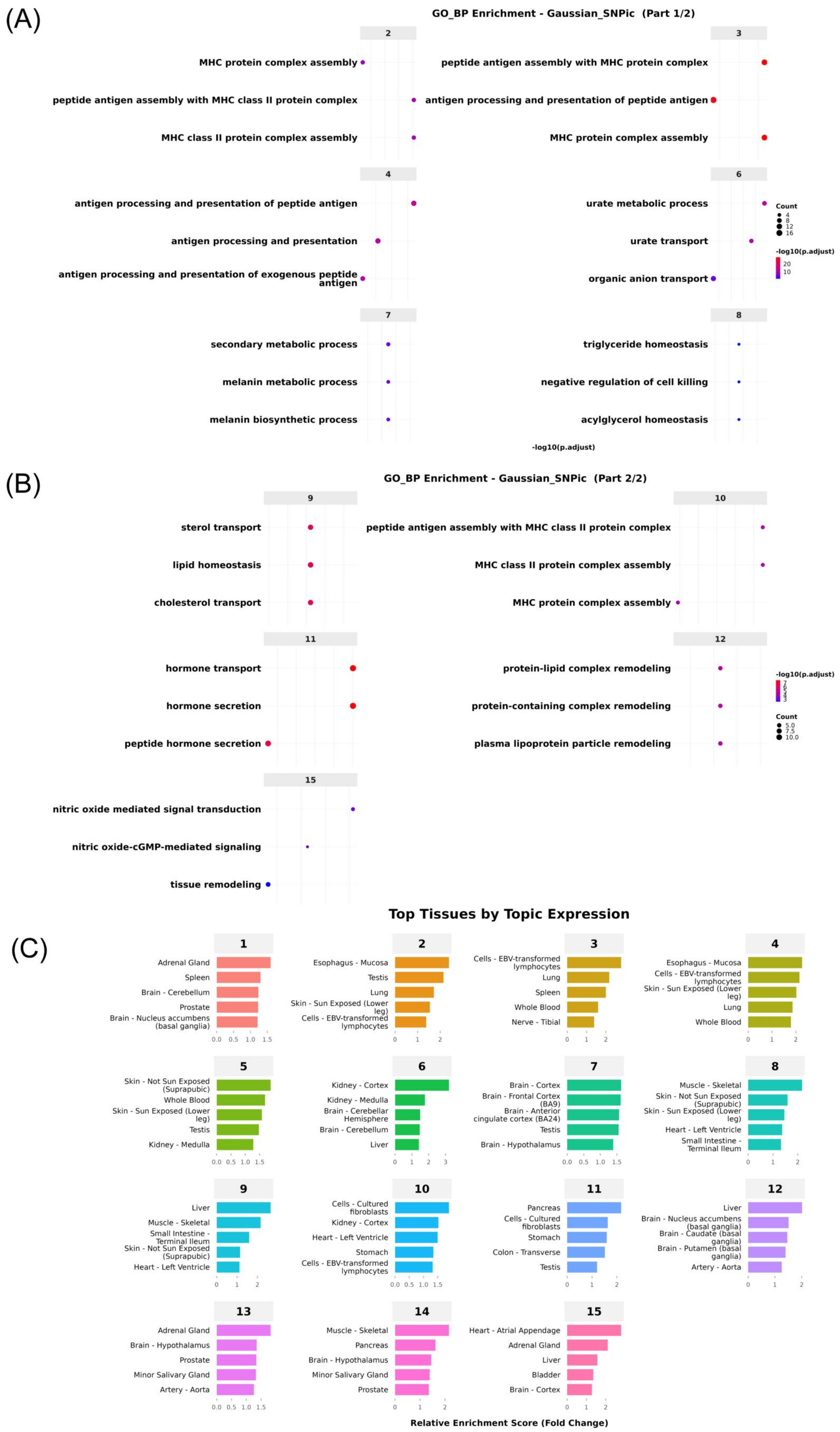


**Fig. 10**: **Biological annotations under the Gaussian Topic Model (*Gene-as-word*).** (**A**) GO pathway enrichments for topics 1-8. (**B**) GO pathway enrichments for topics 9-15. (**C**) Tissue-specific expression profiles.

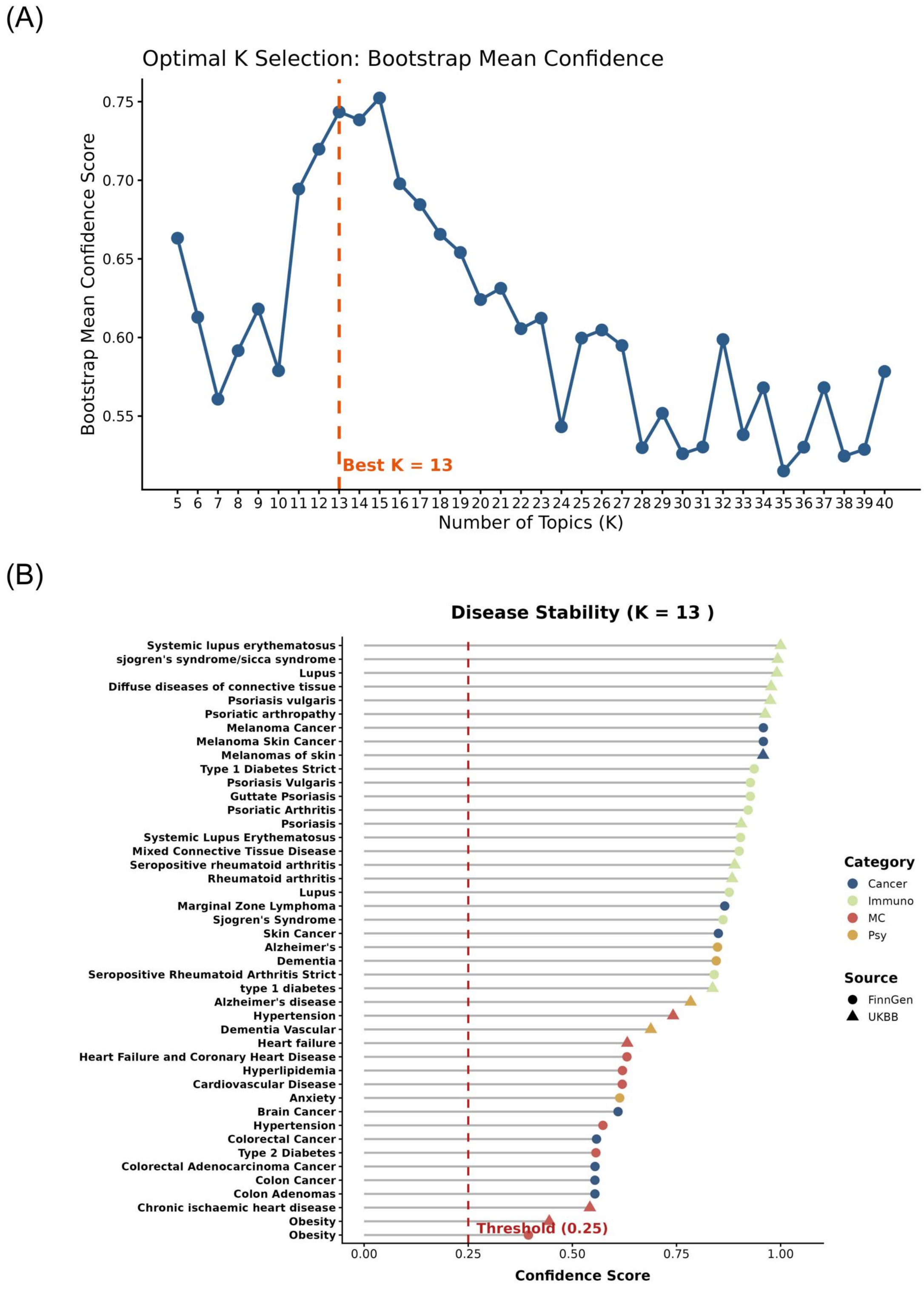


**Fig. 11**: **Stability optimization under the *Sumstat-as-word* LDA scheme.** (**A**) Model stability curve identifying $K = 13$ as the optimal robust structure. (**B**) Disease stability ranking for the optimal model. A dynamic threshold (red dashed line) filters out unstable phenotypes.

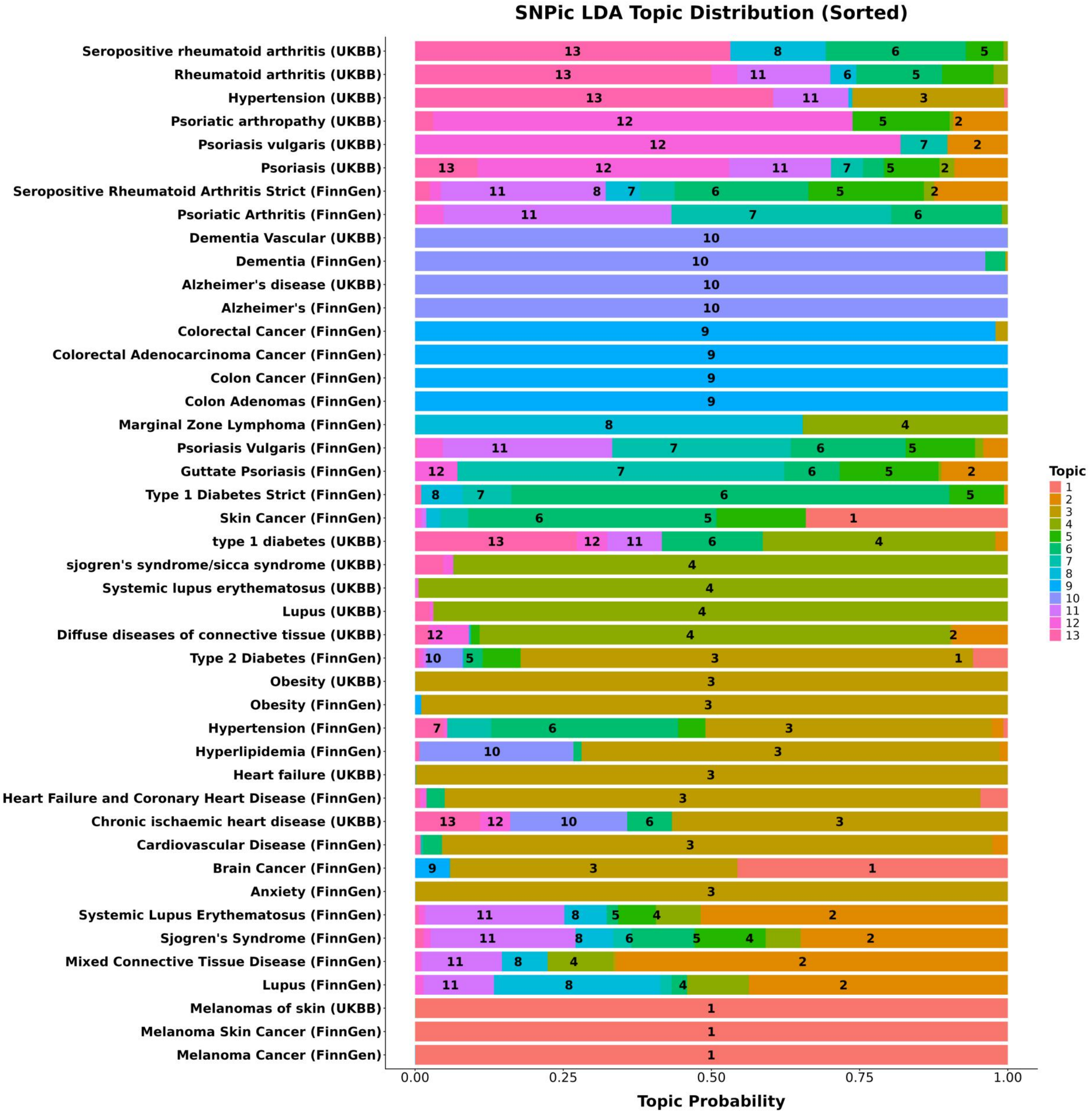


**Fig. 12**: **Topic distribution under the *Sumstat-as-word* LDA scheme.** Topic probability distributions across complex traits.

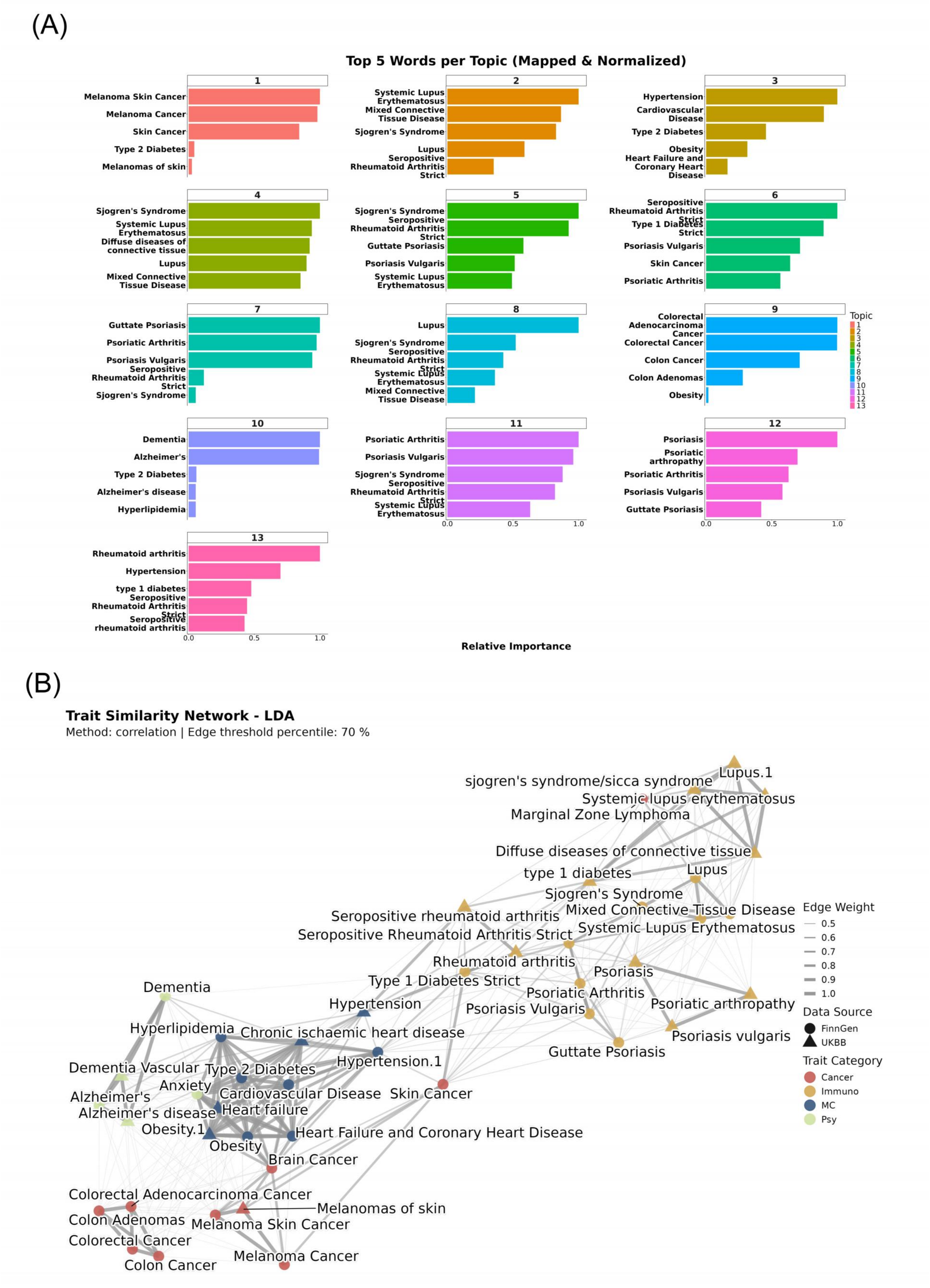


**Fig. 13**: **Genetic architectures under the *Sumstat-as-word* LDA scheme.** (**A**) Top driving variants per topic. (**B**) Disease similarity network based on shared SNP architectures. Nodes are colored by broad disease categories.

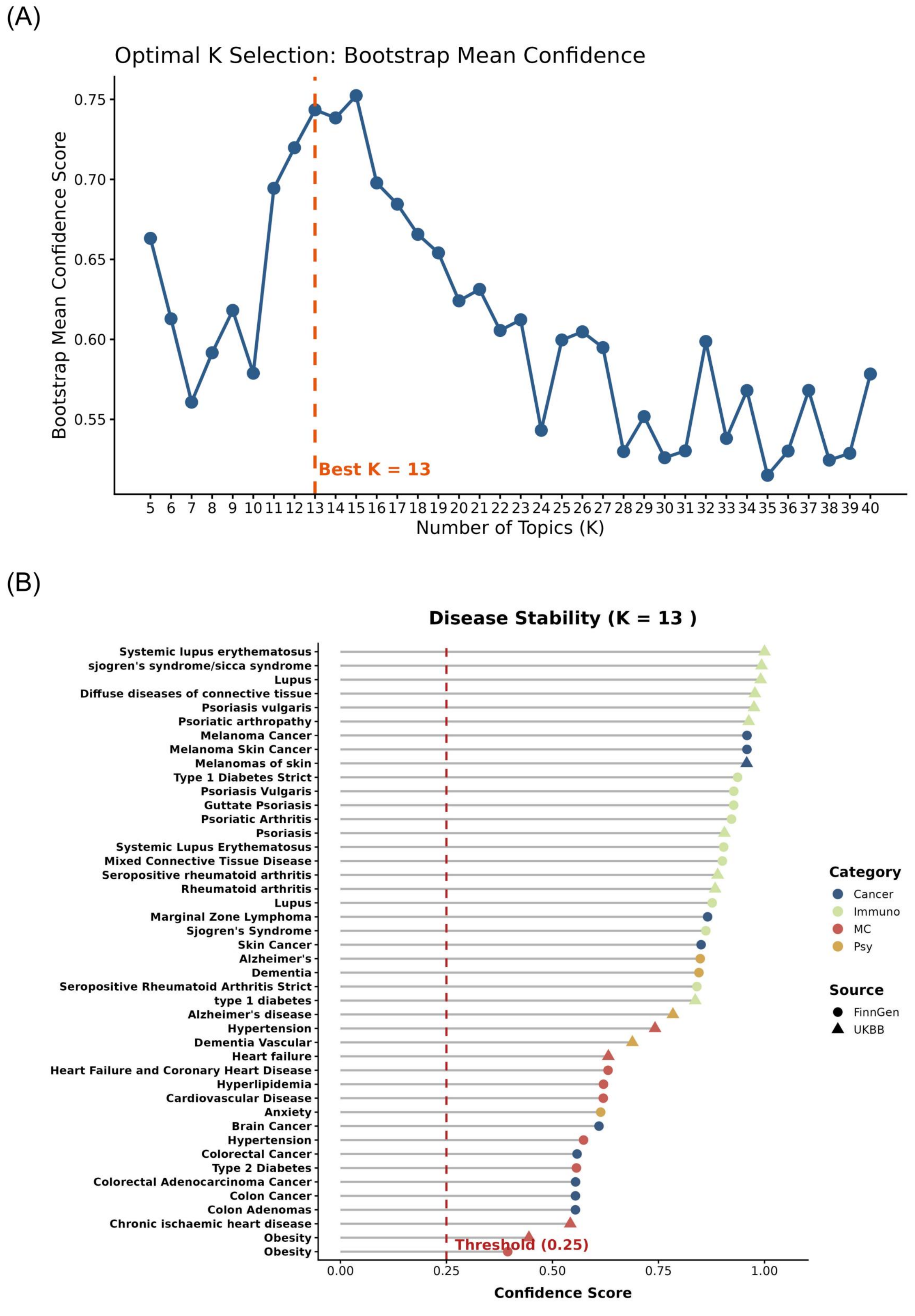


**Fig. 14**: **Stability optimization under the *Sumstat-as-word* Gaussian Topic Model.** (**A**) Model stability curve identifying $K = 13$ as the optimal structure. (**B**) Disease stability ranking.

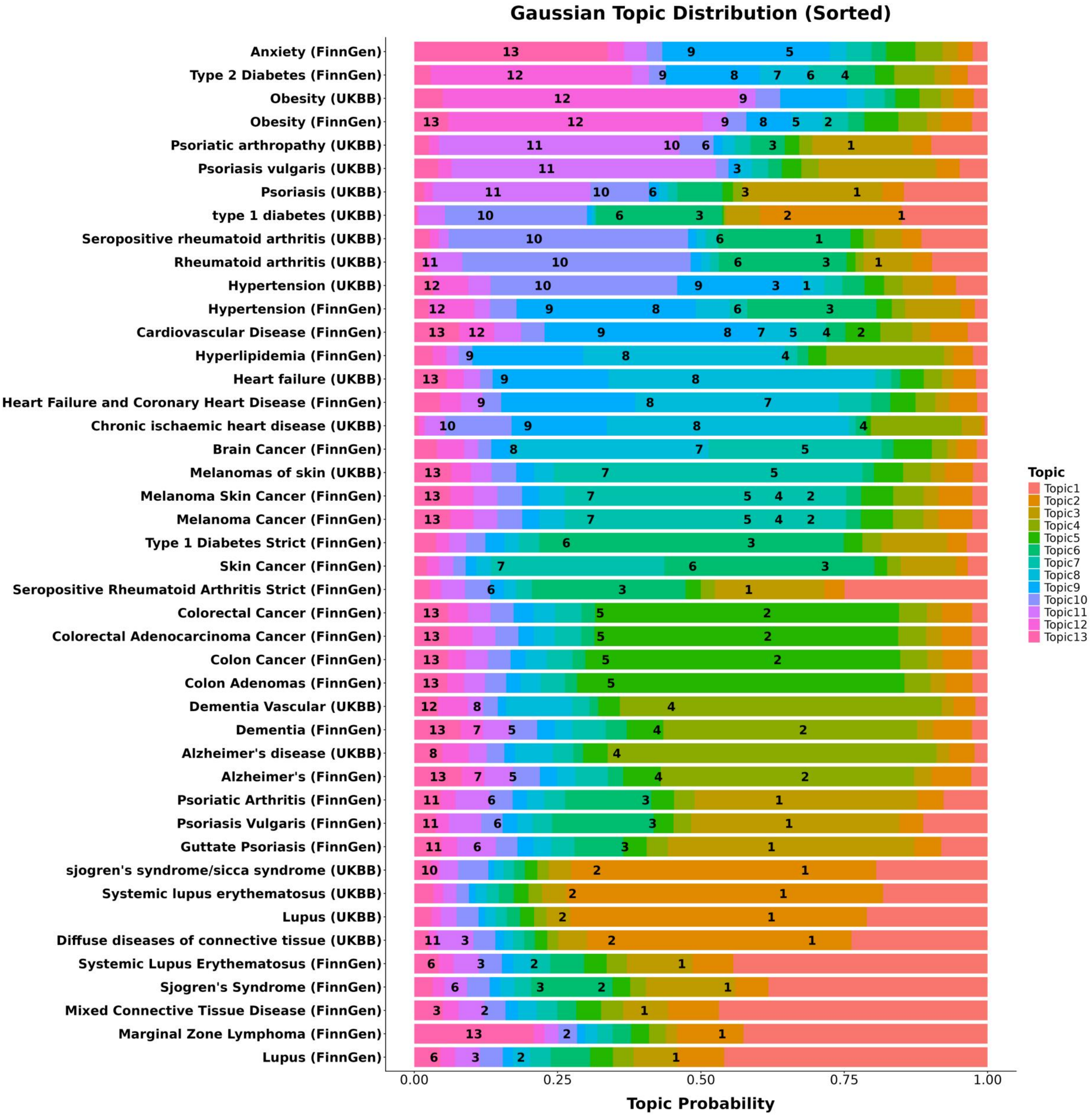


**Fig. 15**: **Topic distribution under the *Sumstat-as-word* Gaussian Topic Model.** Topic probability distributions across complex traits.

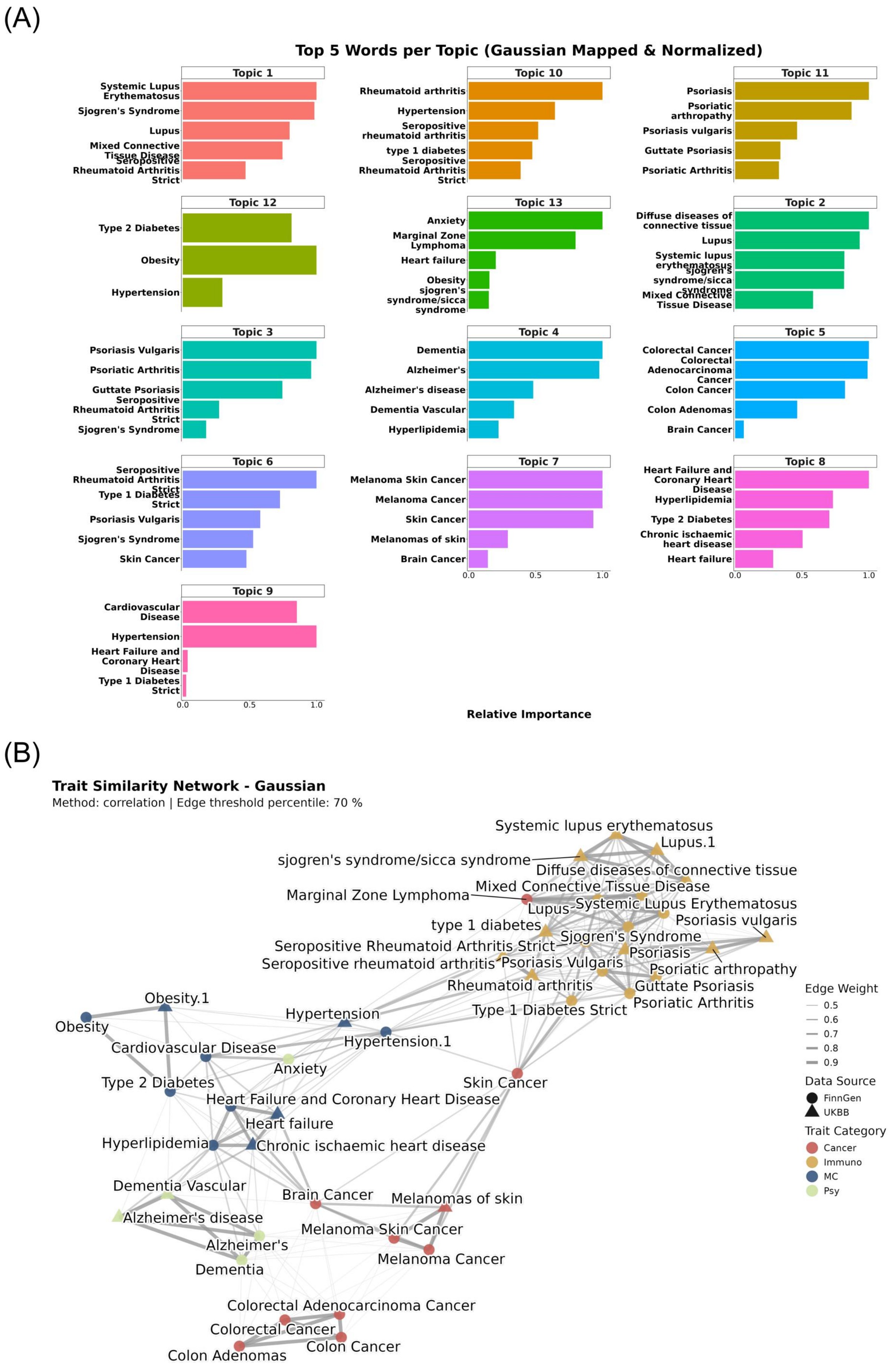


**Fig. 16**: **Genetic architectures under the *Sumstat-as-word* Gaussian Topic Model.** (**A**) Top driving variants per topic. (**B**) Disease similarity network.

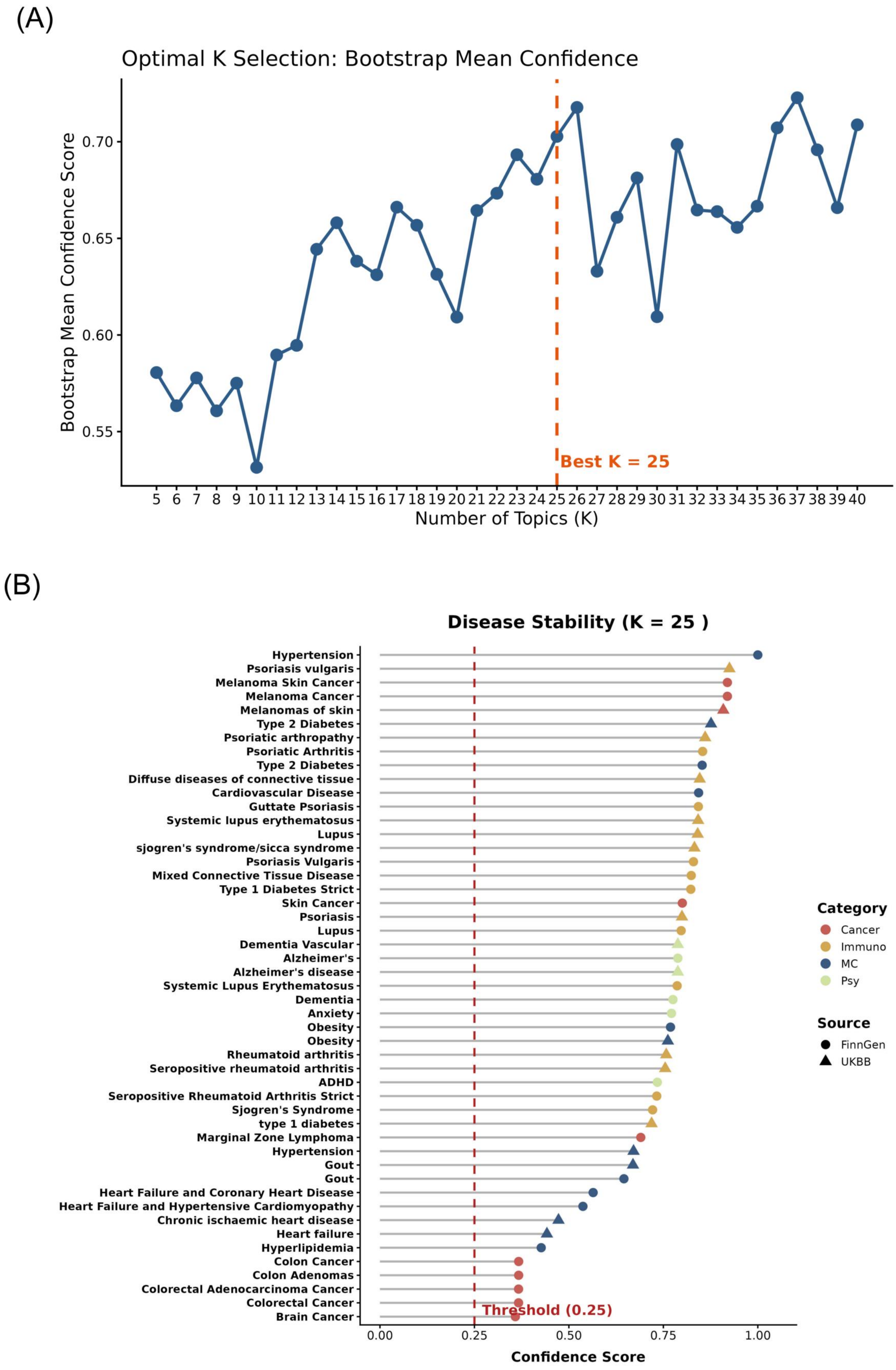


**Fig. 17**: **Stability optimization integrating GTEx multi-tissue V2F mapping (*Gene-as-word* LDA).** (**A**) Model stability curve. The optimal structure shifts to $K = 25$, reflecting network fragmentation caused by off-target tissue noise. (**B**) Disease stability ranking.

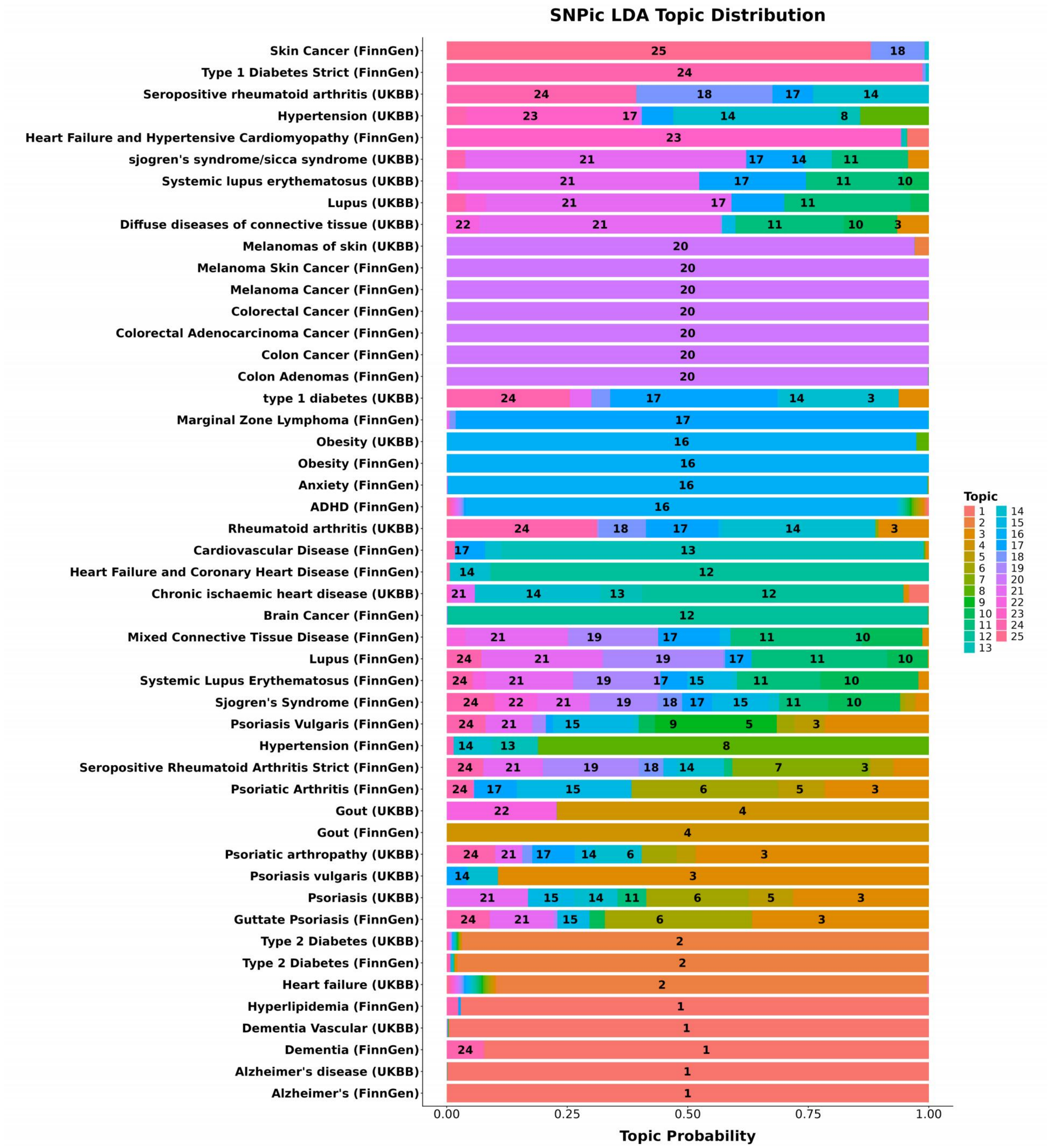


**Fig. 18**: **Topic distribution integrating GTEx V2F mapping (*Gene-as-word* LDA).** The incorporation of global eQTLs artificially hybridizes distinct disease categories.

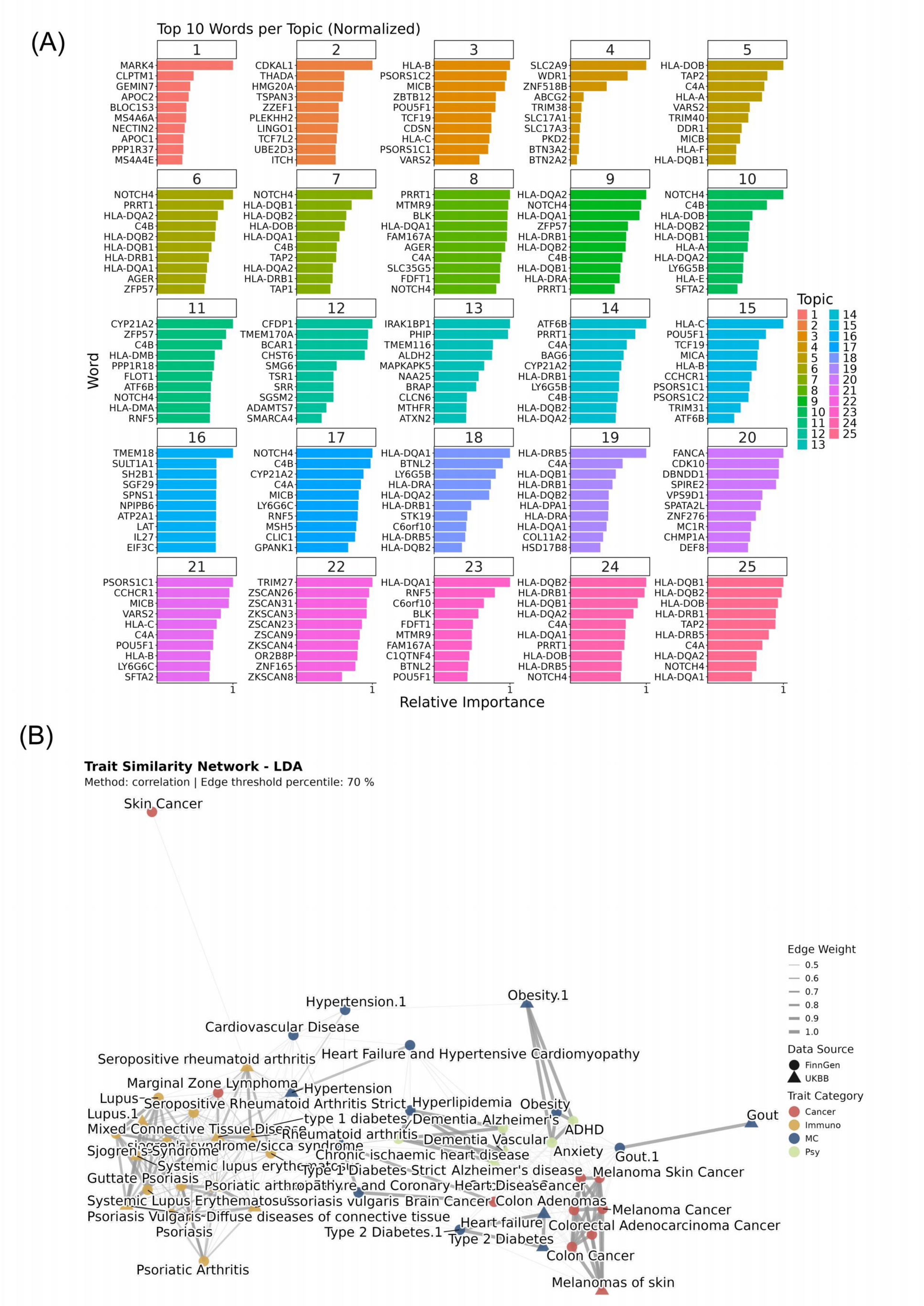


**Fig. 19**: **Genetic architectures integrating GTEx V2F mapping (*Gene-as-word* LDA). (A)** Top driving genes per topic. (**B**) Disease similarity network showing the structural impact of complex variant-to-function mapping.

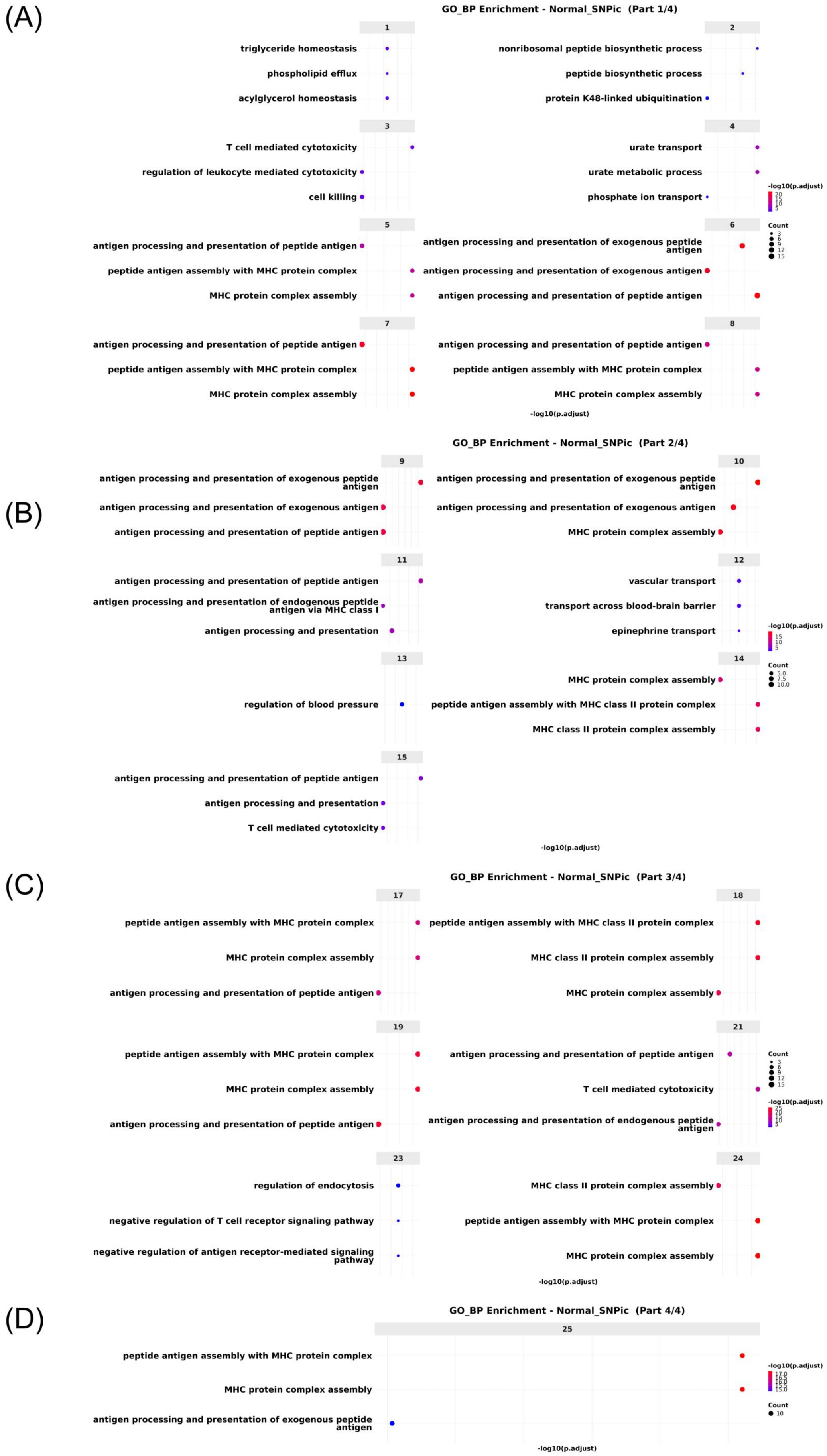


**Fig. 20**: **Biological annotations based on GO Pathway Enrichment integrating GTEx V2F mapping (*Gene-as-word* LDA).**

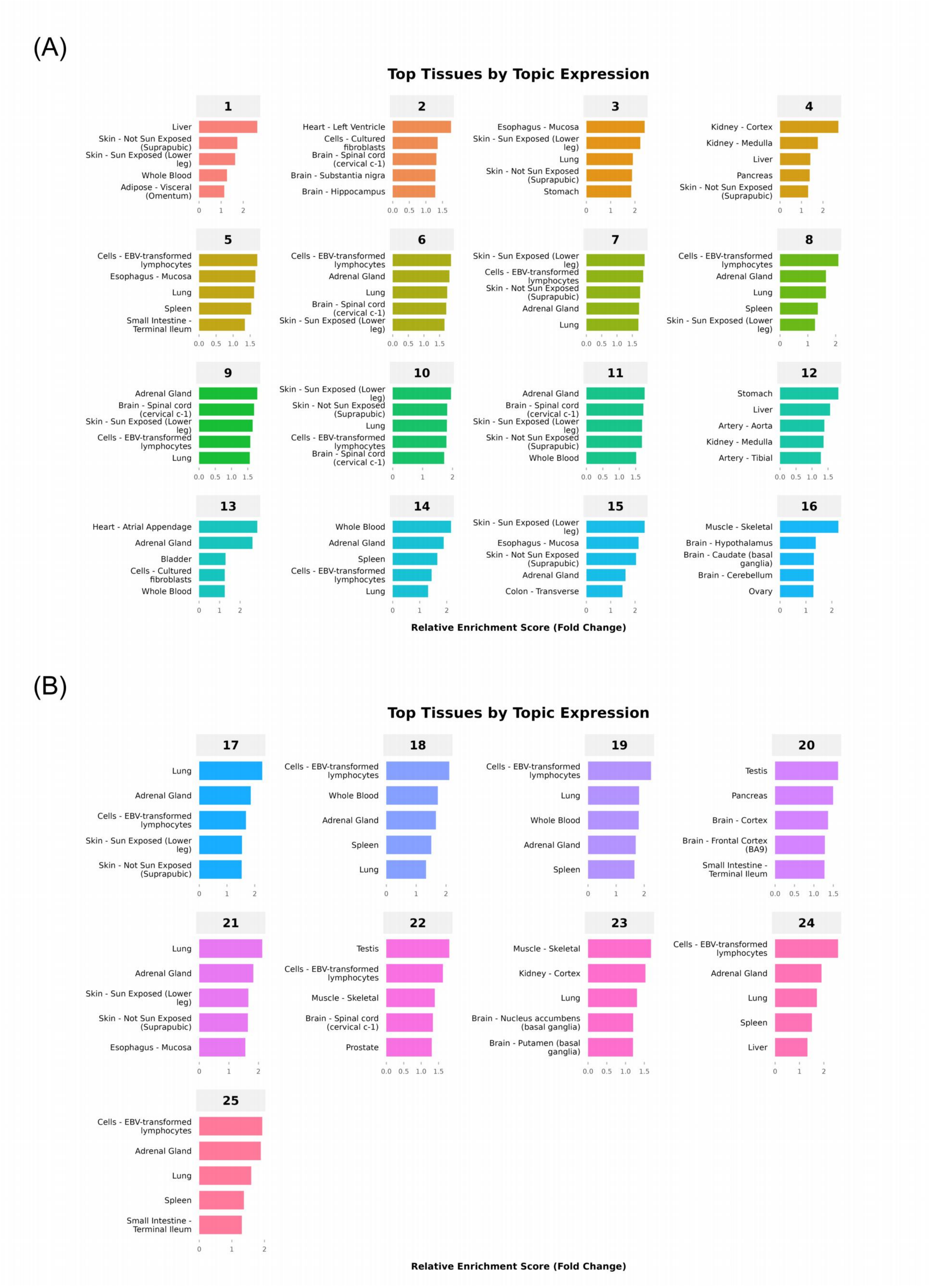


**Fig. 21**: **Biological annotations based on Tissue-specific Expression integrating GTEx V2F mapping (*Gene-as-word* LDA).**

### 5 Tables

**Table 1**: **Descriptions of *Arabidopsis thaliana* Phenotypes.** Detailed descriptions and growth conditions of the phenotypes present in the trait similarity network, as originally defined by Atwell et al. [56].

| Abbreviation | Phenotype Name | Description & Growth Conditions |
|---|---|---|
| 0W | Days to flowering (0W) | Days to flowering time under Long Day (16 hrs daylight) at 23°$C$. Not vernalized. |
| 2W | Days to flowering (2W) | Days to flowering time under Long Day. Vernalized for 2 weeks at 5°$C$. |
| 4W | Days to flowering (4W) | Days to flowering time under Long Day. Vernalized for 4 weeks at 5°$C$. |
| 8W | Days to flowering (8W) | Days to flowering time under Long Day. Vernalized for 8 weeks at 5°$C$. |
| LD | Days to flowering (LD) | Days to flowering time under Long Day (18°$C$, 16 hrs daylight). |
| LDV | Days to flowering (LDV) | Days to flowering time under Long Day. Vernalized for 5 weeks at 4°$C$. |
| SD | Days to flowering (SD) | Days to flowering time under Short Day (18°$C$, 8 hrs daylight). |
| SDV | Days to flowering (SDV) | Days to flowering time under Short Day. Vernalized for 5 weeks at 4°$C$. |
| FT16 | Flowering time (16°$C$) | Days to flowering time of plants grown at 16°$C$, 16 hrs daylight. |
| FT22 | Flowering time (22°$C$) | Days to flowering time of plants grown at 22°$C$, 16 hrs daylight. |
| LN22 | Leaf number (22°$C$) | Average leaf number at flowering time of plants grown at 22°$C$. |
| 0W GH FT | Days to flowering (GH) | Days to flowering time under natural light in the greenhouse. Not vernalized. |
| FT GH | Days to flowering (GH) | Days to flowering time of plants grown in the greenhouse (20°$C$, 16 hrs daylight). |
| LC Duration GH | Life cycle period | Number of days between germination and plant complete senescence. |
| LFS GH | Last flower senescence | Number of days between germination and senescence of the last flower. |
| Seed Dormancy | Seed dormancy level | Number of days of seed dry storage required to reach 50% germination rate. |
| Storage 7 days | Primary dormancy (7d) | Primary dormancy measured as the progressive increase of germination rate after 7 days of dry storage. |
| Storage 28 days | Primary dormancy (28d) | Primary dormancy measured as the progressive increase of germination rate after 28 days of dry storage. |
| LES | Lesioning | Presence or absence of lesioning (necrotic spots spreading to the whole leaf). |

Continued on next page

**Table 1 – continued from previous page**

| Abbreviation | Phenotype Name | Description & Growth Conditions |
|---|---|---|
| YEL | Yellowing | Presence or absence of yellowing (precocious, diffused leaf chlorosis). |
| LY | Lesioning or Yellowing | Presence or absence of either lesioning or yellowing. |
| FLC | *FLC* gene expression | *FLC* gene expression levels extracted from leaves after 4 weeks of growth. |
| avrB | Hypersensitive Response | Disease presence or absence following inoculation with *Pseudomonas syringae* expressing AvrB. |
| avrRpm1 | Hypersensitive Response | Disease presence or absence following inoculation with *Pseudomonas syringae* expressing AvrRpm1. |

**Table 2**: **Cattle Complex Traits Analyzed in SNPic.** List of the 43 economically important cattle traits obtained from the farmGTEx dataset [59], organized by their functional categories as visualized in the trait similarity network.

| Trait Abbreviation | Full Trait Name | Category |
|---|---|---|
| Body_depth_score | Body depth | Body Conformation |
| Dairy_form | Dairy form | Body Conformation |
| Final_score | Final classification score (PTAT) | Body Conformation |
| Foot_angle | Foot angle | Body Conformation |
| Fore_udder_attachment | Fore udder attachment | Body Conformation |
| Front_teat_placement | Front teat placement | Body Conformation |
| Rear_legs_rear_view | Rear legs rear view | Body Conformation |
| Rear_legs_side_view | Rear legs side view | Body Conformation |
| Rear_teat_placement | Rear teat placement | Body Conformation |
| Rear_udder_height | Rear udder height | Body Conformation |
| Rear_udder_width | Rear udder width | Body Conformation |
| Rump_angle | Rump angle | Body Conformation |
| Stature | Stature | Body Conformation |
| Strength | Strength | Body Conformation |
| Teat_length | Teat length | Body Conformation |
| Thurl_width | Thurl width | Body Conformation |
| Udder_cleft | Udder cleft | Body Conformation |
| Udder_depth | Udder depth | Body Conformation |
| Displaced_Abomasum | Displaced abomasum | Health Lifespan |
| Ketosis | Ketosis | Health Lifespan |
| Livability | Livability | Health Lifespan |
| Mastitis | Mastitis | Health Lifespan |
| Metritis | Metritis | Health Lifespan |
| Milk_Fever | Milk fever (Hypocalcemia) | Health Lifespan |
| PL_DYD | Productive life | Health Lifespan |
| Retained_Placenta | Retained placenta | Health Lifespan |
| SCS_DYD | Somatic cell score | Health Lifespan |

Continued on next page

**Table 2 – continued from previous page**

| Trait Abbreviation | Full Trait Name | Category |
|---|---|---|
| FP_DYD | Fat percentage | Milk Production |
| FY_DYD | Fat yield | Milk Production |
| MY_DYD | Milk yield | Milk Production |
| PP_DYD | Protein percentage | Milk Production |
| PY_DYD | Protein yield | Milk Production |
| AFC_DYD | Age at first calving | Reproduction Fertility |
| CCR_DYD | Cow conception rate | Reproduction Fertility |
| CI_DYD | Calving interval | Reproduction Fertility |
| DCE_DYD | Daughter calving ease | Reproduction Fertility |
| DPR_DYD | Daughter pregnancy rate | Reproduction Fertility |
| DSB_DYD | Daughter stillbirth | Reproduction Fertility |
| GL_DYD | Gestation length | Reproduction Fertility |
| HCR_DYD | Heifer conception rate | Reproduction Fertility |
| SCE_DYD | Sire calving ease | Reproduction Fertility |
| SSB_DYD | Sire stillbirth | Reproduction Fertility |

**Table 3**: **Human Complex Traits Analyzed in SNPic.** List of the 47 human diseases and traits included in the cross-biobank integration analysis, organized by disease category and data source. Abbreviations: **MC** = Metabolic/Cardio, **Psy** = Psychiatric/Neuro.

| Trait Name | Category | Data Source |
|---|---|---|
| Brain Cancer | Cancer | FinnGen |
| Colon Adenomas | Cancer | FinnGen |
| Colon Cancer | Cancer | FinnGen |
| Colorectal Adenocarcinoma Cancer | Cancer | FinnGen |
| Colorectal Cancer | Cancer | FinnGen |
| Marginal Zone Lymphoma | Cancer | FinnGen |
| Melanoma Cancer | Cancer | FinnGen |
| Melanoma Skin Cancer | Cancer | FinnGen |
| Skin Cancer | Cancer | FinnGen |
| Melanomas of skin | Cancer | UKBB |
| Diffuse diseases of connective tissue | Immuno | FinnGen |
| Guttate Psoriasis | Immuno | FinnGen |
| Mixed Connective Tissue Disease | Immuno | FinnGen |
| Psoriasis Vulgaris | Immuno | FinnGen |
| Psoriatic Arthritis | Immuno | FinnGen |
| Seropositive Rheumatoid Arthritis Strict | Immuno | FinnGen |
| Sjogren's Syndrome | Immuno | FinnGen |
| Systemic Lupus Erythematosus | Immuno | FinnGen |
| Type 1 Diabetes Strict | Immuno | FinnGen |
| Lupus | Immuno | UKBB |

Continued on next page

**Table 3 – continued from previous page**

| Trait Name | Category | Data Source |
|---|---|---|
| Psoriasis | Immuno | UKBB |
| Psoriasis vulgaris | Immuno | UKBB |
| Psoriatic arthropathy | Immuno | UKBB |
| Rheumatoid arthritis | Immuno | UKBB |
| Seropositive rheumatoid arthritis | Immuno | UKBB |
| Systemic lupus erythematosus | Immuno | UKBB |
| sjogren's syndrome/sicca syndrome | Immuno | UKBB |
| type 1 diabetes | Immuno | UKBB |
| Cardiovascular Disease | MC | FinnGen |
| Gout | MC | FinnGen |
| Heart Failure and Coronary Heart Disease | MC | FinnGen |
| Heart Failure and Hypertensive Cardiomyopathy | MC | FinnGen |
| Hyperlipidemia | MC | FinnGen |
| Hypertension | MC | FinnGen |
| Obesity | MC | FinnGen |
| Type 2 Diabetes | MC | FinnGen |
| Chronic ischaemic heart disease | MC | UKBB |
| gout | MC | UKBB |
| hypertension | MC | UKBB |
| type 2 diabetes | MC | UKBB |
| Alzheimer's | Psy | FinnGen |
| Dementia | Psy | FinnGen |
| Alzheimer's disease | Psy | UKBB |
| Dementia Vascular | Psy | UKBB |

**Table 4**: **Functional Enrichment of SNPic Topics.** Top Gene Ontology Biological Processes (GO) and KEGG pathways enriched within the core gene sets of the 15 inferred genetic topics. *p.adjust* indicates the false discovery rate (FDR) corrected p-value.

| Topic | Analysis | ID | Description | p.adjust |
|---|---|---|---|---|
| Topic 1 | GO | GO:0002396 | MHC protein complex assembly | $8.35 \times 10^{-7}$ |
| | GO | GO:0002501 | peptide antigen assembly with MHC protein complex | $8.35 \times 10^{-7}$ |
| | GO | GO:0048002 | antigen processing and presentation of peptide antigen | $3.62 \times 10^{-6}$ |
| | KEGG | hsa04612 | Antigen processing and presentation | $5.24 \times 10^{-9}$ |
| | KEGG | hsa04145 | Phagosome | $4.38 \times 10^{-7}$ |
| | KEGG | hsa05150 | Staphylococcus aureus infection | $5.78 \times 10^{-7}$ |
| Topic 2 | GO | GO:0046415 | urate metabolic process | $2.04 \times 10^{-16}$ |
| | GO | GO:0015747 | urate transport | $3.46 \times 10^{-14}$ |
| | GO | GO:0015711 | organic anion transport | $3.23 \times 10^{-4}$ |

Continued on next page

**Table 4 – continued from previous page**

| Topic | Analysis | ID | Description | p.adjust |
|---|---|---|---|---|
| | KEGG | hsa04512 | ECM-receptor interaction | $1.24 \times 10^{-2}$ |
| | KEGG | hsa04931 | Insulin resistance | $1.82 \times 10^{-2}$ |
| | KEGG | hsa03082 | ATP-dependent chromatin remodeling | $2.08 \times 10^{-2}$ |
| Topic 3 | GO | GO:0031341 | regulation of cell killing | $8.43 \times 10^{-6}$ |
| | GO | GO:0001906 | cell killing | $9.16 \times 10^{-6}$ |
| | GO | GO:0001910 | regulation of leukocyte mediated cytotoxicity | $4.83 \times 10^{-5}$ |
| | KEGG | hsa05330 | Allograft rejection | $1.06 \times 10^{-7}$ |
| | KEGG | hsa04940 | Type I diabetes mellitus | $1.97 \times 10^{-7}$ |
| | KEGG | hsa05332 | Graft-versus-host disease | $2.22 \times 10^{-7}$ |
| Topic 4 | GO | GO:0002396 | MHC protein complex assembly | $4.26 \times 10^{-16}$ |
| | GO | GO:0002501 | peptide antigen assembly with MHC protein complex | $4.26 \times 10^{-16}$ |
| | GO | GO:0002399 | MHC class II protein complex assembly | $1.14 \times 10^{-15}$ |
| | KEGG | hsa04940 | Type I diabetes mellitus | $1.38 \times 10^{-17}$ |
| | KEGG | hsa04612 | Antigen processing and presentation | $1.12 \times 10^{-14}$ |
| | KEGG | hsa05310 | Asthma | $1.52 \times 10^{-14}$ |
| Topic 5 | GO | GO:0006334 | nucleosome assembly | $6.04 \times 10^{-3}$ |
| | GO | GO:0034728 | nucleosome organization | $6.36 \times 10^{-3}$ |
| | GO | GO:0015747 | urate transport | $4.41 \times 10^{-2}$ |
| | KEGG | hsa05322 | Systemic lupus erythematosus | $2.62 \times 10^{-7}$ |
| | KEGG | hsa05034 | Alcoholism | $3.08 \times 10^{-5}$ |
| | KEGG | hsa04613 | Neutrophil extracellular trap formation | $3.40 \times 10^{-5}$ |
| Topic 6 | GO | GO:0031342 | negative regulation of cell killing | $9.07 \times 10^{-4}$ |
| | GO | GO:0097242 | amyloid-beta clearance | $9.07 \times 10^{-4}$ |
| | GO | GO:0002460 | adaptive immune response ... from immunoglobulin superfamily domains | $9.07 \times 10^{-4}$ |
| | KEGG | hsa04610 | Complement and coagulation cascades | $4.67 \times 10^{-5}$ |
| | KEGG | hsa04640 | Hematopoietic cell lineage | $1.57 \times 10^{-3}$ |
| | KEGG | hsa04979 | Cholesterol metabolism | $6.50 \times 10^{-3}$ |
| Topic 7 | GO | GO:0002396 | MHC protein complex assembly | $2.04 \times 10^{-13}$ |
| | GO | GO:0002501 | peptide antigen assembly with MHC protein complex | $2.04 \times 10^{-13}$ |
| | GO | GO:0048002 | antigen processing and presentation of peptide antigen | $2.93 \times 10^{-11}$ |
| | KEGG | hsa04612 | Antigen processing and presentation | $2.62 \times 10^{-14}$ |
| | KEGG | hsa05320 | Autoimmune thyroid disease | $2.77 \times 10^{-12}$ |
| | KEGG | hsa05330 | Allograft rejection | $1.79 \times 10^{-11}$ |
| Topic 8 | GO | GO:0048002 | antigen processing and presentation of peptide antigen | $1.05 \times 10^{-5}$ |
| | GO | GO:0002486 | antigen processing & presentation ... via ER pathway, TAP-independent | $1.05 \times 10^{-5}$ |

Continued on next page

**Table 4 – continued from previous page**

| Topic | Analysis | ID | Description | p.adjust |
|---|---|---|---|---|
| | GO | GO:0002476 | antigen processing and presentation of endogenous peptide antigen via MHC class Ib | $1.05 \times 10^{-5}$ |
| | KEGG | hsa05330 | Allograft rejection | $4.03 \times 10^{-12}$ |
| | KEGG | hsa04940 | Type I diabetes mellitus | $9.96 \times 10^{-12}$ |
| | KEGG | hsa05332 | Graft-versus-host disease | $1.18 \times 10^{-11}$ |
| Topic 9 | GO | GO:0032330 | regulation of chondrocyte differentiation | $1.66 \times 10^{-2}$ |
| | GO | GO:0061035 | regulation of cartilage development | $2.81 \times 10^{-2}$ |
| | GO | GO:0001657 | ureteric bud development | $2.81 \times 10^{-2}$ |
| | KEGG | hsa04540 | Gap junction | $1.45 \times 10^{-3}$ |
| | KEGG | hsa04350 | TGF-beta signaling pathway | $2.66 \times 10^{-3}$ |
| | KEGG | hsa04722 | Neurotrophin signaling pathway | $3.41 \times 10^{-3}$ |
| Topic 10 | GO | GO:0008217 | regulation of blood pressure | $4.66 \times 10^{-5}$ |
| | GO | GO:0007263 | nitric oxide mediated signal transduction | $3.01 \times 10^{-4}$ |
| | GO | GO:0038060 | nitric oxide-cGMP-mediated signaling | $6.55 \times 10^{-4}$ |
| | KEGG | hsa04924 | Renin secretion | $1.61 \times 10^{-6}$ |
| | KEGG | hsa04022 | cGMP-PKG signaling pathway | $7.70 \times 10^{-6}$ |
| | KEGG | hsa04261 | Adrenergic signaling in cardiomyocytes | $8.14 \times 10^{-5}$ |
| Topic 11 | GO | - | *No significant enrichment* | - |
| | KEGG | hsa00603 | Glycosphingolipid biosynthesis - globo and isoglobo series | $8.68 \times 10^{-4}$ |
| | KEGG | hsa00601 | Glycosphingolipid biosynthesis - lacto and neolacto series | $2.68 \times 10^{-3}$ |
| | KEGG | hsa04218 | Cellular senescence | $9.04 \times 10^{-3}$ |
| Topic 12 | GO | GO:0002396 | MHC protein complex assembly | $1.54 \times 10^{-2}$ |
| | GO | GO:0002501 | peptide antigen assembly with MHC protein complex | $1.54 \times 10^{-2}$ |
| | KEGG | hsa04940 | Type I diabetes mellitus | $2.07 \times 10^{-4}$ |
| | KEGG | hsa05323 | Rheumatoid arthritis | $1.98 \times 10^{-3}$ |
| | KEGG | hsa05310 | Asthma | $3.23 \times 10^{-3}$ |
| Topic 13 | GO | GO:0046879 | hormone secretion | $2.22 \times 10^{-8}$ |
| | GO | GO:0009914 | hormone transport | $2.22 \times 10^{-8}$ |
| | GO | GO:0030072 | peptide hormone secretion | $2.96 \times 10^{-7}$ |
| | KEGG | hsa04950 | Maturity onset diabetes of the young | $1.03 \times 10^{-8}$ |
| | KEGG | hsa04917 | Prolactin signaling pathway | $1.19 \times 10^{-3}$ |
| | KEGG | hsa04911 | Insulin secretion | $2.07 \times 10^{-3}$ |
| Topic 14 | GO | GO:0042438 | melanin biosynthetic process | $1.38 \times 10^{-4}$ |
| | GO | GO:0019748 | secondary metabolic process | $1.38 \times 10^{-4}$ |
| | GO | GO:0006582 | melanin metabolic process | $1.38 \times 10^{-4}$ |
| | KEGG | hsa04382 | Cornified envelope formation | $2.32 \times 10^{-4}$ |
| | KEGG | hsa04145 | Phagosome | $7.82 \times 10^{-3}$ |

Continued on next page

**Table 4 – continued from previous page**

| Topic | Analysis | ID | Description | p.adjust |
|---|---|---|---|---|
| | KEGG | hsa05164 | Influenza A | $1.06 \times 10^{-2}$ |
| Topic 15 | GO | GO:0010876 | lipid localization | $4.16 \times 10^{-7}$ |
| | GO | GO:0006869 | lipid transport | $1.65 \times 10^{-6}$ |
| | GO | GO:0042632 | cholesterol homeostasis | $7.43 \times 10^{-6}$ |
| | KEGG | hsa04979 | Cholesterol metabolism | $9.67 \times 10^{-10}$ |
| | KEGG | hsa04976 | Bile secretion | $2.50 \times 10^{-5}$ |
| | KEGG | hsa05231 | Choline metabolism in cancer | $5.93 \times 10^{-4}$ |
